# Rice blast fungus hijacks a rice RALF peptide to initiate infection

**DOI:** 10.64898/2026.09.30.754903

**Authors:** Fumi Fukada, Shihoko Fujii, Naomi Konishi, Tomoyuki Furuta, Yuta Hino, Alfino Sebastian, Motoki Shimizu, Yuta Harada, Takayuki Arazoe, Ivan Galis, Soichiro Asuke, Falk-Fooken Decker, Ting Guo, Natsuko Ono, Hideki Nishimura, Weilin Cao, Maike Kloos, Haruka Minato, Daisuke Suzuki, Pingyu Wang, Yoji Kawano

## Abstract

Plant-derived peptides regulate immunity and host–microbe interactions, but whether fungal pathogens exploit endogenous plant peptides for infection-site selection remains largely unknown. Rapid alkalinization factors (RALFs) are a family of plant peptide hormones that contribute to plant development, reproduction, stress responses, and cell-wall organization. Here, we show that OsRALF7, which is an immune-activating peptide in rice (*Oryza sativa*), is co-opted by the rice blast fungus *Magnaporthe oryzae* as a host-derived cue for appressorium formation, specialized infection structures required for host penetration. While synthetic OsRALF7 peptide induced defense responses in rice suspension cells, the same peptide induced appressorium formation in *M. oryzae*. OsRALF7-induced appressorium formation was associated with activation of the cAMP/PKA and Pmk1 MAPK pathways and required a hydrophobic surface. *M. oryzae* inoculated onto *Osralf7* knockout plants formed fewer appressoria, and the positioning of these appressoria relative to the boundaries between two adjacent plant epidermal cells was altered. These findings suggest that OsRALF7 provides positional information for infection initiation for *M. oryzae*. OsRALF7 localized to the plasma membrane and cell wall in rice, and its appressorium-inducing activity required a conserved YISY motif and cysteine residues. This activity was not conserved in *Colletotrichum higginsianum*, as RALF23 from *Arabidopsis thaliana* did not induce appressorium formation but instead showed antifungal activity. Together, these findings uncover a dual function of endogenous plant immune-modulating RALF peptides at the plant–pathogen interface, where they can be exploited by *M. oryzae* as a host-derived cue for infection initiation while also retaining antifungal activity against other fungal species.

## Introduction

Small, secreted peptides play central roles in the regulation of growth, development, and stress responses in plants, serving as important mediators of intercellular communication. In plant immunity, a subset of these peptides functions as signaling molecules that alert the plant to pathogen invasion and amplify defense responses. These immune-associated peptides, termed phytocytokines, are typically derived from larger precursor proteins and secreted into the extracellular space (1). Upon release, they are perceived by plasma-membrane receptors and activate defense signaling in autocrine and/or paracrine manners. In addition, since many plant peptides accumulate in the apoplast, a major site of initial host-pathogen contact, some also act as antimicrobial peptides that directly inhibit or kill invading pathogens (2). Despite these important functions, the roles of many plant peptides still remain poorly understood, particularly their direct effects on microbes.

Among plant peptide families, the Rapid Alkalinization Factor (RALF) family is widely conserved across land plants and represents a prominent class of extracellular signaling peptides. RALF-type peptides are synthesized as precursor proteins with four characteristic motifs: an N-terminal secretion signal sequence; a site-1 protease (S1P)-type proteolytic cleavage site, typically with an RRXL motif; a receptor-binding motif containing the YISY motif; and four conserved cysteine residues that form two intramolecular disulfide bonds (3, 4). Mature RALFs are generated by proteolytic processing of their cognate propeptides. RALFs function through receptor complexes composed of *Catharanthus roseus* receptor-like kinase 1-like (CrRLK1L) proteins and glycosylphosphatidylinositol (GPI)-anchored proteins of the LORELEI (LRE)/LORELEI-like GPI-anchored protein (LLG) family. Especially, FERONIA (FER) and LLG1 play a predominant role in RALF perception. This complex regulates cell expansion, extracellular pH, cell-wall integrity, and immune signaling (5–8). In addition to their signaling roles, recent studies have shown that some RALFs contribute to cell-wall organization through interactions with demethylesterified pectin and leucine-rich repeat extensins (LRXs) (9–12). Together, these findings indicate that RALFs function as extracellular signaling peptides with additional roles in cell-wall organization.

AtRALF23 is one of the best-characterized RALFs in the context of immune regulation. Upon pathogen infection in *Arabidopsis* (*Arabidopsis thaliana*), the AtRALF23 propeptide is rapidly processed by S1P protease and released into the apoplast, where the mature peptide binds to the FER–LLG1 receptor complex (6, 13, 14). This inhibits the formation of membrane-bound pattern recognition receptor (PRR) complexes and thereby suppresses PRR-triggered immunity (PTI) signaling (6). Consistent with this, the *atralf23* knockout mutant exhibits increased resistance to the bacterial pathogen *Pseudomonas syringae* pv. *tomato* DC3000. Recently, CRISPR/Cas9-generated multiple mutants lacking *ralf22 ralf23 ralf33* and *ralf1 ralf18 ralf22 ralf23 ralf31 ralf33 ralf34* showed enhanced resistance to powdery mildew, an obligate biotrophic fungus, with resistance levels correlating with the number of disrupted RALF genes (15). This powdery mildew susceptibility likely depends in part on FER–RALF-mediated regulation of apoplastic pH and cell-wall remodeling, although this process also appears to be partly FER-independent. Thus, the effects of RALF peptides on plant–pathogen interactions depend on the specific RALFs, their cognate receptors, and invading pathogens.

*Magnaporthe oryzae* is a globally important fungal pathogen of rice (*Oryza sativa*) and the causal agent of rice blast disease, which causes severe yield losses worldwide. Its pathogenicity depends primarily on the formation of a specialized infection structure called an appressorium, which enables host penetration (16). The infection process of this fungus is initiated by the attachment of a conidium to the leaf surface. Upon germination, the conidium produces a polarized germ tube, which forms a dome-shaped, single-celled appressorium at its tip. The mature appressorium generates enormous turgor pressure of up to 8.0 MPa through the accumulation of compatible solutes such as glycerol (17). This pressure is transmitted through the penetration peg and, together with enzymatic degradation of the plant surface, enables the fungus to breach the cuticle and cell wall. Appressorium formation is induced by physical cues, such as surface hardness and hydrophobicity, as well as chemical signals, including leaf waxes and cutin monomers (18). These cues are sensed by fungal receptors such as the G-protein-coupled receptor (GPCR) Pth11 and the sensor proteins Msb2 and Sho1, which activate downstream cyclic AMP (cAMP) and mitogen-activated protein kinase (MAPK) signaling pathways to trigger appressorium development (18, 19).

Given that the known appressorium-inducing factors are generally ubiquitous on the plant surface, fungi may sense additional, as yet unidentified, host-derived signals to determine the precise infection site within the short time before host recognition and immune responses are activated. In several phylloplane fungal pathogens, such as *Colletotrichum* species, *Bipolaris maydis*, *Ustilago maydis*, and *Botrytis cinerea*, appressorium formation and subsequent penetration preferentially occur at anticlinal cell junctions, the boundaries between adjacent epidermal cells (20–25). For example, in *Alternaria cirsinoxia*, more than 75% of appressoria form at anticlinal junctions of epidermal cells rather than on periclinal walls or stomata, and subsequent penetration occurs at these junctions (26). Thus, these findings suggest that anticlinal junctions may provide a favorable route of entry for foliar fungal pathogens. However, the signals that guide fungi to the anticlinal regions are not fully understood.

We previously identified *OsRALF7* (LOC_Os01g25560) as a candidate phytocytokine involved in rice immunity following inoculation with *M. oryzae* and chitin treatment (27). Although RT-qPCR confirmed the chitin-dependent induction of *OsRALF7*, the function of its encoded peptide remains unclear. Here, we show that OsRALF7 not only induces immune responses in rice, but is also exploited by *M. oryzae* as a host-derived cue that promotes appressorium formation. Application of synthetic OsRALF7 enhanced appressorium formation of *M. oryzae*. This response was associated with activation of cAMP/PKA and Pmk1 MAPK pathways and required a hydrophobic surface. On *Osralf7* mutant plants*, M. oryzae* formed fewer appressoria, and the positioning of appressoria relative to the anticlinal boundaries between two adjacent plant epidermal cells was altered. We further show that *Arabidopsis* RALF23 did not induce appressorium formation in *Colletotrichum higginsianum* but instead exhibited antifungal activity, suggesting that RALFs can have diverse, context-dependent effects on pathogenic fungi. Together, these findings suggest that *M. oryzae* exploits plant endogenous RALFs as potential positional cues for infection initiation, whereas RALFs can also exhibit antifungal activity against *C. higginsianum*.

## Results

### Identification of OsRALF7 as a phytocytokine that elicits immune responses in rice

We previously performed an integrated transcriptomic and proteomic analysis to identify small, secreted peptides that modulate rice immune responses (27). From this analysis, we identified OsRALF7 as one of the most abundant proteins in the culture medium of rice suspension cells treated with the fungal elicitor chitin (Fig. 1*A*). OsRALF7 contains the four canonical features of RALF family members: an N-terminal signal sequence for secretion, a protease recognition motif (RRXL), a putative receptor-binding motif (YISY), and four conserved cysteine residues (Fig. 1*B*) (28). Previous work showed that *OsRALF7* expression is induced in rice suspension cells by chitin and the bacterial elicitor peptidoglycan, suggesting a role for this gene in rice immunity (27). Together, these features prompted us to ask whether OsRALF7 functions as an immune-active peptide in rice.

**Figure 1.**
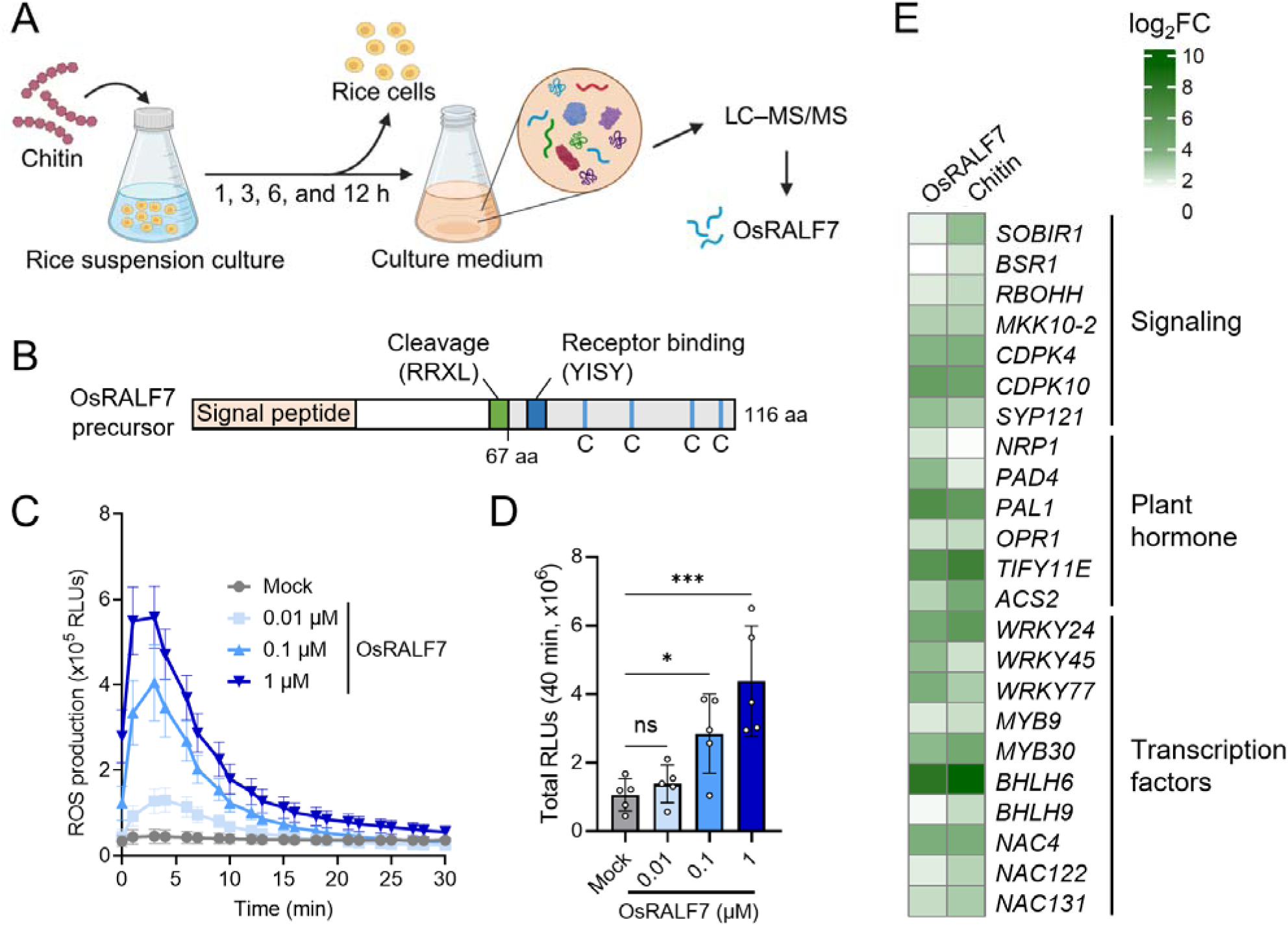
OsRALF7 elicits rice immunity. (A) Diagram illustrating the pipeline followed for the identification of OsRALF7. Multiple peptides derived from OsRALF7 were detected by secretome proteomics of culture filtrates from rice suspension cell cultures treated with 2 μg/mL chitin. (B) Diagram showing the functional domains of OsRALF7. Proteolytic processing of the OsRALF7 precursor is predicted to occur at a cleavage site containing the dibasic RRXL motif, generating a mature peptide that contains a putative receptor-binding motif (YISY) and four cysteine residues. (C) Time course of the ROS burst after treatment with OsRALF7 in rice suspension cells. Wild-type Nipponbare cell cultures were treated with the indicated concentrations of synthetic OsRALF7, and luminescence was recorded every 1.5 min over 30 min. Data are presented as the mean ± SEM from five independently cultured flasks processed in parallel (*n* = 5). Experiments were performed three times, each time with similar results. (D) ROS production as a function of OsRALF7 dosage. Total ROS production over 40 min, expressed as relative light units (RLUs). Each circle represents one independently cultured flask (*n* = 5). Error bars indicate SD. Statistical significance of differences was assessed by one-way ANOVA followed by Dunnett’s multiple-comparisons test. ***, *P* < 0.001; *, *P* < 0.05; ns, not significant. (E) Heatmap representation of transcript levels for the indicated upregulated genes by both OsRALF7 and chitin in rice cell cultures. Color scale indicates log_2_ fold-change values relative to mock treatment.

Given the reported immunomodulatory roles of RALF family members in other species, we examined whether OsRALF7 can trigger immune responses in rice. For these experiments, we synthesized a 49-amino acid (aa) peptide corresponding to the mature OsRALF7 (aa 68–116). When cell suspension cultures derived from wild-type rice plants (cv. Nipponbare) were treated with synthetic OsRALF7, the cells exhibited a rapid, dose-dependent ROS burst in a luminol-based assay (Fig. 1*C* and *D*). Treatment with chitin also triggered a ROS response in the suspension culture cells. Co application of OsRALF7 and chitin induced a ROS response, indicating an additive effect of the two treatments on early immune signaling (*SI Appendix*, Fig. S1). By contrast, a mutant peptide lacking the putative receptor-binding motif (OsRALF7^ΔYISY^) failed to elicit ROS production, suggesting that the YISY motif is required for OsRALF7 phytocytokine activity (*SI Appendix*, Fig. S1). This result supports previous studies showing that RALF perception is receptor dependent in other plant species (8, 29, 30).

To assess whether OsRALF7 causes transcriptome reprogramming, we performed RNA-seq on rice suspension cells treated with water only as mock control or OsRALF7 for 1 h (*SI Appendix*, Table S1). OsRALF7 treatment resulted in 2,768 differentially expressed genes (DEGs), comprising 2,462 upregulated and 306 downregulated genes. Gene Ontology (GO) term enrichment analysis showed that OsRALF7-upregulated DEGs were highly enriched in terms associated with biotic stress and defense responses, whereas OsRALF7-downregulated DEGs were associated with development (*SI Appendix*, Fig. S2).

To determine the extent to which the OsRALF7-responsive immune transcriptome overlaps with chitin-induced PTI, we conducted an additional RNA-seq on rice suspension cells treated with chitin under the same conditions and compared the two DEG datasets. Following chitin treatment, we identified 2,340 DEGs, of which 2,207 were upregulated and 133 were downregulated. Notably, 56% of the OsRALF7-responsive DEGs overlapped with chitin-induced DEGs (*SI Appendix*, Fig. S3), indicating a substantial convergence between OsRALF7-triggered and chitin-triggered transcriptomic programs. The overlapping genes included those associated with immune signaling, plant hormone pathways, and immunity-related transcription factors (Fig. 1*E*). Collectively, these biochemical and transcriptomic data indicate that OsRALF7 functions as a peptide signal that activates immunity-associated gene expression in rice.

### Mature OsRALF7 induces appressorium formation in *M. oryzae in vitro*

Since OsRALF7 contains a predicted signal peptide and is therefore likely secreted into the extracellular space, OsRALF7 may encounter *M. oryzae* at the plant–pathogen interface during infection. In addition, RALFs have previously been classified as defensin-like peptides (31, 32), and plant defensins are cysteine-rich antimicrobial peptides that often directly interact with microbial surface or membrane components (2). We therefore examined whether application of synthetic mature OsRALF7 could influence infection-related morphogenesis in *M. oryzae*. Accordingly, we incubated *M. oryzae* conidia on coverslips in liquid complete medium (CM), which permits vegetative hyphal growth but not infection-related morphogenesis (33). In the mock control with water only, conidia germinated and formed only vegetative hyphae (Fig. 2*A*). By contrast, treatment with mature OsRALF7 induced formation of appressoria. This appressorium-inducing activity was dose dependent, with 2 µM OsRALF7 representing the minimum effective concentration tested (Fig. 2*B*). This effect was markedly weaker with OsRALF7^ΔYISY^, indicating that the YISY motif is needed for the appressorium-inducing activity of OsRALF7. Time course observations further characterized the morphogenetic transitions induced by OsRALF7 treatment. In mock treated samples, conidial germ tubes elongated continuously at all observed time points (Fig. 2*C* and *D*). By contrast, OsRALF7 treatment triggered the formation of swollen, hook-like structures at the tip of germ tubes by 2 h after onset of treatment, followed by non-melanized appressoria at 4 h, and fully melanized mature appressoria at 6 h. These events were rarely observed with the OsRALF7^ΔYISY^-treated samples.

**Figure 2.**
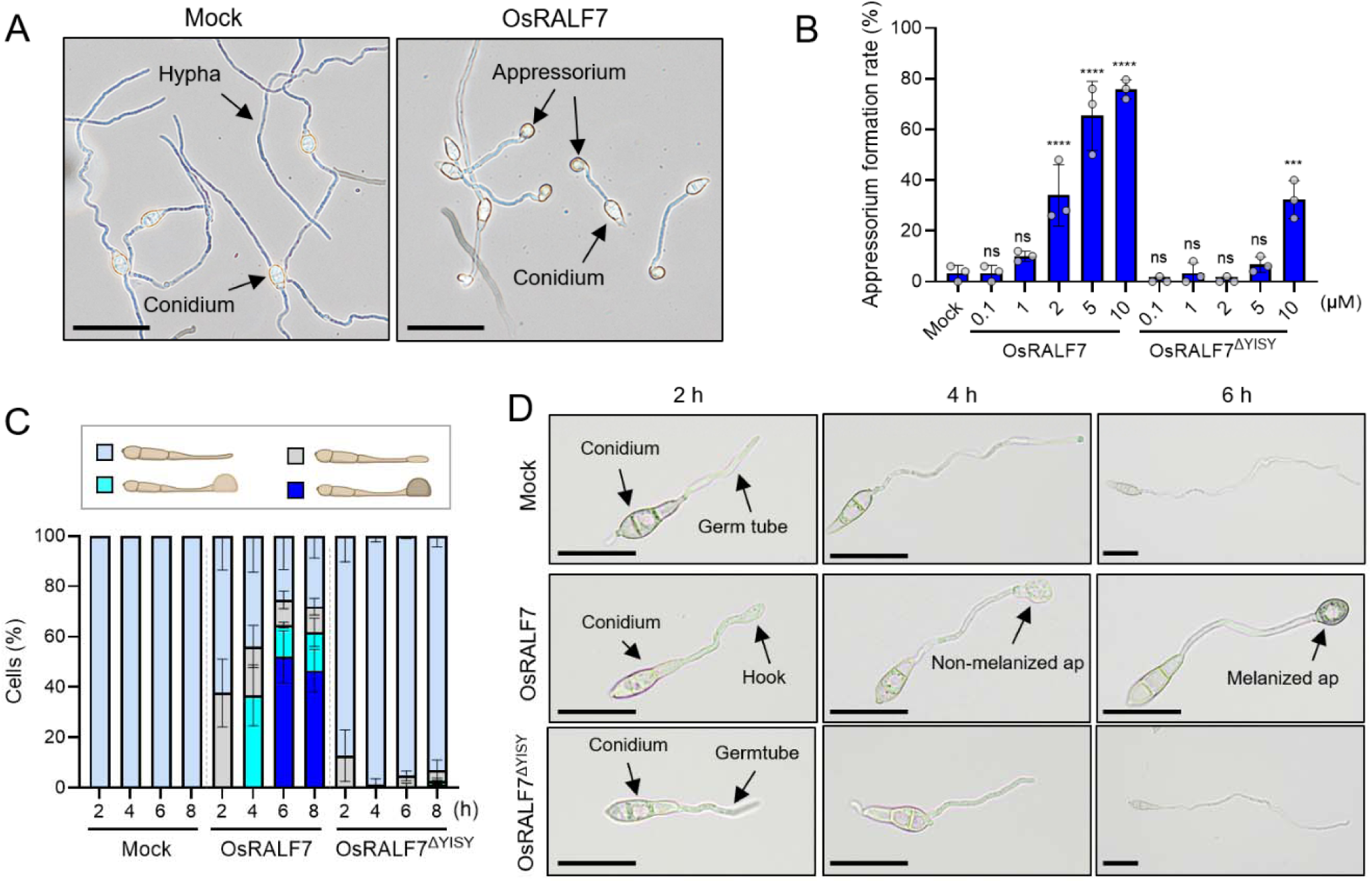
OsRALF7 promotes appressorium formation in *M. oryzae in vitro*. (A) Representative micrographs of *M. oryzae* treated with water only (Mock) or OsRALF7 (5 μM) for 8 h. Arrowheads indicate appressoria. Scale bars, 50 µm. (B) Dose-dependent appressorium formation of *M. oryzae* upon OsRALF7 treatment. Appressorium formation rates were calculated after treatment with the indicated concentrations of OsRALF7 or OsRALF7 lacking the YISY motif (OsRALF7^ΔYISY^) in 50% CM medium for 8 h. Each circle represents one independently cultured plate (*n* = 3), with >50 conidia scored per plate. Statistical significance of differences was assessed by one-way ANOVA followed by Dunnett’s multiple-comparisons test relative to Mock. ****, *P* < 0.0001; ***, *P* < 0.001; ns, not significant. Error bars indicate SD. Experiments were performed three times, each time with similar results. (C) Time-course analysis of appressorium formation following OsRALF7 treatment. Conidia were incubated on 50% CM medium to which water only (Mock), 5 µM OsRALF7, or 5 µM OsRALF7^ΔYISY^ was added. Light blue indicates germ tube elongation only, gray indicates hook formation, bright light blue indicates non-melanized appressorium formation, and dark blue indicates melanized appressorium formation. For each treatment and time point, 50 conidia were scored per independently cultured plate (*n* = 3), and the percentage of appressorium formation was calculated. Error bars indicate SD. Experiments were performed three times, each time with similar results. *(D)* Representative time-course micrographs. Ap, appressorium. Scale bars, 20 µm.

### OsRALF7-induced appressorium formation involves the cAMP/PKA and MAPK pathways

To investigate the mechanism by which OsRALF7 promotes appressorium formation, we examined whether OsRALF7 activates the cAMP/PKA and Pmk1 MAPK pathways in *M. oryzae* (Fig. 3*A*), both of which play key roles in appressorium formation (34). Conidia were treated with synthetic OsRALF7 (5 µM) or water as a mock control, and samples were collected 4 h after treatment, a time point at which OsRALF7-treated cells had formed non-melanized appressoria (Fig. 2*C* and *D*). Intracellular cAMP levels and Pmk1 phosphorylation were then examined. Intracellular cAMP levels were significantly higher in OsRALF7-treated conidia than in mock-treated controls (Fig. 3*B*), and Pmk1 phosphorylation was also increased by OsRALF7 treatment (Fig. 3*C*). Neither response was detected when vegetative mycelia were treated with OsRALF7 instead of conidia (*SI Appendix*, Fig. S4), suggesting that OsRALF7-induced activation of these pathways is specific to germinated conidia. Consistent with these responses, RNA-seq analysis of OsRALF7-treated conidia revealed dose-dependent induction of genes associated with appressorium maturation, melanin biosynthesis, and host penetration (*SI Appendix*, Fig. S5) (35–39). Many of the genes induced by OsRALF7 overlapped with genes down-regulated in a previously reported Δ*pmk1* mutant and with genes induced by cAMP treatment (37, 40, 41). Together, these results suggest that OsRALF7 promotes appressorium formation by activating the cAMP/PKA and Pmk1 MAPK pathways and their downstream transcriptional programs in *M. oryzae*.

**Figure 3.**
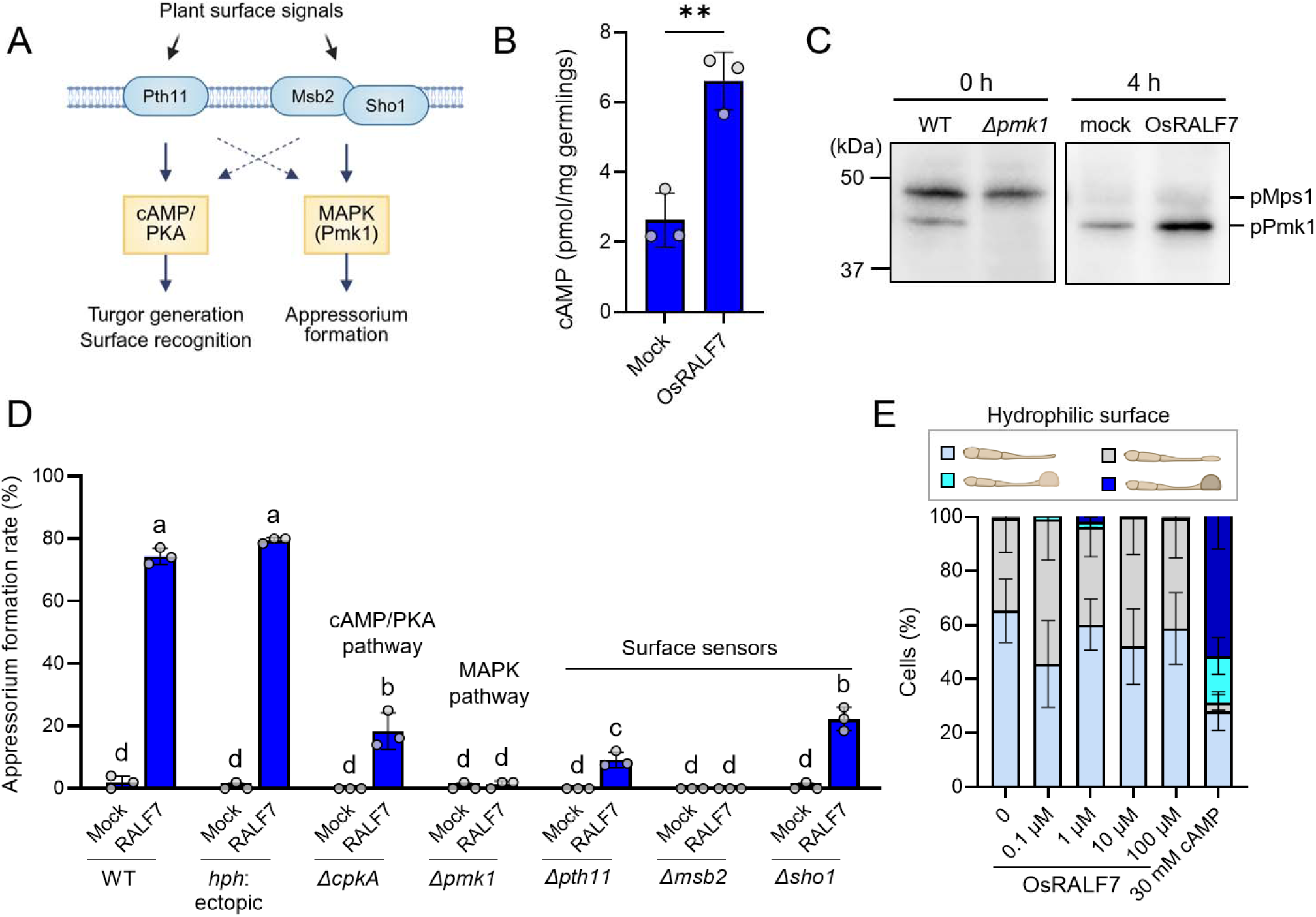
OsRALF7-induced appressorium formation is associated with the cAMP/PKA and MAPK signaling. (A) Schematic model of the reported signaling pathways regulating appressorium formation in *M. oryzae*. (B) Intracellular cAMP levels in germinated conidia. *M. oryzae* conidia were incubated in 25% CM medium containing water (mock) or OsRALF7 (5 μM) for 4 h, followed by cAMP measurement. Each circle represents an independent experiment (*n* = 3). Error bars indicate SD. Statistical significance of differences was assessed by an unpaired two-tailed *t*-test. **, *P* < 0.01. (C) Immunoblot analysis of phosphorylated Pmk1. Wild-type (WT) and Δ*pmk1* conidia were collected from plates without incubation, or WT conidia were incubated in 25% CM medium containing water (mock) or OsRALF7 (5 μM) for 4 h, followed by total protein extraction. Phosphorylated Pmk1 was detected using an anti-phospho-p44/42 MAPK antibody, which has also been reported to detect phosphorylated Mps1. Experiments were performed three times, each time with similar results. (D) Appressorium formation in appressorium-defective *M. oryzae* mutants. Conidia were incubated in 25% CM medium containing water (mock) or OsRALF7 (5 μM) for 8 h. Each circle represents one independently cultured plate (*n* = 3), with >50 conidia scored per plate. Statistical significance of differences was assessed by one-way ANOVA followed by Tukey’s multiple-comparison test. Different letters indicate significant differences at *P* < 0.05. Experiments were performed three times, each time with similar results. (E) Appressorium formation on a hydrophilic GelBond surface following OsRALF7 treatment. Conidia were incubated in water for 24 h with the indicated concentrations of OsRALF7. cAMP (30 mM) served as a positive control. Light blue indicates germ tube elongation only, gray indicates hook formation, bright light blue indicates non-melanized appressorium formation, and dark blue indicates melanized appressorium formation. For each treatment and time point, 50 conidia were scored per independently cultured plate (*n* = 3), and the percentage of appressorium formation was calculated. Error bars indicate SD. Experiments were performed three times, each time with similar results.

To genetically assess the contribution of the cAMP/PKA and Pmk1 MAPK pathways to OsRALF7-induced signaling, we generated Δ*cpkA* and Δ*pmk1* mutants in the *M. oryzae* P2 strain background (*SI Appendix*, Table S2). *CPKA* encodes the catalytic subunit of cAMP-dependent protein kinase A, whereas *PMK1* encodes the MAPK Pmk1. We first confirmed that the Δ*cpkA* mutant accumulated higher levels of intracellular cAMP than the wild type, consistent with previous reports (*SI Appendix*, Fig. S4*A*) (42). Immunoblotting also confirmed the absence of the phosphorylated Pmk1 in the Δ*pmk1* mutant (Fig. 3*C*; *SI Appendix*, Fig. S4*B*). We next examined appressorium formation in response to OsRALF7. The Δ*cpkA* mutant showed reduced OsRALF7-induced appressorium formation compared with the wild type, and the Δ*pmk1* mutant completely failed to form appressoria (Fig. 3*D*). Together with the OsRALF7-induced increases in intracellular cAMP and Pmk1 phosphorylation, these genetic data are consistent with the involvement of both the cAMP/PKA and Pmk1 MAPK pathways in the OsRALF7 response. However, given the general defect in appressorium formation of the Δ*pmk1* mutant, as previously reported (34), and observed in our study (*SI Appendix*, Fig. S6), we cannot exclude the possibility that the observed phenotype in the Δ*pmk1* mutant reflects a general defect in appressorium development rather than a specific requirement for Pmk1 in the response to OsRALF7.

### OsRALF7-induced appressorium formation depends on a hydrophobic surface

We next examined whether the known surface sensors Pth11, Msb2, and Sho1 contribute to OsRALF7-induced appressorium formation. Pth11 is a GPCR that perceives hydrophobic surface cues and primarily activates cAMP signaling (19, 43), whereas Msb2 and Sho1 are transmembrane sensors implicated in the perception of surface hydrophobicity, cutin monomers, and waxes and primarily activate the Pmk1 MAPK pathway (Fig. 3*A*) (18). We generated Δ*pth11*, Δ*msb2*, and Δ*sho1* mutants in the P2 background, all of which failed to form appressoria when incubated in water on the hydrophobic surface (*SI Appendix*, Fig. S7; Table S2). These defects in appressorium formation were consistent with previous reports, although the mutants analyzed here exhibited more severe phenotypes than those reported in other strain backgrounds (18, 19, 43). Nevertheless, when treated with synthetic OsRALF7 in CM medium, the Δ*pth11* and Δ*sho1* mutants showed lower appressorium formation rates than the wild type, but retained partial responsiveness to OsRALF7 (Fig. 3*D*, *SI Appendix*, Fig. S6). By contrast, the Δ*msb2* mutant completely failed to form appressoria even after OsRALF7 treatment. However, since this mutant was severely impaired in appressorium formation under multiple conditions tested, we could not determine whether its phenotype reflected a specific requirement for Msb2 in the OsRALF7 response or a general defect in appressorium development. Together, these results suggest that at least Pth11 and Sho1, known upstream components of the cAMP/PKA and Pmk1 MAPK pathways, contribute to efficient OsRALF7-induced appressorium formation. However, loss of either protein alone does not completely abolish responsiveness to OsRALF7.

Given the involvement of Pth11, Msb2, and Sho1 in hydrophobic-surface sensing (18, 19), we next examined whether OsRALF7-induced appressorium formation depends on a hydrophobic surface. To this end, *M. oryzae* conidia were incubated with OsRALF7 on a hydrophilic GelBond surface, and appressorium formation was assessed. Although exogenous cAMP induced appressorium formation, as previously reported (44), OsRALF7 failed to induce appressorium formation on the hydrophilic surface at all concentration tested (Fig. 3*E*). These results indicate that OsRALF7-induced appressorium formation requires a hydrophobic-surface input and OsRALF7 alone is insufficient to substitute for this physical cue.

To determine whether OsRALF7 further promotes appressorium formation in the absence of nutrients, we assessed appressorium formation in water on a hydrophobic surface. OsRALF7 treatment produced a modest but nonsignificant increase in appressorium formation (*SI Appendix*, Fig. S7). The limited effect may be attributable to the high basal level of appressorium formation in mock-treated controls under these conditions.

### Loss of OsRALF7 decreases appressorium formation and alters appressorium positioning

To examine whether endogenous plant-derived OsRALF7, in addition to synthetic OsRALF7, contributes to appressorium formation in *M. oryzae*, we generated knockout lines for *OsRALF7* (*Osralf7*) using clustered regularly interspaced short palindromic repeats (CRISPR)/CRISPR-associated nuclease 9 (Cas9)-mediated editing (*SI Appendix*, Fig. S8). We inoculated *M. oryzae* onto leaf sheaths of wild-type and *Osralf7* mutant rice plants and quantified appressorium formation at 6 h post-inoculation. Consistent with the *in vitro* effects of synthetic mature OsRALF7, the rate of appressorium formation was significantly lower on the leaf sheath of the *Osralf7* mutants compared to those of the wild type (Fig. 4*A* and *B*). We also observed differences in the spatial distribution of appressoria on the leaf sheath surface. We therefore classified appressoria according to their contact with epidermal cell boundaries. In images acquired perpendicular to the leaf surface, appressoria were classified as “contacting anticlinal cell boundaries” when any part of their perimeter contacted a visible boundary between adjacent epidermal cells. Appressoria whose entire perimeter remained within the outer surface of a single epidermal cell were classified as “confined to periclinal cell surfaces” (Fig. 4*C*). On wild-type leaf sheaths, approximately 90% of all appressoria were contacted anticlinal cell boundaries (Fig. 4*A* and *D*). By contrast, this proportion decreased to approximately 70% on *Osralf7* mutant leaf sheaths, with a corresponding increase in appressoria confined to periclinal cell surfaces. To test whether appressoria in wild-type plants were randomly positioned or preferentially contacted anticlinal cell boundaries, we estimated the expected probability of an appressorium contacting an anticlinal cell boundary under a random positioning scenario. We calculated this probability using the equation 1−(x−2z)(y−2z)/xy, where x and y represent the longitudinal and transverse dimensions of the plant cell, respectively, and z represents the appressorium radius (Fig. 4*C*). Using the same random positioning model, the expected probabilities of appressorium contact with anticlinal cell boundaries were 69% in wild-type leaf sheaths, and 64% and 61% in the two *Osralf7* mutant lines (Fig. 4*D*). The observed frequency exceeded the random expectation by more than 20 percentage points in the wild type, but by only approximately 10 percentage points in the *Osralf7* mutants. These results suggest that *M. oryzae* preferentially forms appressoria that contact anticlinal cell boundaries in wild-type plants, whereas loss of *OsRALF7* reduces this preference.

**Figure 4.**
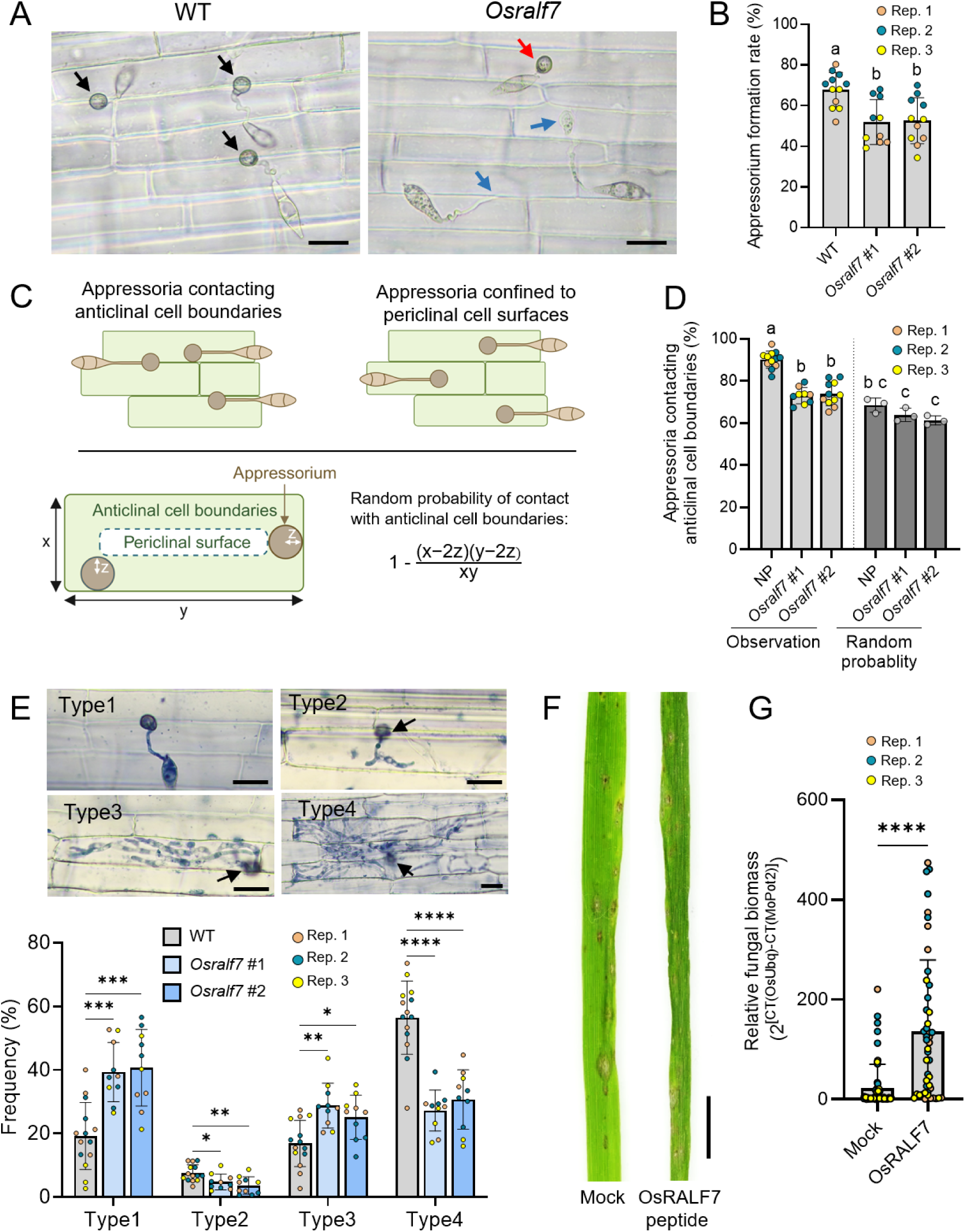
Loss of OsRALF7 function compromises appressorium formation and alters appressorium positioning. (A) Representative micrographs of *M. oryzae* on leaf sheaths of wild-type and *Osralf7* knockout rice plants at 6 h post inoculation (hpi). Black arrows, appressoria contacting anticlinal cell boundaries; red arrow, an appressorium confined to periclinal cell surfaces; blue arrows, continued germ tube elongation without appressorium formation or immature appressoria. Scale bars, 20 μm. (B) Appressorium formation frequency on rice leaf sheaths in wild-type and the *Osralf7* knockout. Each circle represents one independent rice plant. For each genotype, three or four individual plants were analyzed, and 150 conidia were scored per plant. Error bars indicate SD. Statistical significance of differences was assessed by one-way ANOVA followed by Tukey’s multiple-comparison test. Different letters indicate significant differences at *P* < 0.05. Data are from three independent experiments. (C) Diagram of appressorium positioning of *M. oryzae* on rice leaf sheath cells. The probability of contact with an anticlinal cell boundaries was calculated as 1−(x−2z)(y−2z)/xy, where x and y represent the longitudinal and transverse dimensions of the plant cell, respectively, and z represents appressorium radius. (D) Quantification of the proportion of appressoria contacting anticlinal cell boundaries of rice leaf sheath cells. Each circle represents one independent rice plant. For each genotype, three or four individual plants were analyzed, and 150 conidia were scored per plant. To assess the random probability of contact with anticlinal cell boundaries, three individual plants were analyzed for each genotype. For each plant, 50 plant cells and associated appressoria were counted, and the expected probability was calculated using the formula shown in (*C*). Error bars indicate SD. Statistical significance of differences was assessed by one-way ANOVA followed by Tukey’s multiple-comparison test. Different letters indicate significant differences at *P* < 0.05. Observation data are from three independent experiments. (E) Frequency of invasive hyphal development on leaf sheaths of wild-type and *Osralf7* knockout rice plants at 48 hpi. Infection sites were classified into four types: type 1, appressoria that failed to penetrate host cells; and types 2–4, successfully penetrated sites with short invasive hyphae (type 2), invasive hyphae expanding within a single host cell (type 3), or invasive hyphae extending into multiple host cells (type 4). Arrows indicate penetration sites. Invasive hyphae were visualized by aniline blue staining. Scale bars, 20 μm. Each circle represents one independent rice plant. For each genotype, three plants were analyzed, with 150 conidia scored per plant. Statistical significance was assessed by one-way ANOVA followed by Dunnett’s multiple-comparisons test relative to WT. \*\**P* < 0.01; \**P* < 0.05; ns, not significant. Error bars indicate SD. Experiments were performed three times, each time with similar results. (F) Spray-inoculation assay with synthetic OsRALF7. The highly blast-susceptible rice cultivar Moukoto was spray-inoculated with an *M. oryzae* conidial suspension containing OsRALF7 (5 μM). Representative images were acquired at 5 dpi. Scale bar, 5 mm. (G) Fungal biomass was quantified by qPCR after DNA extraction and is shown as ^[CT(OsUbq)−CT(MoPot2)]^, representing fungal DNA relative to plant DNA. Each dot represents one rice plant. Statistical significance of differences was assessed by an unpaired two-tailed *t*-test. ****, *P* < 0.0001. Data are from three independent experiments.

To determine whether altered leaf-surface properties could account for the reduced appressorium formation and altered appressorium positioning observed on *Osralf7* mutant plants, we compared leaf-surface characteristics between wild-type plants and two *Osralf7* mutant lines. Scanning electron microscopy revealed no apparent differences in epidermal surface topography between the genotypes (*SI Appendix*, Fig. S9*A* and *B*). The water contact angle, an indicator of leaf-surface hydrophobicity, did not differ significantly between wild-type and *Osralf7* mutants (*SI Appendix*, Fig. S9*C*). Finally, gas chromatography–mass spectrometry (GC–MS) analysis detected no significant differences in the major cuticular wax components, including long-chain fatty acids, alkanes, aldehydes, and primary alcohols (*SI Appendix*, Fig. S10). Together, these results suggest that the phenotypes observed on *Osralf7* mutant plants are unlikely to result from major alterations in leaf-surface topography, hydrophobicity, or cuticular wax composition.

To assess whether preferential appressorium contact with anticlinal cell boundaries was conserved across different rice–*M. oryzae* combinations, we analyzed three rice cultivars, including japonica and indica types, inoculated with four *M. oryzae* strains isolated from different geographic origins (*SI Appendix*, Fig. S11; Table S2). Across all combinations examined, the observed proportion of appressoria contacting anticlinal cell boundaries significantly exceeded the probability expected under random positioning. Thus, preferential appressorium contact with anticlinal cell boundaries was reproducible across the rice–*M. oryzae* combinations examined, although the magnitude of this preference may differ among fungal strain–rice cultivar combinations.

### OsRALF7 affects early fungal invasion and disease development

To determine whether the altered appressorium formation and positioning on *Osralf7* mutant plants influence subsequent fungal invasion, we quantified invasive hyphal development at 48 h post-inoculation. Infection sites were classified into four types: type 1, appressoria that failed to penetrate host cells; types 2, successfully penetrated sites with short invasive hyphae; type 3, invasive hyphae expanding within a single host cell; and type 4, invasive hyphae extending into multiple host cells (Fig. 4*E*). Compared with wild-type plants, *Osralf7* mutant plants showed a lower proportion of type 4 infection sites and higher proportion of type 1 appressoria. These findings suggest that the reduced appressorium formation on *Osralf7* mutant plants likely contributed to reduced subsequent host-cell penetration and impaired invasive hyphal development. At a later time point, however, lesion development and fungal biomass did not differ significantly between wild-type and *Osralf7* mutant plants (*SI Appendix*, Fig. S12), indicating that the early difference in invasive hyphal development was not maintained at the later disease-development stage examined.

To test whether exogenous OsRALF7 treatment could promote fungal infection *in planta*, we added synthetic OsRALF7 (5 µM) to *M. oryzae* conidial suspension and sprayed the suspension onto the highly blast-susceptible rice cultivar Moukoto. OsRALF7 treatment resulted in more severe lesions and significantly increased fungal biomass compared with the mock treatment, indicating that OsRALF7 treatment enhances fungal virulence *in planta* (Fig. 4*F* and *G*).

### Subcellular localization of OsRALF7-eGFP

Based on the above results, we hypothesized that *M. oryzae* promotes appressorium formation in response to rice-derived OsRALF7 present at the interface between rice cells and *M. oryzae*. We therefore assessed where OsRALF7 localizes in rice cells during appressorium development. To address this, we generated transgenic rice lines carrying *OsRALF7-eGFP*, which encodes a fusion of the OsRALF7 precursor with enhanced green fluorescent protein under the control of the native *OsRALF7* promoter (*proOsRALF7:OsRALF7-eGFP*). The transgenic plants showed no obvious growth defects, based on overall appearance and plant height (*SI Appendix*, Fig. S13). Immunoblotting analysis with an anti-GFP antibody detected a prominent band corresponding to the predicted size for the OsRALF7 propeptide, as well as bands consistent with cleaved mature OsRALF7-eGFP and free GFP (Fig. 5*A*). The relative abundance of these bands was 9.0:2.5:1.0 when that of free GFP was set to 1, indicating that the propeptide is the predominant detectable form, whereas a smaller fraction appears to be proteolytically processed into the mature form. This processing pattern is consistent with previous reports on AtRALF22 and AtRALF23 in *Arabidopsis* (13, 45, 14).

**Figure 5.**
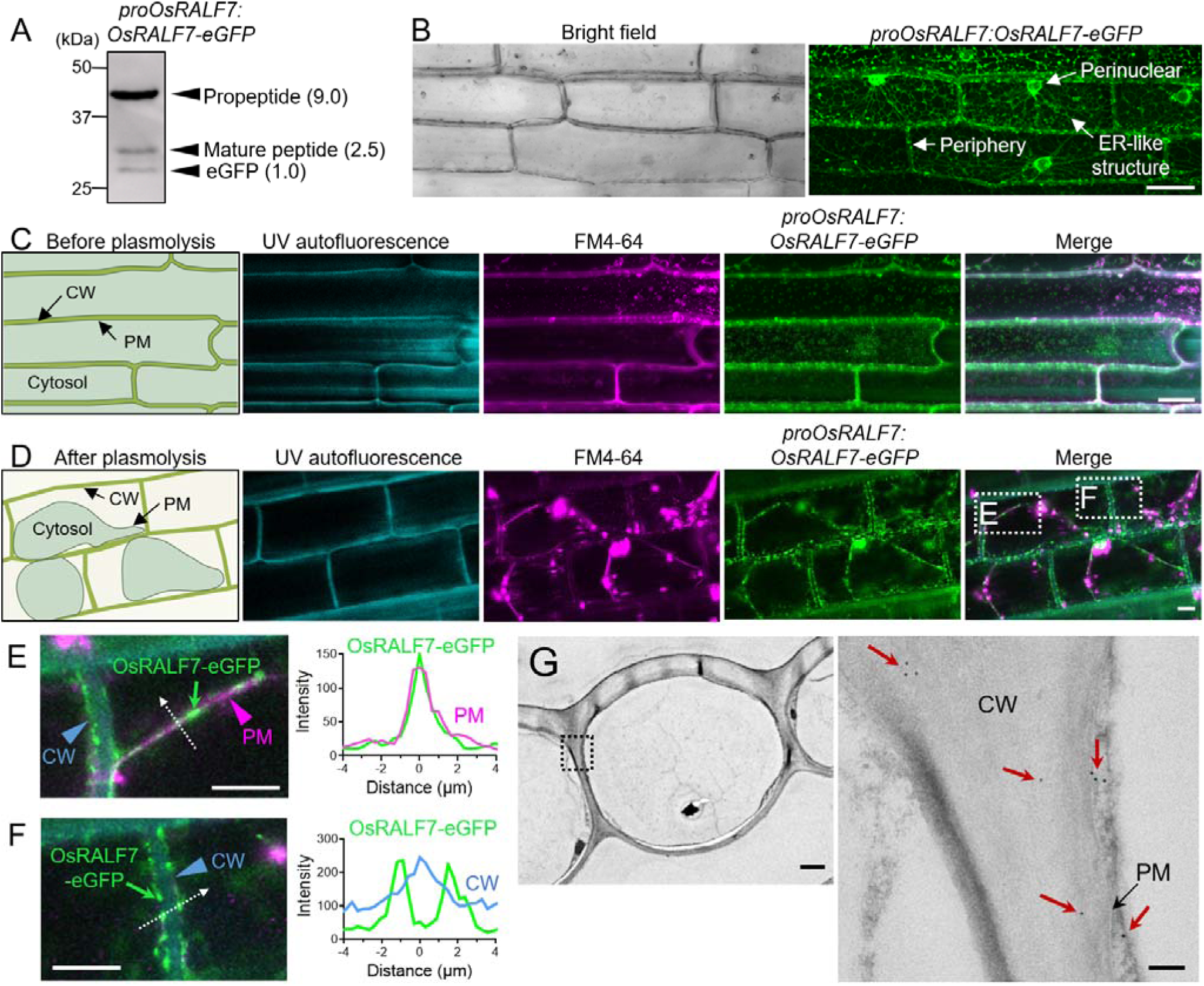
OsRALF7 associates with the cell wall and plasma membrane. (A) Immunoblot analysis of GFP in *proOsRALF7:OsRALF7*-*eGFP* transgenic plants. Total proteins were extracted from leaf sheaths of four-week-old plants and probed with an anti-GFP antibody. Bands corresponding to the predicted OsRALF7 propeptide, the processed mature form, and free GFP are indicated. Parentheses indicate the relative band intensity ratio analyzed by ImageJ, with eGFP intensity set to 1. (B) Representative confocal micrograph of OsRALF7-eGFP fluorescence in the leaf sheath of four-week-old plants. Arrows indicate eGFP signals in endoplasmic reticulum (ER)-like structures, the perinuclear region, and the cell periphery. Scale bar, 20 µm. (C) Subcellular distribution of OsRALF7-eGFP in leaf sheath cells of 4-week-old plants before plasmolysis. UV autofluorescence marks lignin-or phenolic-enriched regions of the cell wall (CW), and FM4-64 stains the plasma membrane (PM). (*D–F*) Subcellular distribution of OsRALF7-eGFP in leaf sheath cells after plasmolysis. Enlarged views of the insets in (*D*) are shown in (*E*) and (*F*), together with the corresponding line-scan analyses. Panel (*E*) shows OsRALF7-eGFP at the PM with FM4-64, whereas panel (*F*) shows OsRALF7-eGFP in the CW region. Scale bars, 10 µm. (*G*) Ultrastructural localization of OsRALF7-eGFP by immunogold labeling and transmission electron microscopy. Red arrows indicate immunogold signals. The right panel shows a higher-magnification view of the boxed region in the left panel. Scale bars, 1 μm (left) and 100 nm (right).

Confocal microscopy observations of rice leaf sheath epidermal cells revealed strong fluorescent signals from OsRALF7-eGFP in ER-like structures and perinuclear regions, suggesting that the propeptide form of OsRALF7-eGFP was accumulated and/or retained in the ER and that the fusion protein followed the conventional secretion pathway (Fig. 5*B*). Moreover, we also detected OsRALF7-eGFP signals at the cell periphery, suggesting that this fluorescence represents the mature form of OsRALF7-eGFP. To examine cell-surface localization of OsRALF7-eGFP in more detail, we stained membranes with FM4-64, a lipophilic dye commonly used to label the plasma membrane in living cells and monitor cell walls using UV autofluorescence. After FM4-64 staining, the OsRALF7-eGFP signal overlapped with that of FM4-64-positive plasma membranes at the cell periphery (Fig. 5*C*). Following plasmolysis induced by incubation in 0.75 M sucrose, we observed OsRALF7-eGFP signals along the retracted plasma membrane and at the cell periphery (Fig. 5*D*). Line-scan analysis showed that the peaks of OsRALF7-eGFP fluorescence overlap with FM4-64 fluorescence, supporting a localization of OsRALF7-eGFP at the plasma membrane (Fig. 5*E*). Two peaks of OsRALF7-eGFP fluorescence also overlapped with the rising edge of the UV autofluorescence, suggesting the localization of some OsRALF7-eGFP in the cell wall periphery (Fig. 5*F*). To further examine the ultrastructural localization of OsRALF7-eGFP, we performed immunogold labeling and transmission electron microscopy using the same OsRALF7-eGFP-expressing plants. Immunogold signals were detected in the cell wall between adjacent epidermal cells (Fig. 5G), supporting the localization of a fraction of OsRALF7-eGFP in this region. Together, these data suggest that although most detectable OsRALF7-eGFP is retained within the secretory pathway, presumably as the propeptide, a processed fraction of mature OsRALF7-eGFP may localize to the cell wall and to plasma membrane–associated compartments. We therefore tested whether OsRALF7-eGFP localization changes during infection. In infection assays using a *M. oryzae* strain accumulating cytosolic mCherry to track the position of fungal cells, OsRALF7-eGFP showed a localization pattern similar to that in non-infected tissues and did not appear to accumulate at fungal infection sites (*SI Appendix*, Fig. S14).

### Structure–function analysis of OsRALF7 and related RALFs

To identify structural features in OsRALF7 that are required for its appressorium-inducing activity, we synthesized variant peptides carrying substitutions or deletions in the conserved YISY motif or in the four conserved C-terminal cysteine residues and tested their activities on *M. oryzae* conidia (Fig. 6*A*). Alanine substitutions or deletion of the YISY motif significantly diminished appressorium formation compared to the intact peptide, consistent with the role of this motif in mediating CrRLK1L–LLG receptor interaction with *Arabidopsis* RALFs (8). By contrast, a synthetic peptide containing only the YISY sequence failed to induce appressorium formation, indicating that the YISY motif is necessary but not sufficient for inducing appressorium formation. Likewise, replacing the conserved cysteine residues with alanines markedly lowered appressorium formation, suggesting that proper folding and structural stability mediated by disulfide bonds are also essential for OsRALF7 function. Collectively, these results indicate that OsRALF7-dependent induction of appressorium formation requires both the conserved YISY motif and disulfide bond–mediated structural integrity.

**Figure 6.**
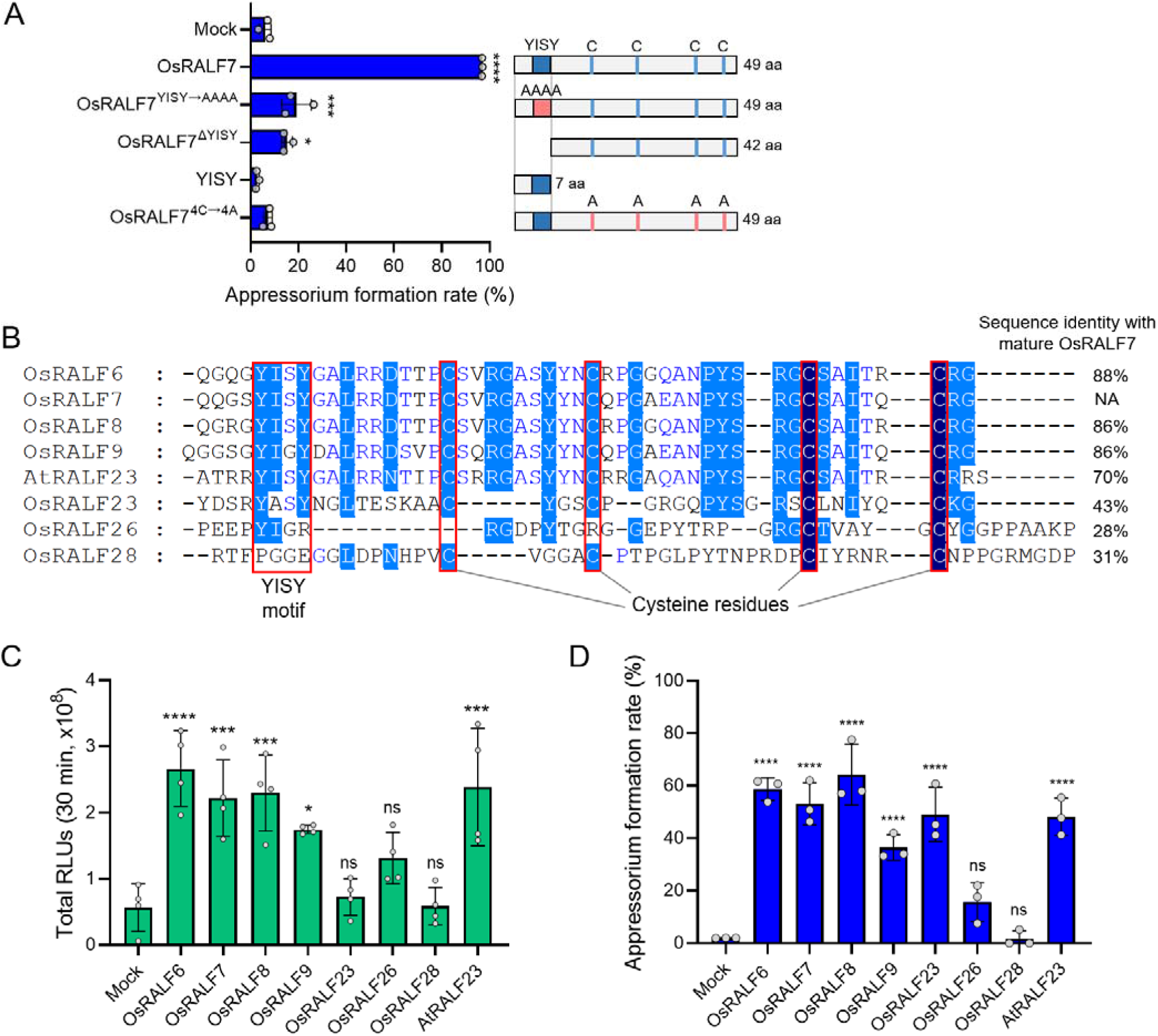
Conserved motifs in RALFs differentially regulate appressorium induction and host immune responses. (A) Roles of the conserved YISY motif and the four cysteine residues in OsRALF7 in appressorium induction. *M. oryzae* conidia were treated with 10 μM intact OsRALF7 or mutant peptides in 50% CM medium; appressorium formation rates were scored 8 h later. Appressorium formation included non-melanized, melanized, and double appressoria; conidia showing only germ tube elongation were excluded from the appressorium-formed category. Each circle represents one independently cultured plate (*n* = 3), with >100 conidia scored per plate. Error bars indicate SD. Diagrams on the right illustrate each peptide variant: OsRALF7^YISY→AAAA^, YISY-to-alanine substitutions; OsRALF7^ΔYISY^, YISY deletion; YISY, YISY motif only; OsRALF7^4C→4A^, all four cysteines replaced with alanine. (B) Multiple sequence alignment of the C-terminal sequences, downstream of the predicted RRXL protease cleavage site, of RALF peptides from rice and *Arabidopsis*. NA, Not applicable. (C) ROS production in rice suspension cells treated with 1 μM of the indicated RALF peptides or water only (Mock) control. Luminescence was measured using a luminol-based assay. Each circle represents one independently cultured flask (*n* = 4). Error bars indicate SD. RLUs, relative luminescence units. (D) Appressorium formation by various RALFs. *M. oryzae* conidia were treated with 5 μM of the indicated concentrations of nine RALFs or the unrelated peptide flg22 as a negative control in 50% CM medium; appressorium formation was quantified 8 h later. Each circle represents one independently cultured plate (*n* = 3), with >50 germinated conidia scored per plate. Error bars indicate SD. For (*A*), (*C*), and (*D*), statistical significance of differences was assessed by one-way ANOVA followed by Dunnett’s multiple-comparison

We examined whether other OsRALFs could induce appressorium formation. From the MSU Rice Genome Annotation Project, we retrieved 41 putative *OsRALF* genes and adopted the RALF numbering provided in the database annotation. Analysis of our previously published RNA-seq dataset from non-infected rice leaves showed that several *OsRALF*s, in addition to *OsRALF7*, are highly expressed in leaves (*SI Appendix*, Fig. S15) (27). We therefore selected six representative leaf-expressed *OsRALF*s: *OsRALF6*, *OsRALF8*, *OsRALF9*, *OsRALF23*, *OsRALF26*, and *OsRALF28*, covering a range in the degree of conservation of canonical RALF features (*SI Appendix*, Table S3).

Sequence comparison showed that OsRALF6, OsRALF8, and OsRALF9 are highly similar to OsRALF7 and retain all four canonical RALF features, whereas OsRALF23 carries a YASY motif in place of the YISY motif (Fig. 6*B*). OsRALF26 and OsRALF28 are more divergent and lack several canonical features. In rice suspension cells, treatment with OsRALF7, OsRALF6, OsRALF8 or OsRALF9 induced significant ROS production, whereas treatment with OsRALF23, OsRALF26 or OsRALF28 did not (Fig. 6*C*). Treatment of *M. oryzae* conidia with OsRALF6, OsRALF8 or OsRALF9 induced appressorium formation similarly to OsRALF7; treatment with OsRALF23 also showed significant activity despite its lower sequence similarity (Fig. 6*D*). By contrast, OsRALF26 and OsRALF28 were inactive in this assay. These results indicate that a subset of leaf-expressed *OsRALF*s encode peptides that can activate both host ROS production and fungal appressorium formation, while also suggesting that the sequence requirements for appressorium induction are not identical to those for ROS induction in rice.

Since OsRALF6 and OsRALF8 induced appressorium formation to an extent similar to OsRALF7, we examined their effects on *M. oryzae* appressorium formation *in planta*. We generated rice plants carrying double knockouts of *OsRALF6* and *OsRALF8* (*SI Appendix*, Fig. S16). Compared with wild-type plants, the *Osralf6/Osralf8* double-knockout plants showed reduced appressorium formation and an altered distribution of appressoria relative to anticlinal cell boundaries, similar to the phenotype observed in *Osralf7* mutant plants (*SI Appendix*, Fig. S17). These results suggest that OsRALF6 and OsRALF8, like OsRALF7, contribute to appressorium formation and may influence infection-site selection *in planta*.

To place these findings in the context of the extensive *Arabidopsis* RALF literature, we tested whether *Arabidopsis* RALF peptides also affect appressorium formation in *M. oryzae*. We estimated the phylogenetic distances between mature OsRALF7 and 37 mature *Arabidopsis* RALFs based on their amino acid sequences (28). This analysis identified AtRALF23 as the closest *Arabidopsis* homolog to OsRALF7 (*SI Appendix*, Table S4). Mature AtRALF23 shares approximately 70% aa sequence identity with mature OsRALF7 and shares all four hallmark features of RALFs also present in OsRALF7 (Figs. 1*B* and 6*B*). Treatment of rice suspension cells with synthetic mature AtRALF23 elicited ROS production (Fig. 6*C*). Moreover, treatment of *M. oryzae* conidia with mature AtRALF23 induced appressorium formation to a similar extent as OsRALF7 (Fig. 6*D*).

### Divergent roles of RALF family members in appressorium formation in *Colletotrichum higginsianum*

We investigated whether the ability of RALFs to induce appressorium formation is a conserved response among appressorium-forming fungi or is specific to *M. oryzae*. To this end, we examined the effects of RALF treatment on a distinct fungal species, namely *C. higginsianum*. *M. oryzae* and *C. higginsianum* both belong to the class Sordariomycetes and employ similar infection strategies, including the formation of melanized appressoria and host penetration driven by turgor pressure and cell wall–degrading enzymes (46). Because *C. higginsianum* infects Brassicaceae hosts, we tested OsRALF7 as well as AtRALF23 in appressorium formation assays. When synthetic OsRALF7 or AtRALF23 was applied to *C. higginsianum* in CM medium, neither peptide induced appressorium formation across a range of concentrations (Fig. 7*A*). This result suggested that RALF-dependent appressorium induction is not a universal feature of appressorium-forming fungi. Notably, 10 μM AtRALF23 markedly inhibited conidial germination and appressorium formation of *C. higginsianum*. We next performed the same assay under water-culture conditions. Under these conditions, OsRALF7 inhibited conidial germination at 5 μM, with a corresponding drop in appressorium formation at 10 μM, whereas 1 μM AtRALF23 completely suppressed conidial germination and the associated appressorium formation.

**Figure 7.**
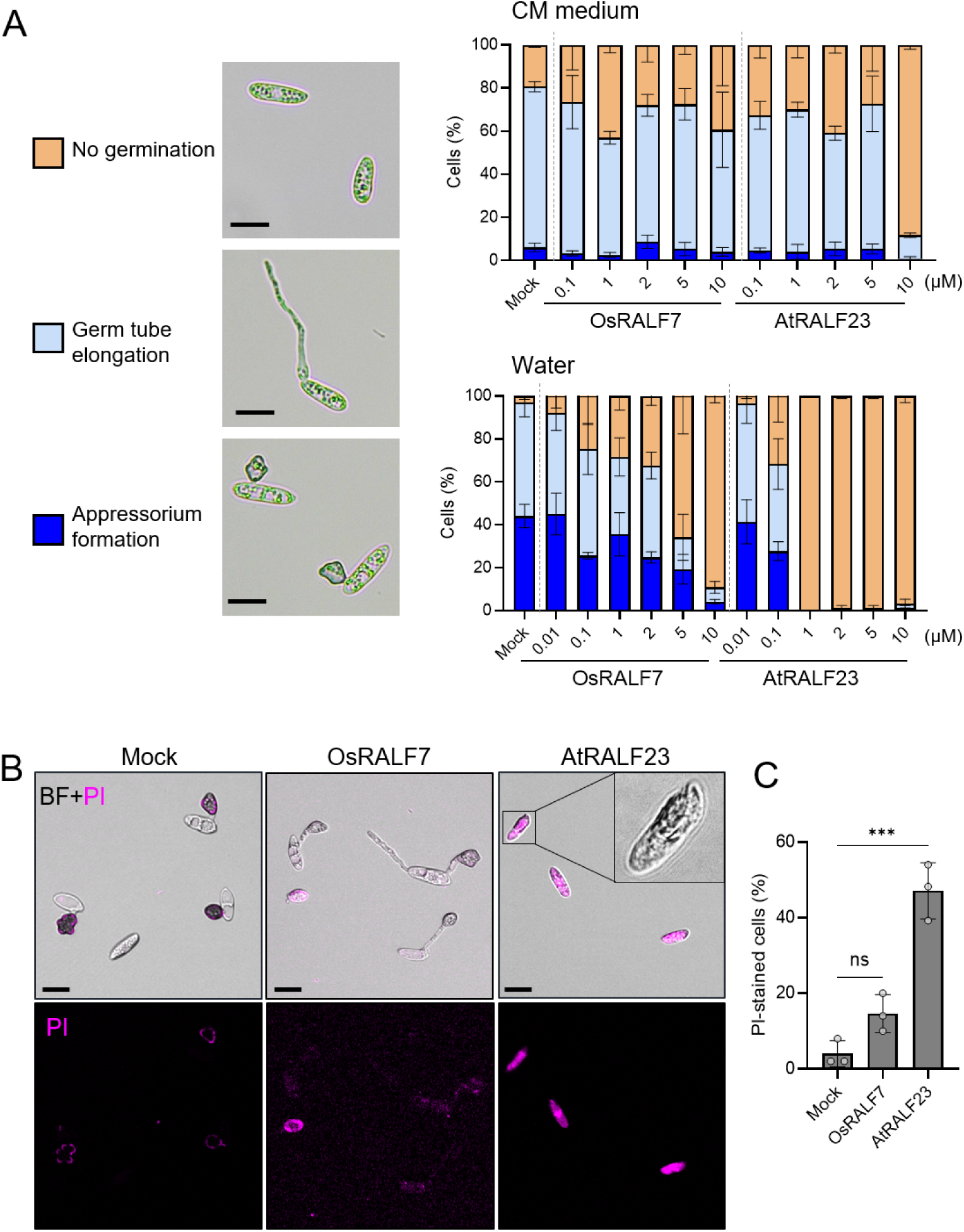
*Arabidopsis* RALF23 exhibits antimicrobial activity against *C. higginsianum*. (A) OsRALF7 and AtRALF23 do not induce appressorium formation in *C. higginsianum*, whereas AtRALF23 suppresses conidial germination of *C. higginsianum. C. higginsianum* conidia were treated with water only (Mock), OsRALF7, or AtRALF23 at the indicated concentrations and incubated either on 25% CM medium for 10 h or in water for 20 h. For each treatment and condition, at least 50 conidia were assessed per independently cultured plate (*n* = 3) and classified into three morphological categories (no germination, germ tube elongation, and appressorium formation); the percentage of conidia in each category was calculated. Appressorium formation included both non-melanized and melanized appressoria. Error bars indicate SD. Experiments were performed three times, each time with similar results. Scale bars, 10 µm. (B) Propidium iodide (PI) staining reveals the antimicrobial activity of AtRALF23. *C. higginsianum* conidia were incubated in water with water only (Mock), OsRALF7 (1 μM), or AtRALF23 (1 μM) for 20 h, followed by PI staining. Higher-magnification images highlight the abnormal cellular morphology with condensed cytoplasm observed after AtRALF23 treatment. Representative epifluorescence micrographs of PI-stained cells are shown. Experiments were performed three times, each time with similar results. Scale bars, 10 µm. BF, Bright field. (C) Quantification of PI-positive cells. Each circle represents one biological replicate (*n* = 3), with 50 cells scored per replicate. Statistical significance of differences was assessed by one-way ANOVA followed by

Microscopy analysis of AtRALF23-treated conidia under water-culture conditions revealed abnormal cellular morphology with condensed cytoplasm, suggestive of cell death– associated changes (Fig. 7*B*) (47). To test whether AtRALF23 induces cell death, we performed propidium iodide (PI) staining, which detects dead or dying cells with compromised plasma membranes. Indeed, AtRALF23 treatment substantially raised the proportion of PI-positive *C. higginsianum* cells under water-culture conditions (Fig. 7*B* and *C*), indicating that inhibition of germination is associated with loss of membrane integrity and direct cytotoxic activity rather than a simple delay in germination. Treatment with OsRALF7 showed a modest and nonsignificant increase in the fraction of PI-positive cells relative to mock. Together, these results suggest that AtRALF23 may exhibit antifungal and/or cytotoxic activity against *C. higginsianum*.

We examined whether similar effects occur in *M. oryzae* by treating conidia with OsRALF7 or AtRALF23 under water-culture conditions. Consistent with our observations in *C. higginsianum*, AtRALF23 suppressed conidial germination and appressorium formation in *M. oryzae* at concentrations ≥5 μM (*SI Appendix*, Fig. S18). By contrast, OsRALF7 had no measurable effect on germination or appressorium formation at concentrations up to 10 μM. Collectively, these findings suggest that RALF family members elicit distinct outcomes depending on fungal species and environmental conditions, ranging from appressorium induction to growth inhibition and cell death. Notably, AtRALF23 exhibited stronger antimicrobial activity against both *C. higginsianum* and *M. oryzae* than OsRALF7.

## Discussion

This study demonstrates that plant-derived RALFs have multifaceted functions during plant– fungal interactions: they activate host immune responses, but they can also promote appressorium formation in *M. oryzae* and may provide spatial cues for host penetration (Fig. 8). In addition, RALFs exhibit direct antimicrobial activity against *C. higginsianum*, indicating that they contribute to antifungal defense not only through immune regulation but also through direct antimicrobial effects.

**Figure 8.**
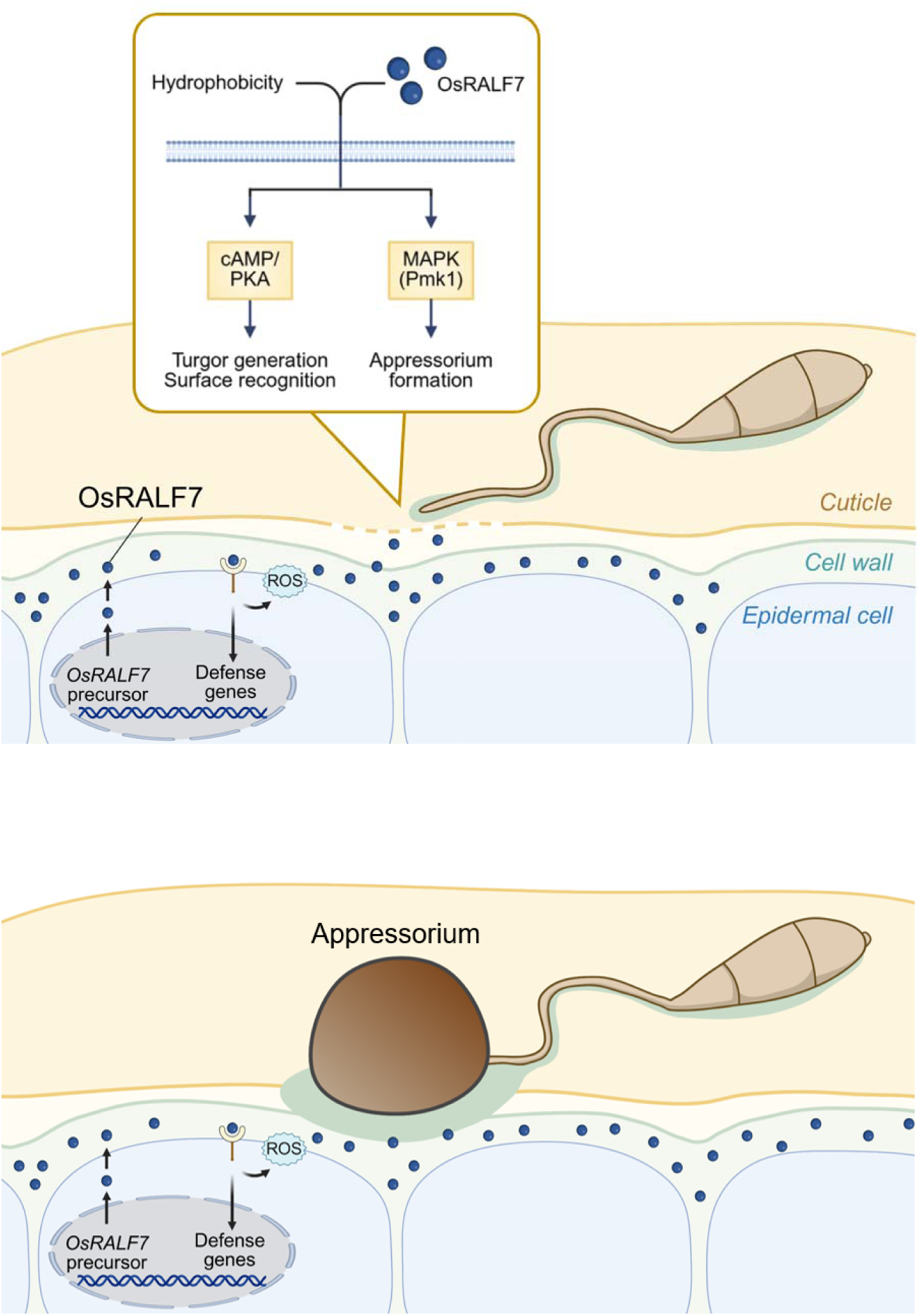
Model illustrating the dual roles of OsRALF7 in rice–*M. oryzae* interactions. Rice-derived OsRALF7 promotes host immunity by inducing ROS production and the expression of defense-related genes. At the same time, *M. oryzae* exploit OsRALF7 as a host-derived cue during surface sensing, thereby promoting appressorium formation and influencing appressorium positioning relative to anticlinal cell boundaries. Together with hydrophobic surface cues, OsRALF7 is associated with cAMP/PKA- and Pmk1-associated signaling and appressorium formation.

### Host-derived OsRALF7 promotes appressorium in *M. oryzae*

Appressorium formation is an essential developmental stage for *M. oryzae* to successfully infect plant hosts. The formation of this infection structure is induced by physicochemical cues, including leaf-surface hydrophobicity, surface rigidity, and the presence of primary alcohols, which trigger downstream signaling cascades via fungal GPCRs and cell-surface receptors (18). However, the induction of appressorium formation by plant peptide or proteins has not been previously identified.

In this study, we identified rice-secreted OsRALF7 as a host-derived peptide that promotes appressorium formation in *M. oryzae*. We first showed that mature OsRALF7 activates rice immunity (Fig. 1), consistent with previous studies in other plant species (3, 48). We then treated *M. oryzae* with OsRALF7 under *in vitro* conditions lacking plant tissue. Treatment with OsRALF7 promoted appressorium formation even under conditions that normally support vegetative growth and inhibit appressorium development (Fig. 2). Subsequent analysis showed that OsRALF7 treatment activated both cAMP/PKA and Pmk1 MAPK pathways and this response requires a hydrophobic surface (Fig. 3). Additionally, the appressorium formation was reduced on *Osralf7* mutant plants, which also exhibited following reduced host-cell penetration and invasive hyphal development (Fig. 4*A*, *B*, and *E*). Conversely, exogenous application of OsRALF7 increased fungal biomass in the infected rice leaves (Fig. 4*F* and *G*). Collectively, these findings suggest that *M. oryzae* responds to host-derived OsRALF7 by inducing appressorium formation through cAMP/PKA and Pmk1 MAPK pathways together with hydrophobic input, thereby supporting rice invasion (Fig. 8).

Supporting evidence for this concept comes from independent studies of the interaction between *Arabidopsis* and the powdery mildew fungus *Erysiphe cruciferarum* (*Ecr*) (49). Higher-order mutants such as *ralf22 ralf23 ralf31* and *ralf1 ralf18 ralf22 ralf23 ralf31 ralf33 ralf34* exhibited enhanced resistance to *Ecr*, as indicated by diminished conidiophore production (i.e., formation of asexual spore-bearing structures) and lower fungal biomass. These findings suggest that several Arabidopsis RALFs promote, or otherwise support, fungal colonization. *Ecr* conidiophore formation also depends on host cell-wall integrity and a stable apoplastic pH, process associate with canonical RALF function. Thus, host-derived RALF signaling may affect fungal development by altering the host environment. Moreover, analyses of *fer* mutants suggested that synthetic AtRALF23 or *AtRALF23* overexpression could promote *Ecr* conidiophore formation, even when canonical FER-dependent host responses were compromised. Together, these observations suggest that RALF peptides can influence fungal development through mechanisms that are not explained solely by their established roles in host immunity.

In parallel, RALF mimics have been identified in various plant-associated microorganisms, including pathogenic and symbiotic fungi, as well as in root-knot nematodes; these RALF mimics are secreted during host colonization and promote pathogen/symbiont fitness (50–54). By contrast, *Ecr* and *M. oryzae* lack genes encoding RALF mimics in their genome (51), raising the possibility that these fungi instead exploit endogenous host RALF signaling to support colonization. Together with our findings, these observations support an emerging model in which host-derived peptide signals can directly modulate fungal morphogenetic programs during plant infection.

### RALFs may as a spatial cue for appressorium formation

Our study showed that OsRALF7 both promote appressorium formation by *M. oryzae* and possibly contribute to its spatial positioning. On wild-type plants, appressoria preferentially contacted anticlinal cell boundaries, whereas this spatial preference was lost in *Osralf7* mutant plants (Fig. 4*D*). This finding suggests that OsRALF7 contributes not only as a general inducer of appressorium formation but also may act as a positional cue. Preferential targeting of anticlinal walls has been reported in other plant pathogens, including *Colletotrichum* species (20, 21, 55), suggesting that these regions may represent favorable sites for host entry. However, the underlying cues do not appear to be universally conserved. In this study, neither OsRALF7 nor AtRALF23 induced appressorium formation in *C. higginsianum* (Fig. 7*A*), arguing against a general role for RALFs in anticlinal targeting across fungi. In *B. maydis*, by contrast, appressorium formation at anticlinal walls has been linked to recognition of pectin, which is enriched in these regions (22). Together, these observations suggest that although anticlinal regions are recurrently targeted by diverse pathogens, different species may rely on distinct host-derived cues and sensing mechanisms to recognize them.

There are three non-exclusive mechanisms by which OsRALF7 could influence appressorium positioning. The first is a direct RALF spatial cue model: RALFs may act as direct spatial cues for appressorium positioning by accumulating preferentially at anticlinal cell boundaries. This idea is supported by studies in *Arabidopsis* showing that RALF binds to demethylesterified pectin and associates with LRXs that contribute to cell-wall architecture (11). Although the distribution of demethylesterified pectin in rice epidermal cells remains unclear, the pectin-rich middle lamella between adjacent cells could provide a matrix for local retention of OsRALF7. OsRALF7 may therefore be present near anticlinal cell boundaries before infection or become locally accessible following fungal modification of the host surface. Consistent with its association with the cell wall, immunogold transmission electron microscopy detected OsRALF7-eGFP–associated signals in the cell wall between adjacent epidermal cells (Fig. 5*G*). However, the low immunogold-labeling signal intensity, potentially resulting from technical limitations of leaf-sheath section analysis, prevented quantitative assessment of its spatial distribution.

The second mechanism by which OsRALF7 could influence appressorium positioning is a RALF induced pH gradient model: RALF signaling can inhibit proton pump activity via CrRLK1L–LLG pathways, causing local apoplastic alkalinization (5, 8). If such inhibition occurs at anticlinal cell boundaries, the resulting pH change could be sensed by *M. oryzae* to trigger appressorium formation. Our *in vitro* finding that alkaline conditions promote appressorium formation is consistent with this idea (*SI Appendix*, Fig. S19). In addition, the pH of rice apoplastic fluid becomes more alkaline after infection by *M. oryzae* (56), raising the possibility that infection-associated apoplastic alkalinization contributes to appressorium formation.

Finally, OsRALF7 may influence appressorium positioning through a multifactorial recognition model: *M. oryzae* may integrate direct RALF-dependent spatial cues and RALF-induced pH changes, together with secondary RALF-mediated effects on cell-wall integrity (11, 12), ROS production (57) or phytohormonal signaling (58). In this study, we showed that leaf-surface topography, hydrophobicity, and cuticular wax composition did not differ significantly between wild-type and *Osralf7* mutant plants (*SI Appendix*, Figs. S9 and S10). These results suggest that major changes in these known appressorium-inducing surface properties are unlikely to explain the altered appressorium distribution. Future studies addressing other local features associated with anticlinal cell boundaries may reveal additional spatial cues that regulate appressorium positioning in *M. oryzae*.

The use of RALFs as positional cues may be particularly advantageous for *M. oryzae*, as a hemibiotrophic pathogen that initially colonizes living host cells before transitioning to necrotrophic growth (59). For hemibiotrophic fungi, successful infection likely depends on the ability to recognize living, physiologically active host tissue. In this context, host-derived RALF signals could help the pathogen rapidly detect suitable infection sites on active host cells.

### Possible mechanisms by which RALFs promote appressorium formation

In *M. oryzae*, the cAMP/PKA and Pmk1 MAPK pathways are essential regulators of appressorium development and pathogenicity (34). Pth11 is a well-characterized GPCR involved in sensing surface hydrophobicity and signaling primarily through the cAMP/PKA pathway (19). By contrast, the membrane sensors Msb2 and Sho1 recognize surface hydrophobicity, cutin monomers, and primary alcohols and activate primarily the Pmk1 MAPK pathway to promote appressorium formation (18). Several observations suggest potential crosstalk between the cAMP/PKA and Pmk1 MAPK pathways (60). For example, MoRas2 functions upstream of both cAMP/PKA and Pmk1 MAPK signaling (61), and simultaneous loss of the two PKA catalytic subunits, CpkA and Cpk2, increases Pmk1 phosphorylation (42). The involvement of multiple surface-sensing components and the potential crosstalk between their downstream pathways may enable *M. oryzae* to respond to diverse host-surface signals.

Our results suggest that the OsRALF7 response is integrated with surface sensing and downstream cAMP/PKA and Pmk1 MAPK signaling. OsRALF7 increased intracellular cAMP levels and Pmk1 phosphorylation in germinated conidia (Fig. 3*B* and *C*), indicating OsRALF7 activates both pathways. By contrast, these responses were not detected in vegetative mycelia (*SI Appendix*, Fig. S4), suggesting that responsiveness to OsRALF7 might be specific to the infection-related morphogenesis. Consistent with these biochemical responses, OsRALF7 induced genes associated with appressorium maturation, melanin biosynthesis, and host penetration (*SI Appendix*, Fig. S5). Many of these genes overlapped with those previously reported to be induced by cAMP treatment or down-regulated in the Δ*pmk1* mutant (37, 40, 41), further linking OsRALF7 to downstream transcriptional programs regulated by these pathways. Furthermore, the effect of OsRALF7 on appressorium formation depended on a hydrophobic surface. OsRALF7 failed to induce appressorium formation on a hydrophilic GelBond surface at all concentrations tested, whereas exogenous cAMP induced appressoria under the same conditions (Fig. 3*E*). The inability of OsRALF7 to substitute for a hydrophobic-surface cue suggests that OsRALF7 does not act as an autonomous trigger of appressorium formation but instead cooperates with signaling initiated by hydrophobic-surface perception.

Analysis of surface-sensor mutants further supports this interpretation. Deletion of either *PTH11* or *SHO1* reduced OsRALF7-induced appressorium formation but did not abolish responsiveness to OsRALF7 (Fig. 3*D*; *SI Appendix*, Fig. S6). Thus, Pth11 and Sho1 contribute to the efficiency of this response but do not individually account for it. The residual responsiveness of these mutants may reflect functional compensation by other surface sensors and/or crosstalk between the cAMP/PKA and Pmk1 MAPK pathways. The Δ*msb2* mutant failed to form appressoria in response to OsRALF7. However, its broad defect in appressorium formation under multiple conditions precluded an OsRALF7-specific interpretation. Moreover, the severe phenotypes of the Δ*pth11*, Δ*msb2*, and Δ*sho1* mutants may be influenced by the P2 strain background, because less severe defects have been reported in other *M. oryzae* strain backgrounds (18, 19, 43).

Together, these findings raise the possibility that OsRALF7 reinforces signaling initiated by hydrophobic-surface perception through the cAMP/PKA and Pmk1 MAPK pathways. In this model, inputs from multiple surface sensors, including Pth11, Msb2, and Sho1, activate these pathways, and OsRALF7 cooperates with these inputs to promote appressorium formation and subsequent infection. This model explains why disruption of individual surface sensors reduces but does not abolish responsiveness to OsRALF7 and why OsRALF7 cannot induce appressorium formation on a hydrophilic surface. Thus, OsRALF7 may function not as an autonomous trigger but as a host-derived signal that reinforces the developmental response of germinated conidia after they encounter a surface favorable for infection.

### RALF peptides have antimicrobial activity

Our results indicate that RALFs, particularly AtRALF23, exert clear antimicrobial activity against *C. higginsianum* and weaker activity against *M. oryzae* (Fig. 7). These observations raise the possibility that the antifungal effects of RALFs are not restricted to *Colletotrichum* species but may extend more broadly across filamentous fungi. One feature that distinguishes AtRALF23 from the less active OsRALF7 is its higher arginine content and correspondingly higher isoelectric point (AtRALF23, 11.36; OsRALF7, 8.87), which may help explain the difference in activity. Consistent with this idea, RALFs have been classified as defensin-like peptides (31, 32); plant defensins are cysteine-rich antimicrobial peptides that often act through interactions with microbial surface or membrane components (2). The greater positive charge of AtRALF23 may therefore facilitate initial electrostatic interactions with negatively charged components on the fungal cell surface, potentially promoting membrane disruption and/or peptide uptake. Positively charged RALFs have also been shown to associate with the plant cell wall through electrostatic interactions with demethylesterified pectin (11). It is therefore conceivable that RALFs accumulate in the host cell wall and are released locally during pathogen-induced degradation of the cell wall or cuticle, after which they interact with fungal surface structures to exert antifungal activity. Such properties raise the possibility that RALFs are bifunctional molecules that integrate host signaling with direct modulation of fungal behavior. Indeed, other plant-derived peptides, such as the PR1-derived peptide CAPE1, are shown to combine immune-eliciting activity with direct antimicrobial effects (62). At the same time, it remains unclear why the antifungal activity of OsRALF7 and AtRALF23 differs between *C. higginsianum* and *M. oryzae*. Possible explanations include differences in fungal cell-wall architecture that influence peptide access to the fungal plasma membrane, as well as differences in fungal plasma-membrane composition or surface charge that affect peptide binding and susceptibility.

### Potential agricultural applications of RALFs

RALFs have three distinct functions: immunity inducer, possible spatial cue for *M. oryzae*, and antifungal activity against *C. higginsianum.* If these functions could be strategically coordinated, RALFs would be promising targets for innovative non-transgenic disease control strategies. For example, RALFs engineered to disable their activity as possible spatial cues for pathogens while retaining their immune-inducing and antifungal activities might be used for crop protection. Another possible strategy is the application of exogenous RALFs to crops (e.g., through spraying); this might induce ectopic appressorium formation in *M. oryzae,* decreasing the infection rate.

## Materials and Methods

### Plant and fungal materials and growth conditions

Rice (*Oryza sativa* L. ssp. *japonica* cv. Nipponbare, Moukoto, and LTH and *indica* cv. CO-39) plants were used in this study. Seeds were incubated on moist filter paper at 30°C for 48 h to induce germination. Germinated seedlings were then sown in Kumiai Ube Baido No. 2 nutrient-rich soil pellets (MC Ferticom) without any fertilizer in a seedling case of 4.5 × 13.5 × 9.5 cm, and grown in a controlled-environment room under a 16-h light/8-h dark photoperiod with illumination provided by plant growth fluorescent lamps (Plantlux FL40S BRN, Toshiba) at a PPFD of 100 µmol m^-2^ s^-1^ and a 25°C/20°C (day/night) cycle for the infection and growth assays. For the leaf sheath assays, plants were grown in a controlled-environment room as described above, or a growth chamber (Nippon Medical & Chemical Instruments) under a 16-h light/8-h dark photoperiod with fluorescent lighting provided by white fluorescent lamps at a PPFD of 250 µmol m^-2^ s^-1^ with a 30°C/25°C (day/night) cycle, 70% relative humidity, and 475 ppm CO_2_. Rice suspension cells derived from Nipponbare calli were cultured in R2S medium (63). For fungal materials, all *M. oryzae* isolates were stored on dried filter paper or sterilized barley seeds. Isolates were first cultured on potato sucrose agar (PSA) or potato dextrose agar (PDA) (Shimadzu) slants for more than 1 week at 25°C. Mycelial plugs from these slant cultures were then transferred to oatmeal agar medium (40 g oatmeal, 5 g sucrose, and 20 g agar per L) and incubated at 25°C for 1 week. After the aerial hyphae were removed by gently scraping the colony surface, the plates were incubated at 25°C under black light BM-20BLB/50 (Tokyo Metal Industrial) for 4 to 7 days to induce sporulation. *C. higginsianum* was routinely recovered from glycerol stocks on PDA and incubated under black light at 25°C for 1 week. Fungal strains used in this study are listed in *SI Appendix*, Table S2.

### ROS production assay

Analysis of ROS production after peptide or chitin treatment of rice suspension cells was performed as described previously (64) with minor modifications. Briefly, rice suspension cells were cultured in R2S medium for 1 week, transferred to fresh R2S medium, and incubated for 1 day at 28°C. Cells were dispensed into the wells of a 96-well plate at approximately 20 mg per well and incubated in the dark at 28°C for 16 h. The medium was then replaced with fresh R2S medium containing 0.5 mM L-012 and 10 μg/mL peroxidase, and the cells were treated with peptides at the indicated concentrations with or without chitin (hexaacetylchitohexaose; Biosynth) at a final concentration of 1 μg/mL. Water was used as a mock control. At least three biological replicates were prepared for each treatment. ROS levels were determined by measuring L-012-dependent chemiluminescence with a multimode reader TriStar2 LB 942 (Berthold Japan).

### *In vitro* appressorium formation assay

Conidia were harvested in sterile water and filtered once through Miracloth. *M. oryzae* conidia were resuspended at 2 × 10^5^ conidia/mL, whereas *C. higginsianum* conidia were resuspended at 5 × 10^5^ conidia/mL. Depending on the experiment, conidia were suspended in sterile water or CM diluted to 25%, 30%, or 50% of its full strength. Full-strength CM contained 0.3% (w/v) yeast extract, 0.3% (w/v) Casamino acids, and 0.5% (w/v) sucrose. Drops of 5 μL of the conidial suspension were placed onto glass coverslips (Muto pure chemicals 25602) and incubated in a moist chamber at 25°C. Water was used as a mock control. Conidial germination and appressorium formation were examined at the indicated times under a light microscope Primostar 3 (ZEISS).

### Fungal infection spray assay

Three-week-old rice seedlings were used, when the third leaf was fully expanded. Just before fungal inoculation, the third leaves were fixed onto a plastic board with rubber bands. Infection assays were conducted according to a previously described protocol (65), with minor modifications. Conidia of *M. oryzae* were harvested, filtered once through Miracloth (Merck Millipore), and resuspended in water. For the OsRALF7 peptide-treatment assay, the conidial suspension was adjusted to 3.5–5.0 × 10^5^ conidia/mL and supplemented with OsRALF7 peptide to a final concentration of 5 μM, or water was added as a mock control. A total of 10– 15 mL of the conidial suspension containing 0.01% Tween 20 was sprayed onto the adaxial surfaces of the third leaves of 21 plants using an air compressor. For infection assays with wild-type and *Osralf7* rice plants, the conidial suspension was adjusted to 2 × 10^5^ conidia/mL, and 20 mL of the suspension containing 0.01% Tween 20 was sprayed onto the adaxial surfaces of the third leaves of 15 plants. The inoculated seedlings were incubated in darkness under high humidity for 24 h and then transferred to dry conditions under fluorescent light with a 16-h light/8-h dark photoperiod at 25°C. Symptoms and sample collections were evaluated 5, 6 or 7 days after inoculation based on lesion size and color. The respective fungal biomass was examined by qPCR using specific primers for *Pot2* of *M. oryzae* and normalized to the reference gene *OsUBQ* (66).

### Rice leaf sheath assay

Rice leaf sheath inoculation was conducted as described previously (67), with minor modifications. Conidia were harvested in sterile water, filtered once through Miracloth, and resuspended to 5×10^4^ conidia/mL in 0.25% (w/v) gelatin. The conidial suspension was infiltrated with a micropipette into dissected leaf sheaths obtained from the four-or five-week-old rice plants until the inner cavity was completely filled. Samples were incubated in a sealed moist chamber in the dark at 28 °C for the indicated times. After incubation, leaf sheaths were trimmed with razor blades to obtain thin sections for microscopy. For invasive hypha staining, a 2× trypan blue staining solution containing 10 mL lactic acid, 10 g phenol, 10 mL glycerol, 10 mL water, and 10 mg trypan blue was diluted with ethanol at a 1:2 ratio immediately before use. Leaf sheath sections were immersed in the diluted staining solution and heated in a boiling-water bath until the solution began to boil. The samples were then incubated for 3 min, removed from the heat while remaining in the water bath, and allowed to cool for 30 min.

Subsequently, the samples were transferred to a preservation solution consisting of lactic acid and glycerol (1:1, v/v) and stored until observation by light microscopy. For OsRALF7-eGFP localization, leaf sheaths from four-week-old rice plants were trimmed and observed directly. For plasma-membrane staining, trimmed leaf sheath sections were mounted in 10 μM FM4-64 FX (Thermo Fisher Scientific) prepared in 10 mM HEPES buffer (pH 7.4) and incubated in the dark for 30 min. Plasmolysis was induced by treating the stained sections with 0.75 M sucrose. In both cases, samples were observed directly under a confocal microscope without washing.

### Confocal microscopy and image analysis

Imaging was performed under an FV4000 confocal microscope (EVIDENT). Fluorescence detection settings were: UV autofluorescence, excitation at 359 nm, emission collected at 430–470 nm; eGFP, excitation at 488 nm, emission collected at 500–580 nm; FM4 64, excitation at 515 nm, emission collected at 668–728 nm; propidium iodide, excitation at 538 nm, emission collected at 570–620 nm. Line scan analyses were performed using ImageJ (https://imagej.net/ij/). Z stacks were acquired with optical section intervals of 0.39 μm for the leaf sheath samples stained with FM4-64, and 0.70 μm for the leaf sheath sample without staining.

### Immunogold labeling and transmission electron microscopy

Leaf-sheath samples from OsRALF7-eGFP-expressing rice plants and nontransgenic wild-type plants used as a negative control were processed by rapid freezing, freeze substitution, and LR White embedding. Ultrathin sections were labeled with an anti-GFP antibody and a 10-nm gold-conjugated secondary antibody and examined by transmission electron microscopy. Detailed procedures are provided in *SI Appendix*, Materials and Methods.

### Sample preparation for cAMP, Pmk1 phosphorylation, and RNA-seq analyses in *M. oryzae* germinated conidia

Germinated *M. oryzae* conidia were treated with the indicated peptides in diluted complete medium and incubated as droplets on glass coverslips at 25°C. For cAMP and Pmk1 phosphorylation analyses, conidia were incubated in 25% CM at 5.0 × 10^5^ conidia/mL; for RNA-seq analysis, conidia were incubated in 30% CM at 3.0 × 10^5^ conidia/mL. Conidial germination and appressorium formation were verified before fungal material was collected for cAMP measurement, immunoblot analysis, or RNA extraction. Detailed sample preparation, extraction, and analysis procedures are provided in *SI Appendix*, Materials and Methods.

### cAMP analysis

Intracellular cAMP levels were measured as described previously (Liu et al.; Zhang et al.), with modifications. Lyophilized fungal material was weighed to determine dry weight before cAMP extraction. Intracellular cAMP was extracted with trichloroacetic acid, quantified using the cAMP-Screen Direct Cyclic AMP Immunoassay System (Thermo Fisher Scientific), and normalized to fungal dry weight. Detailed extraction and assay procedures are provided in *SI Appendix*, Materials and Methods.

### Immunoblot analyses

Pmk1 phosphorylation in *M. oryzae* protein extracts was assessed by immunoblotting with a phospho-p44/42 MAPK (Erk1/2) (Thr202/Tyr204) antibody (Cell Signaling Technology, 4370S). Equal amounts of protein (20 µg per sample) were analyzed by SDS–PAGE and immunoblotting. OsRALF7-eGFP expression in rice leaf-sheath extracts was assessed by immunoblotting with a GFP antibody B-2 (Santa Cruz Biotechnology, sc-9996). Detailed protein-extraction and immunoblotting procedures are provided in *SI Appendix*, Materials and Methods.

### Peptide synthesis

All synthetic peptides were mature peptides designed based on the conserved C-terminal regions listed in *SI* Appendix, Table S5. The synthesized peptides are listed in *SI Appendix*, Table S5, with purities exceeding 90% (SCRUM Inc., Japan).

### Statistical analysis

Statistical analyses were performed using GraphPad Prism 11 (GraphPad Software, USA). Differences among multiple groups were assessed using one-way analysis of variance (ANOVA) followed by Dunnett’s multiple-comparisons test or Tukey’s multiple-comparisons test. Comparisons between two groups were performed using unpaired two-tailed Welch’s *t*-tests.

## Data Availability

The RNA-seq reads generated in this study have been deposited in the DDBJ Sequence Read Archive. Reads from rice suspension cells treated with OsRALF7 or chitin are available under accession numbers DRR959814–DRR959822 and are associated with BioProject PRJDB42064 and BioSamples SAMD01899517–SAMD01899525. Reads from *M. oryzae* treated with OsRALF7 are available under accession numbers DRR963041–DRR963052 and are associated with BioProject PRJDB42122 and BioSamples SAMD01908479– SAMD01908490.

## Acknowledgments

We thank Keiichiro Nemoto and Kiwamu Hyodo for their advice on the conception of this study. We also thank Yasuyuki Kubo, Sayo Kodama, and Ken Harata for pBITEFpro:mCherry plasmid and sharing the method for *M. oryzae* transformation. We thank Kenichi Ikeda for kindly providing LTH and CO-39 seeds. This work was supported in part by PRESTO from a JST grant (JPMJPR22D8), JSPS KAKENHI grants (21K14856 and JP16H06279 (PAGS)) for F.F., JSPS KAKENHI grants (23K18030, 24H01371), JSPS Bilateral Programs (JPJSBP20237408), the Ohara foundation, the Yakumo Foundation for Environmental Science, and the Ryobi Teien Memory Foundation for Y.K., and the Joint Usage/Research Center, the Institute of Plant Science and Resources, and JSPS J-PEAKS grant (JPJS00420230010). Some diagrams were created with BioRender.com.

## Author Contributions

F.F. and Y.K. designed research; F.F., S.F., N.K., Y.Hino, A.S., I.G., F.F.D., T.G., N.O., H.N., W.C., M.K., and P.W. performed research; M.S., Y.Harada, T.A., S.A., H.M., and D.S., contributed new reagents/analytic tools; F.F., T.F., and M.S. analyzed the data; F.F. and Y.K. wrote the paper.

