## Supplemental file for "Rice blast fungus hijacks a rice RALF peptide to initiate infection"

### Supporting Information Text

### Extended materials and methods

#### Plasmid construction and transformation

Rice *Osralf7* and *Osralf6/Osralf8* mutants were generated using the CRISPR–Cas9 system (1). Guide RNA target sequences were designed using the CRISPR-P v2 web tool (http://crispr.hzau.edu.cn/cgi-bin/CRISPR2/CRISPR). To construct the gRNA expression plasmids, three DNA fragments were PCR-amplified from the pU6gRNA plasmid template: (i) a partial pUC19 backbone fragment containing the OsU6−2 promoter fused to the *OsRALF7* or *OsRALF6*/*OsRALF8* gRNA target sequence, (ii) a fragment containing the *OsRALF7* or *OsRALF6*/*OsRALF8* gRNA target sequence fused to the gRNA scaffold followed by a poly(T) terminator, and (iii) a pUC19 backbone fragment. The three fragments were assembled by In-Fusion Snap Assembly (Takara) to generate the subcloning vectors pU6gRNA_RALF7_gRNA2 or pU6gRNA_RALF6/8_gRNA. The resulting gRNA cassettes, driven by the OsU6 promoter, was cloned into the pZH_gYSA_PubiMMCas9 vector after digestion with AscI and PacI, generating pZH_gYSA_PubiMMCas9-RALF7_gRNA2 and pZH_gYSA_PubiMMCas9-RALF6/8_gRNA, respectively. These vectors were introduced into *Agrobacterium tumefaciens* strain EHA101 and used for *Agrobacterium*-mediated transformation of the Nipponbare cultivar as described by Okuyama et al.(2). The mutation types in the regenerated T0 plants were confirmed by PCR using the primers listed in Table S6.

To examine the subcellular localization of OsRALF7-eGFP, the plasmid pPBV[Exp]-Hygro-{RALF7}>{RALF7}/3xGGGGS/EGFP was generated by VectorBuilder (https://en.vectorbuilder.com/; vector ID: VB231211-1141myr). This vector was introduced into *A. tumefaciens strain* EHA101 and used for *Agrobacterium*-mediated transformation of Nipponbare. Transgene insertion in the regenerated T0 plants was confirmed by PCR using the primers listed in Table S6 and by GFP fluorescence.

To visualize *M. oryzae*, the pBITEFpro:mCherry vector (3) was introduced into *A. tumefaciens* strain AGL1 and used for *Agrobacterium*-mediated transformation of *M. oryzae* strain Ao92-06-02, as previously described with minor modifications (4). The bacterial culture was diluted in induction medium (IM) to an optical density at 660 nm of 1.5–2.0. Fungal conidia were suspended in IM and adjusted to 1–5×$10^6$ conidia/mL. The bacterial suspension was mixed with the conidial suspension at a volume ratio of 1:10, and the mixture was spread onto filter paper placed on IM agar plates containing 200 μM acetosyringone. After co-cultivation at 25 °C for 3 days, the filter papers were transferred to defined complex medium (DCM) plates containing 0.17% yeast nitrogen base without amino acids, 0.2% asparagine, 0.1% ammonium nitrate, 1% glucose, and 1.5% agar, adjusted to pH 6.0 with $Na_2HPO_4$, and supplemented with 100 μg/mL chlorimuron-ethyl (prepared from 400 mg ATTRACTIVE; MARUWA Biochemical, per L DCM) and 50 μg/mL meropenem hydrate (Sumitomo Dainippon Pharma). After incubation at 25 °C for 2 days, the filter papers were removed. After an additional 2–3 days of incubation at 25 °C, individual transformants were transferred to fresh selection medium.

*M. oryzae* knockout mutants of *MSB2*, *SHO1*, *PTH11*, *CPKA*, and *PMK1* were generated using a homologous recombination-mediated method with the CRISPR–Cas9 system (5–7). pCRISPR/Cas9-PU6-FoNLSx2 was constructed from pCRISPR/Cas9-FoU6-FoNLSx2 (7) by replacing the *Fusarium oxysporum* U6 promoter with the *M. oryzae* U6-1 promoter (5). The target sequences were designed using the CRISPRdirect web tool (https://crispr.dbcls.jp/)(8). Each gRNA expression plasmid was generated by inserting the annealed oligonucleotides (Table S6) into pCRISPR/Cas9-PU6-FoNLSx2, as previously described (6). The resulting vectors were designated pcc-msb2, pcc-sho1, pcc-pth11, pcc-pka and pcc-pmk1.

Each donor vector was constructed by cloning the PCR-amplified flanking regions (ca. 1,000-bp) of target genes into pMK-dGFP (5). For the *CPKA* knockout, the 5′ and 3′ homology arms were cloned into the vector after digestion with SpeI/HindIII and SphI/XhoI, respectively. The resulting vector, pPka-donor, was used for double-crossover-mediated gene deletion (5). For the knockouts of *MSB2*, *SHO1*, *PTH11*, and *PMK1*, the 5′ and 3′ homology arms for each gene were fused by PCR and cloned after digestion with the appropriate restriction enzymes (Table S6). The resulting vectors–pMsb2-donor, pSho1-donor, pPth11-donor, and pPmk1-donor–were used for single-crossover-mediated gene deletion (6).

The corresponding CRISPR and donor vectors were simultaneously introduced into protoplasts of the *M. oryzae* P2 strain via the PEG-mediated transformation method (5, 6). Transformants were selected on YG agar medium supplemented with 100 µg/mL hygromycin B, and the knockout mutants were confirmed by PCR using the primers listed in Table S6.

All primers and plasmids used in this study are listed in Tables S6 and S7.

**Preparation of germinated conidia for cAMP, Pmk1 phosphorylation, and RNA-seq analyses**
Conidia prepared as described above were harvested and filtered once through Miracloth. For cAMP and MAPK analyses, conidia were resuspended in 25% CM at $5.0\times10^5$ conidia/mL, and 15 mL of the suspension was used for each biological replicate. For RNA-seq analysis, conidia were resuspended in 30% CM at $3.0\times10^5$ conidia/mL, and 10 mL of the suspension was used for each biological replicate. The indicated peptide was added to each suspension, which was then dispensed as 20-µL droplets onto glass coverslips and incubated in a humid chamber at 25°C. Conidial germination and appressorium formation were verified under a light microscope before sample collection.

For cAMP analysis, fungal material was scraped from the coverslips with a razor blade. The coverslips were rinsed with 1× PBS (pH 7.4), and the recovered material was pooled and maintained on ice. Samples were centrifuged at 20,000 × *g* for 20 min at 4°C, and the supernatant was removed to obtain a final volume of 500 µL. Samples were flash-frozen in liquid nitrogen and lyophilized for 16 h before cAMP extraction and analysis.

For MAPK analysis, the conidial material on coverslips was treated with protein extraction buffer containing 50 mM Tris-HCl (pH 7.5), 100 mM NaCl, 1% Triton X-100, 10% glycerol, 5 mM EDTA, 10 mM *β*-glycerophosphate, 1 mM $Na_3VO_4$, 50 mM NaF, 1 mM PMSF, and 1× protease inhibitor cocktail. Fungal material was then scraped from the coverslips with a razor blade, pooled, kept on ice, and centrifuged at 20,000 × *g* for 20 min at 4°C. The supernatant was removed, leaving approximately 400–500 µL of buffer. The pellet was supplemented with approximately 100 µL of prechilled glass beads (425–600 µm; Sigma-Aldrich) and one 3-mm zirconia bead (Bio Medical Science), and homogenized using a FastPrep-24 5G instrument (MP Biomedicals) at 4.0 m/s for 30 s. After cooling on ice for 5 min, homogenization was repeated for a total of three cycles. Lysates were centrifuged at 16,000 × *g* for 20 min at 4°C, and the supernatants were collected for protein quantification and immunoblot analysis.

For RNA-seq analysis, conidial material on coverslips was treated with a one-fifth dilution of RNAlater (Thermo Fisher Scientific) and scraped from the coverslips with a razor blade. Recovered material was pooled, maintained on ice, and centrifuged at 20,400 × *g* for 30 min at 4°C. The pellet was resuspended in prechilled water and washed twice, with centrifugation at 20,000 × *g* for 10 min at 4°C after each wash. The final pellet was stored at −20°C until RNA extraction. Frozen pellets were resuspended on ice in 500 µL of RA1 buffer supplemented with DTT, supplied with the NucleoSpin RNA kit (Takara). Each sample was supplemented with approximately 100 µL of prechilled glass beads (425–600 µm; Sigma-Aldrich) and one 3-mm zirconia bead (Bio Medical Science). Samples were homogenized using a FastPrep-24 5G instrument (MP Biomedicals) at 4.0 m/s for 30 s, cooled on ice for 5 min, and homogenized for a total of three cycles. RNA was immediately purified according to the manufacturer's instructions.

**Preparation of fungal mycelia for cAMP and Pmk1 phosphorylation analyses**

The *M. oryzae* P2 wild-type strain was grown on CM agar plates for 4–7 days. Agar plugs containing fungal mycelia were then transferred to 50% CM liquid medium and cultured with shaking at 130 rpm for 2 days at room temperature. The resulting mycelia were washed twice with 50% CM. Two 1 $cm^2$ agar plugs were transferred to each well of a 24-well plate containing 500 µL of 50% CM supplemented with 5 µM OsRALF7 or mock control. Plates were incubated with shaking at 100 rpm at room temperature. At the indicated time points, mycelia were harvested and immediately flash-frozen in liquid nitrogen. For cAMP analysis, samples were lyophilized for 16 h before extraction and measurement. For Pmk1 phosphorylation analysis, samples were ground to a fine powder for protein extraction, and suspended in protein extraction buffer containing 50 mM Tris-HCl (pH 7.5), 100 mM NaCl, 1% Triton X-100, 10% glycerol, 5 mM EDTA, 10 mM *β*-glycerophosphate, 1 mM $Na_3VO_4$, 50 mM NaF, 1 mM PMSF, and 1× protease inhibitor cocktail. The samples were centrifuged at 16,000 × *g* for 20 min at 4°C, and the supernatants were collected for immunoblot analysis.

**cAMP measurement**

Lyophilized fungal material was weighed to determine dry weight before extraction. Dried samples were ground to a fine powder in liquid nitrogen and resuspended in 200 µL of ice-cold 6% (w/v) trichloroacetic acid for 10 min on ice. After centrifugation at 4,000 rpm for 15 min at 4°C, supernatants were washed four times with five volumes of water-saturated diethyl ether. The remaining aqueous extracts were lyophilized, resuspended in 100 µL of Assay/Lysis buffer, and analyzed using the cAMP-Screen Direct Cyclic AMP Immunoassay System (Thermo Fisher Scientific) according to the manufacturer's instructions. Intracellular cAMP concentrations were calculated from a standard curve, normalized to fungal dry weight, and expressed as pmol/g dry weight.

**Immunoblot analysis of Pmk1 phosphorylation**

Protein concentrations were determined using the Rapid Gold BCA Protein Assay Kit (Thermo Fisher Scientific). A total of 20 µg of protein per sample was heated at 94°C for 1 min in SDS sample buffer, separated on a 12% SDS-polyacrylamide gel and transferred to a PVDF membrane. Membranes were blocked with 5% BSA in TBS-T (0.1 M Tris-HCl, pH 7.5, 0.15 M NaCl, and 0.1% Tween 20) for 1 h at room temperature and incubated overnight at 4°C with a phospho-p44/42 MAPK (Erk1/2) (Thr202/Tyr204) antibody (Cell Signaling Technology, 4370S; 1:2000) diluted in 5% BSA in TBS-T. After washing with TBS-T, membranes were incubated for 1 h at room temperature with an HRP-conjugated anti-rabbit IgG secondary antibody (Cytiva, NA934V, 1:5000) diluted in TBS-T. Signals were detected using SuperSignal West Dura Substrate (Thermo Fisher Scientific) and a FUSION Light-Capture System (Vilber).

**Immunoblot detection of OsRALF7-eGFP**

7–8 cm of Leaf-sheath segments were collected from fully expanded sixth leaves of five-week-old rice plants, frozen in liquid nitrogen, and ground to a fine powder. The powder was mixed with 3× sample buffer and heated at 95°C for 5 min. Lysates (10 µL per sample) were separated by SDS–PAGE and transferred to PVDF membranes. Membranes were blocked with 5% nonfat dry milk in TBS-T for 1 h at room temperature and incubated overnight at 4°C with a GFP antibody B-2 (Santa Cruz Biotechnology, sc-9996, 1:2000) diluted in 5% nonfat dry milk in TBS-T. After washing with TBS-T, membranes were incubated for 1 h at room temperature with an HRP-conjugated anti-mouse IgG secondary antibody (Amersham Biosciences, NA931-1ML, 1:4000) diluted in TBS-T. Signals were detected using SuperSignal West Dura Substrate and a FUSION Light-Capture System. Band intensities were quantified using ImageJ.

**Measurement of leaf-surface hydrophobicity**

Leaf-surface hydrophobicity was assessed in wild-type and *Osralf7* mutant plants by measuring the contact angle of a 9-µL water droplet on the adaxial surface using a PMTA wettability measurement system (Meiwafosis, Tokyo, Japan). Measurements were obtained from a 1-cm-long region located 5–6 cm from the tip of the fully expanded fifth leaf of five-week-old plants. For each sample, the contact angle was measured three times within 5 s of droplet deposition, and the mean value was used for analysis.

#### Scanning electron microscopy of leaf surfaces

Surface structures on the adaxial leaf surface were visualized using a scanning electron microscope. A 1-cm-long segment was excised from a region 5–6 cm from the tip of the fully expanded sixth leaf of six-week-old plants. The adaxial surface of each leaf segment was observed using a Miniscope TM3000 scanning electron microscope (Hitachi High-Technologies, Tokyo, Japan) operated at 15 kV.

#### Leaf cuticular wax composition analysis by GC-MS

Leaf cuticular waxes were extracted with minor modifications of published methods (9, 10). Whole leaf blades were excised from fully expanded sixth leaves of five-week-old rice plants, and approximately 7.5 cm of each blade was immersed in chloroform for 30 s at 60°C. 10 µg of tetracosane was added to each extract as an internal standard. The extracts were evaporated under a nitrogen stream at approximately 70°C. The extracted leaf regions were scanned to determine the extracted leaf area, and wax amounts were normalized to leaf area. The dried extracts were derivatized by adding 100 µL pyridine and 100 µL N,O-bis(trimethylsilyl)trifluoroacetamide (BSTFA). Samples were incubated at 70°C with shaking at 130 rpm for 40 min and then cooled to room temperature.

Wax components were identified by GC–MS using an Orbitrap$^{TM}$ Exploris$^{TM}$ GC MS system (Thermo Fisher Scientific) equipped with a TG-5MS column (30 m × 0.25 mm i.d., 0.25-µm film thickness; Thermo Fisher Scientific). Helium was used as the carrier gas at 1 ml $min^{-1}$. The injection port was maintained at 320°C. The column temperature was held at 50 °C for 2 min, increased to 200°C at 40 °C $min^{-1}$, held for 2 min, then increased to 320 °C at 3 °C $min^{-1}$ and held for 40 min. Wax components were identified by comparing their mass spectra with entries in the NIST library and by evaluating their retention times. An alkane standard mixture (C8–C40) was analyzed by GC–MS to determine retention (Kovats) indices. Wax quantities were determined by comparing total ion chromatogram (TIC) peak areas with those of the internal standard (tetracosane) and were expressed per unit extracted leaf-surface area.

#### Immunogold labeling and transmission electron microscopy

Leaf-sheath samples were collected from OsRALF7-eGFP-expressing rice plants and nontransgenic wild-type plants used as a negative control. Samples were rapidly frozen by immersion in liquid propane at approximately −175°C. Freeze substitution was performed in ethanol containing 2% tannic acid and 2% distilled water at −80°C for 48 h. Samples were subsequently warmed to −20°C for 3 h and then to 4°C for 1 h. After freeze substitution, samples were washed three times for 30 min each and then overnight in anhydrous ethanol at room temperature. Samples were infiltrated at 4°C three times for 30 min each with a 1:1 mixture of anhydrous ethanol and LR White resin (London Resin Co., Berkshire, UK), followed by three 30-min incubations in LR White resin. Samples were transferred to fresh LR White resin and polymerized overnight under ultraviolet light. Ultrathin sections (80 nm) were prepared using an Ultracut UCT ultramicrotome (Leica, Vienna, Austria) equipped with a diamond knife and mounted on nickel grids. Sections were incubated overnight at 4°C with a rabbit polyclonal anti-GFP antibody (abcam, sb6556). After three 1-min washes in PBS containing 1% BSA, sections were incubated for 2 h at room temperature with a goat anti-rabbit IgG antibody conjugated to 10-nm colloidal gold particles (BBinternational, Golden Gate, Cardiff, UK, EM.GAR10). Sections were then washed three times for 1 min each in PBS and fixed with 2% glutaraldehyde in 0.1 M phosphate buffer for 3 min at room temperature. Sections were stained with 2% uranyl acetate for 5 min and Lead Stain Solution (Sigma-Aldrich) for 3 min at room temperature. Samples were examined using a JEM-1400Plus transmission electron microscope (JEOL, Tokyo, Japan) operated at 100 kV. Images were acquired using an EM-14830RUBY2 CCD camera (JEOL) at a resolution of 3,296×2,472 pixels.

#### RNA isolation and RNA-seq analysis of rice suspension cells

For RNA-seq analysis of peptide- or chitin-treated rice suspension cells, cells were cultured in R2S medium for 4 days at 25–30 °C. Approximately 150 mg of cells were transferred to each well of a

12-well plate containing 2 mL of fresh R2S medium and incubated overnight in the dark at 25–30 °C. OsRALF7 peptide was added to a final concentration of 1 μM, and chitin was added to a final concentration of 2 μg/mL. 1× PBS was used as the mock control. After 1 h of treatment, the cells were harvested, immediately frozen in liquid nitrogen, and ground to a fine powder for RNA extraction. Total RNA was isolated using TRIzol reagent (Thermo Fisher Scientific) according to the manufacturer's instructions, followed by Ambion DNase I treatment (Thermo Fisher Scientific) to remove contaminating genomic DNA. RNA was further purified using a Direct-zol RNA MiniPrep kit (Zymo Research). RNA integrity was assessed using an Agilent 2100 Bioanalyzer, and only samples with an RNA integrity number (RIN) greater than 8.5 were used for library preparation. RNA concentration was measured using a Qubit RNA BR Assay Kit (Thermo Fisher Scientific). Sequencing libraries were prepared using the NEBNext Poly(A) mRNA Magnetic Isolation Module (NEB), NEBNext Ultra II Directional RNA Library Prep Kit for Illumina (NEB), and NEBNext Multiplex Oligos for Illumina (NEB) according to the manufacturers' instructions. Libraries were pooled and sequenced in paired-end 100-bp mode on a NovaSeq platform at the National Institute of Genetics.

**RNA-seq data processing of rice suspension cells**

The raw FASTQ files obtained from the RNA-seq runs described in the above section were first subjected to quality checks and filtering using fastp (v. 0.23.4) with the flags `--cut_front --cut_tail --cut_right --correction` (11). We primarily employed the read aligner STAR (v. 2.7.11b) for read alignment (12), whereas Bawtie2 (v. 2.5.4) was used for reads that STAR failed to align (13). Index files for read alignment were generated based on the Nipponbare reference genome (IRGSP-1.0) and reference transcripts with the settings `--runMode genomeGenerate --genomeSAindexNbases 13 --sjdbOverhand 149` and the default settings for STAR and Bawtie2, respectively. Exon-intron structure information of the reference transcripts stored as a GTF file was also supplemented to STAR through the sjdbGTFfile argument. Since the Nipponbare genome has two different sets of gene models in RAP-DB and the MSU database (14), both datasets in GFF format were downloaded from the databases (https://rapdb.dna.affrc.go.jp/ and https://rice.uga.edu/). The gene models in the MSU database were merged with those in RAP-DB, if they were absent in RAP-DB. Transcript sequences and exon-intron structures were obtained based on the merged dataset of the gene models and prepared as reference transcripts, as described above. The quality-filtered clean reads were aligned to the Nipponabre genome using STAR with the settings `--outSAMtype BAM SortedByCoordinate --quantMode TranscriptomeSAM --outSAMunmapped Within KeepPairs --outFilterMultimapNmax 1000`. The unmapped reads were then extracted from the output BAM files using the `view` module of samtools (v. 1.19.2) with the settings `-h -f 13` (15). These unmapped reads were subjected to further read alignment using Bawtie2 with the default settings, which aligns reads to transcript sequences rather than genome sequences. The output read alignments generated by STAR and Bawtie2 were merged into one BAM file for each sample, respectively, using the `merge` module of samtools. The merged BAM files were subjected to quasi-mapping using the read quantifier Salmon (16). As input BAM files store read alignments to transcript sequences rather than genome sequences, a FASTA file of the Nipponbare transcript sequences was supplied to Salmon via the argument `-t`, leaving all other settings at their default values. The resultant read quantification data were then analyzed using R (17) on a pipeline consisting of an array of R packages as described below.

**Differential expression analysis**

The quantification data generated by Salmon were imported into an R environment using the tximport function of the tximport package with the default settings (18). Differential expression gene (DEG) analysis was conducted using the DESeq2 package (19), with the regression model 'read counts ~ condition', where each combination of genotype and treatment was modeled as a unique level of the condition factor. As described in the section above, we had three conditions in total: wildtype under mock treatment (WT-Mock), wildtype under Chitin treatment (WT-Chitin), and wildtype under RALF7 treatment (WT-RALF7). The Wald test was performed to evaluate the statistical significance of the condition on the changes in the expression profile. Independent Hypothesis Weighting (IHW), as implemented in the IHW package (20), was employed for p-value adjustment, as instructed in the official vignette of the DESeq2 package (https://doi.org/doi:10.18129/B9.bioc.DESeq2). DEGs were identified using thresholds of an absolute log2-transformed fold change (|log2FC|) ≥ 1 and an adjusted p-value (Padj) < 0.05. A Venn diagram was drawn to visualize the subset using the VennDiagram package in R (21).

**Gene set enrichment analysis**

Significantly overrepresented functional and biological terms were analyzed in the RALF7-induced DEGs. The culsterprofiler package was used for the statistical evaluation of enriched terms (22). GO terms and the KEGG pathway terms were assigned to the Nipponbare genes using interProScan (v. 5.65-97.0) (23) and eggNOG-mapper v2 (v. 2.1.12) (24, 25). The hierarchical structure of the GO terms was obtained from the Gene Ontology knowledgebase (https://geneontology.org/docs/download-ontology). Enriched terms were visualized using the ggplot2 package in R (26). Since GO terms have a hierarchical structure, enriched terms were visualized separately for terms at the same level in each term hierarchy. To provide a clearer overview of the enriched terms, GO terms were visualized together for each GO category, namely Biological Process, Molecular Function, and Cellular Component, using an in-house script that generated an interactive network graph. The in-house script internally uses the igraph and htmlwidgets packages for visualization (27, 28).

**RNA isolation, sequencing and transcriptome analysis of *M. oryzae***

To evaluate the transcriptional profile during appressorium formation under peptide treatment, conidia produced as described above were harvested, filtered once through Miracloth, and resuspended to $3.0 \times 10^5$ conidia/mL in 30% CM liquid medium containing the indicated peptide concentrations. A total of 10 mL of each conidial suspension was dispensed onto glass coverslips as 20 µL droplets and incubated in a moist chamber at 25 °C. Conidial germination and appressorium formation were confirmed prior to sampling.

For sample collection, coverslips were treated with a one-fifth dilution of RNAlater (Thermo Fisher Scientific), and fungal material was scraped off with a razor blade. Recovered material was pooled, kept on ice, and centrifuged at 20,400×g for 30 min at 4 °C. The pellet was resuspended and washed twice with prechilled water, followed each time by centrifugation at 20,000×g for 10 min at 4 °C. The final pellet was stored at −20 °C until RNA extraction. Frozen cell pellets were resuspended on ice in 500 µL of RA1 buffer containing DTT from the NucleoSpin RNA kit (Takara). Each sample was supplemented with approximately 100 µL of pre-chilled glass beads (425–600 µm; Sigma-Aldrich) and one 3-mm zirconia bead (Bio Medical Science). Samples were homogenized using a FastPrep-24 5G instrument (MP Biomedicals) at 4.0 m/s for 30 s, cooled on ice for 5 min, and subjected to a total of three cycles. RNA was then purified immediately according to the manufacturer's instructions.

Libraries were prepared from the extracted RNA using the NEBNext Ultra II Directional RNA Library Prep Kit for Illumina. The resulting libraries were sequenced on the Illumina NovaSeq platform. The obtained sequencing reads were filtered using FaQCs (29) and then mapped to the reference genome of the rice blast fungus strain 70-15 using HISAT2 (30). The 70-15 genome sequence was obtained from Ensembl Fungi (https://ftp.ensemblgenomes.ebi.ac.uk/pub/fungi/release-60/fasta/magnaporthe_oryzae/dna/). Transcript abundance was subsequently quantified as transcripts per million (TPM) using TPMCalculator (31).

**Heatmap visualization of transcriptomic data**

Heatmaps of gene expression changes in rice suspension cells and *M. oryzae* were generated in R using the Heatmap function from the ComplexHeatmap package (32, 33) based on $\log_2$ fold-change values calculated from TPM values relative to the mock treatment. Color scales were defined using the colorRamp2 function (32) from the circlize package.

**Evolutionary distance estimation**

Amino acid sequences of mature RALF peptides from *A. thaliana* and *O. sativa* were aligned using the MUSCLE algorithm implemented in the msa package (v. 1.39.5) in R (35). A distance matrix was calculated from the alignment using the dist.ml function with the WAG substitution model, after converting the alignment format using the as.phyDat function of the phangorn package (36). To identify the closest AtRALF peptide to OsRALF7, distances from OsRALF7 to all other peptides were extracted from the distance matrix.

**Accession Numbers**

The accession numbers for the sequences analyzed in this study are listed in *SI Appendix*, Table S8.

**Figures**

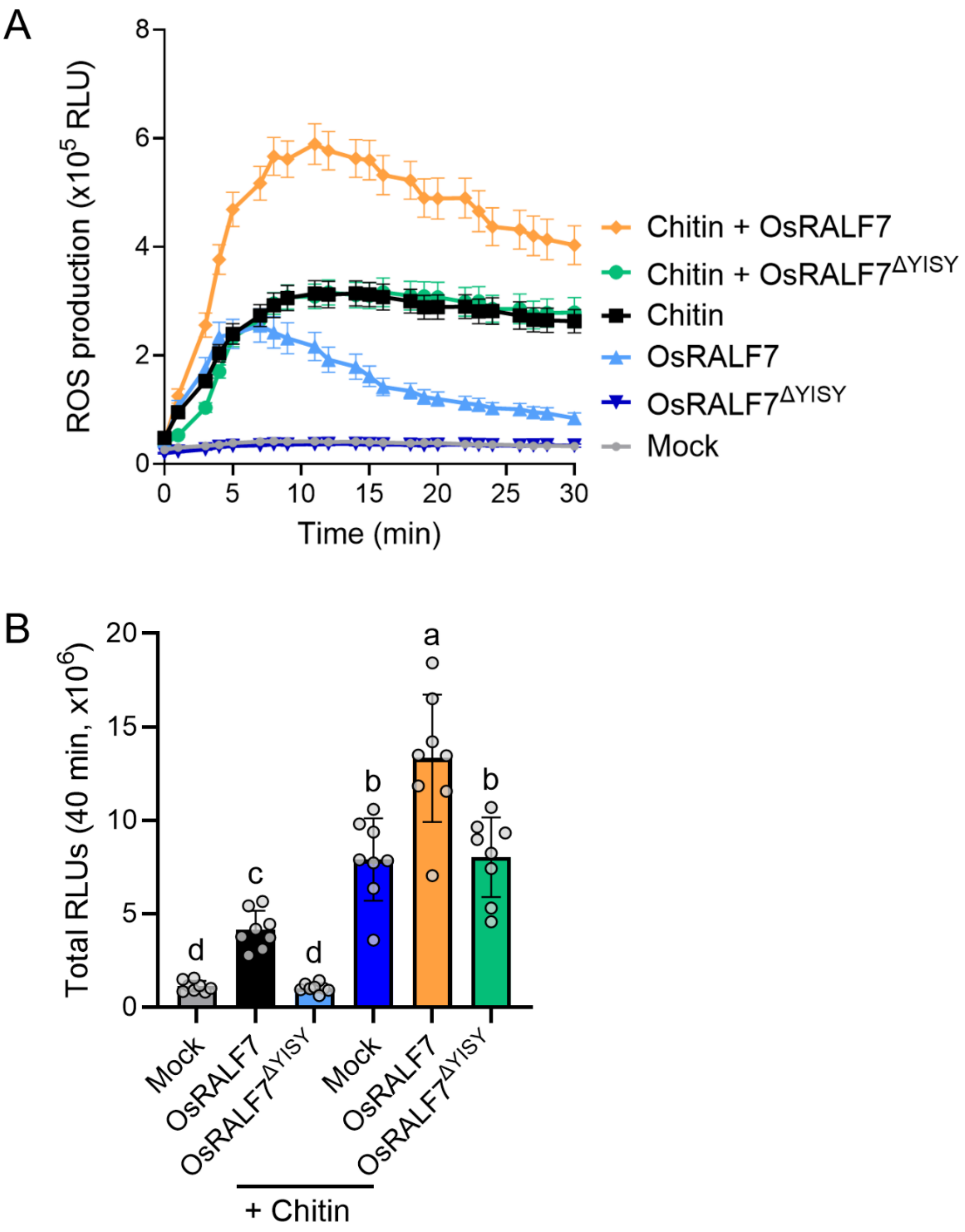


**Fig. S1.** OsRALF7 triggers chitin-enhanced ROS production.

(A) Time course of ROS production after treatment with OsRALF7 or OsRALF7$^{\Delta YISY}$ in the presence or absence of chitin. Wild-type Nipponbare cell cultures were treated with the synthetic OsRALF7 (1 µM) or OsRALF7$^{\Delta YISY}$ (1 µM) in the presence or absence of chitin (1 µg/mL). Luminescence was recorded every 1.5 min over 40 min. Data represented as the mean ± SEM from eight independently cultured flasks ($n$ = 8). Experiments were performed three times, each time with similar results.

(B) Total ROS production over 40 min, expressed as relative light units (RLUs). Each circle represents one independently cultured flask ($n$ = 8). Statistical significance of differences was assessed by one-way ANOVA followed by Tukey's multiple comparison test. Different letters indicate significant differences at $P < 0.05$.

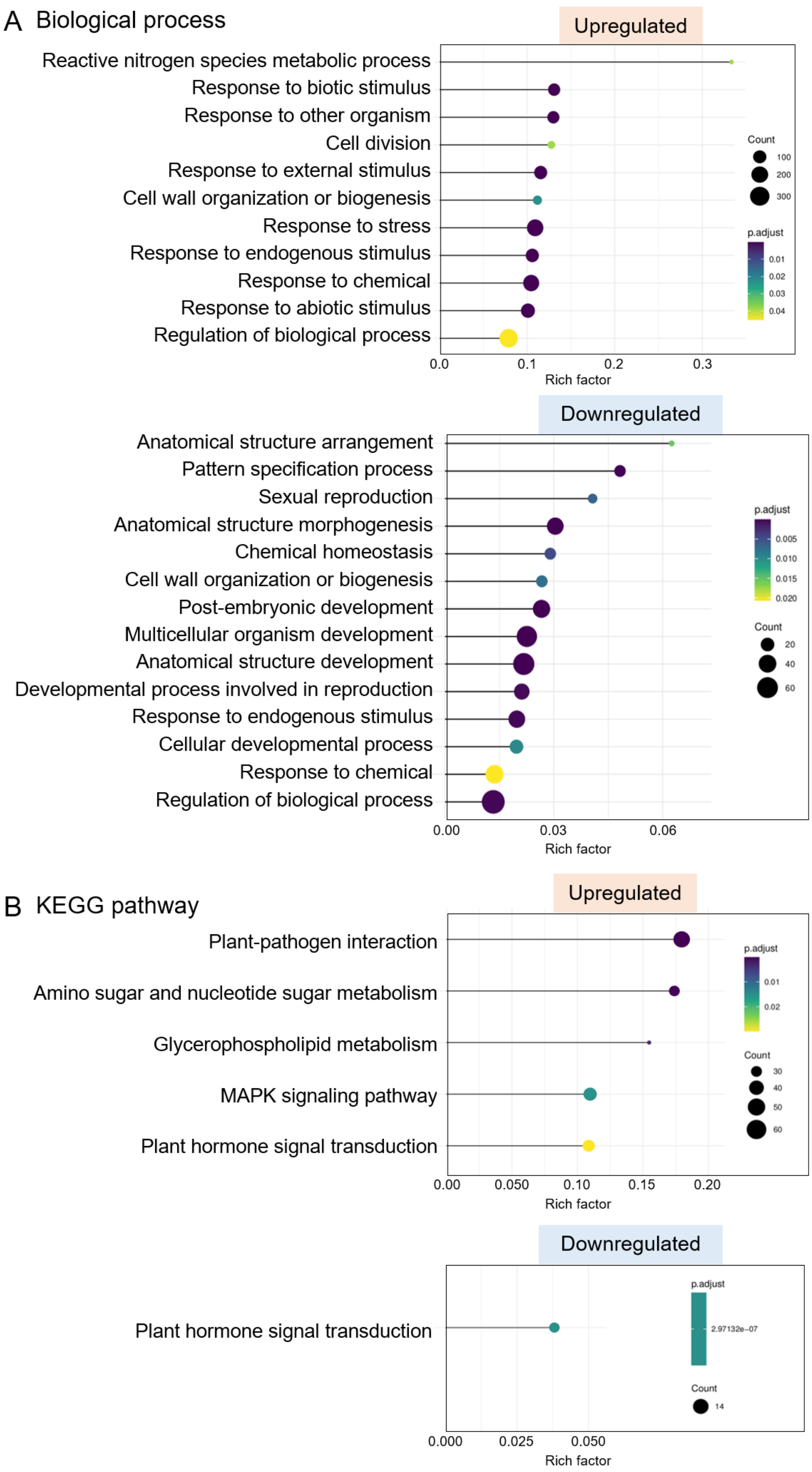

A Biological process
Upregulated
Reactive nitrogen species metabolic process
Response to biotic stimulus
Response to other organism
Cell division
Response to external stimulus
Cell wall organization or biogenesis
Response to stress
Response to endogenous stimulus
Response to chemical
Response to abiotic stimulus
Regulation of biological process
Count
100
200
300
p.adjust
0.01
0.02
0.03
0.04
0.0
0.1
0.2
0.3
Rich factor
Downregulated
Anatomical structure arrangement
Pattern specification process
Sexual reproduction
Anatomical structure morphogenesis
Chemical homeostasis
Cell wall organization or biogenesis
Post-embryonic development
Multicellular organism development
Anatomical structure development
Developmental process involved in reproduction
Response to endogenous stimulus
Cellular developmental process
Response to chemical
Regulation of biological process
p.adjust
0.005
0.010
0.015
0.020
Count
20
40
60
0.00
0.03
0.06
Rich factor
B KEGG pathway
Upregulated
Plant-pathogen interaction
Amino sugar and nucleotide sugar metabolism
Glycerophospholipid metabolism
MAPK signaling pathway
Plant hormone signal transduction
p.adjust
0.01
0.02
Count
30
40
50
60
0.00
0.050
0.10
0.15
0.20
Rich factor
Downregulated
Plant hormone signal transduction
p.adjust
2.97132e-07
Count
14
0.000
0.025
0.050
Rich factor

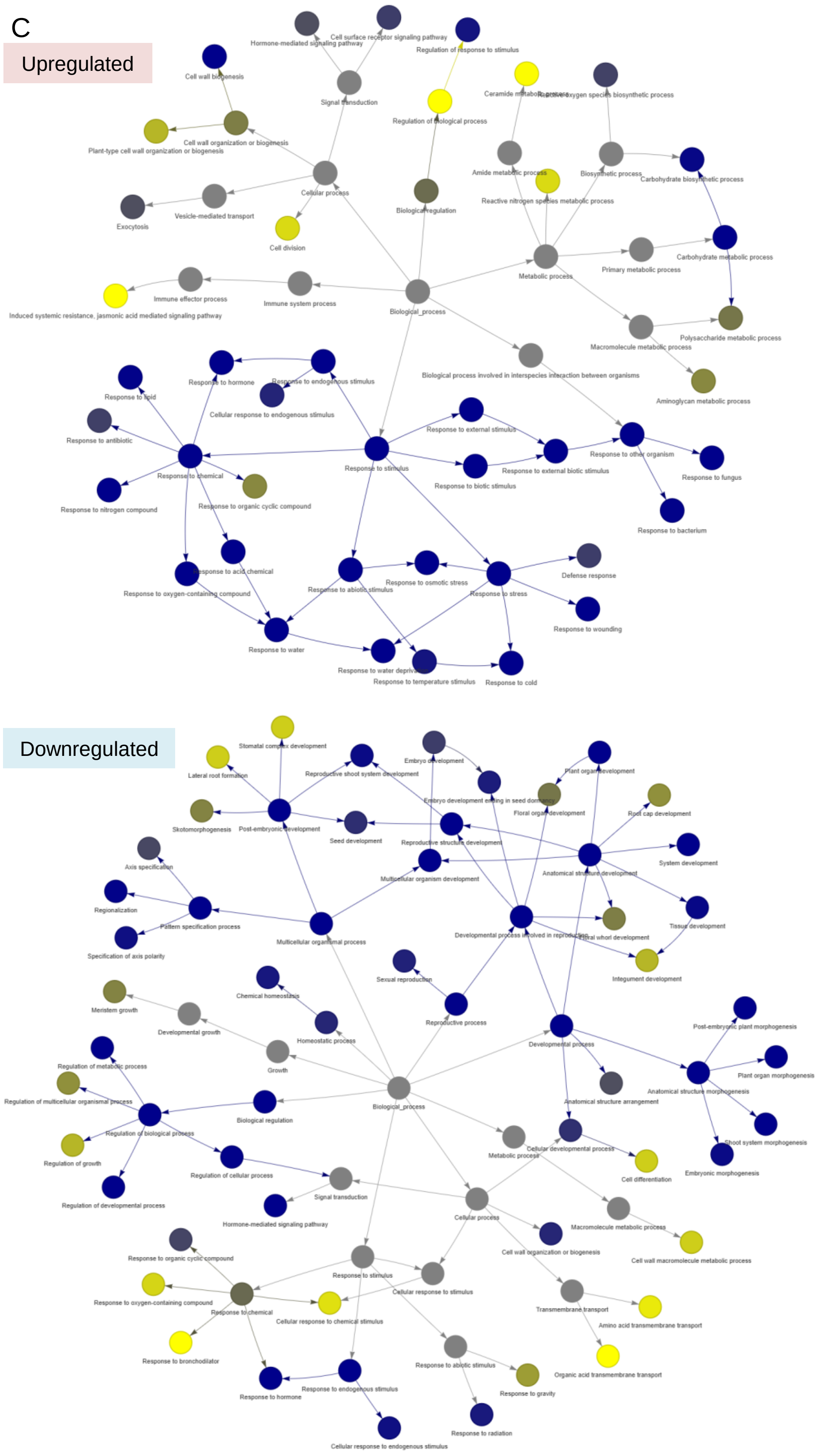
C
Upregulated
Hormone-mediated signaling pathway
Cell surface receptor signaling pathway
Regulation of response to stimulus
Cell wall biogenesis
Signal transduction
Ceramide metabolic process
Reactive oxygen species biosynthetic process
Regulation of biological process
Plant-type cell wall organization or biogenesis
Cell wall organization or biogenesis
Amide metabolic process
Biosynthetic process
Carbohydrate biosynthetic process
Exocytosis
Vesicle-mediated transport
Cellular process
Biological regulation
Reactive nitrogen species metabolic process
Cell division
Metabolic process
Primary metabolic process
Carbohydrate metabolic process
Immune effector process
Immune system process
Induced systemic resistance, jasmonic acid mediated signaling pathway
Biological_process
Macromolecule metabolic process
Polysaccharide metabolic process
Biological process involved in interspecies interaction between organisms
Aminoglycan metabolic process
Response to lipid
Response to hormone
Response to endogenous stimulus
Cellular response to endogenous stimulus
Response to antibiotic
Response to external stimulus
Response to stimulus
Response to chemical
Response to external biotic stimulus
Response to other organism
Response to fungus
Response to biotic stimulus
Response to nitrogen compound
Response to organic cyclic compound
Response to bacterium
Response to acid chemical
Response to oxygen-containing compound
Response to osmotic stress
Response to abiotic stimulus
Response to stress
Defense response
Response to wounding
Response to water
Response to water deprivation
Response to temperature stimulus
Response to cold
Downregulated
Stomatal complex development
Lateral root formation
Reproductive shoot system development
Embryo development
Plant organ development
Embryo development ending in seed dormancy
Floral organ development
Root cap development
Skotomorphogenesis
Post-embryonic development
Seed development
Reproductive structure development
System development
Axis specification
Multicellular organism development
Anatomical structure development
Regionalization
Pattern specification process
Tissue development
Specification of axis polarity
Multicellular organismal process
Developmental process involved in reproduction
Floral whorl development
Integument development
Sexual reproduction
Chemical homeostasis
Meristem growth
Developmental growth
Homeostatic process
Reproductive process
Post-embryonic plant morphogenesis
Developmental process
Growth
Plant organ morphogenesis
Regulation of metabolic process
Biological_process
Anatomical structure morphogenesis
Regulation of multicellular organismal process
Anatomical structure arrangement
Biological regulation
Regulation of biological process
Shoot system morphogenesis
Metabolic process
Cellular developmental process
Regulation of growth
Cell differentiation
Embryonic morphogenesis
Regulation of cellular process
Signal transduction
Regulation of developmental process
Hormone-mediated signaling pathway
Cellular process
Macromolecule metabolic process
Response to organic cyclic compound
Cell wall organization or biogenesis
Cell wall macromolecule metabolic process
Response to stimulus
Cellular response to stimulus
Response to oxygen-containing compound
Response to chemical
Cellular response to chemical stimulus
Transmembrane transport
Amino acid transmembrane transport
Response to bronchodilator
Organic acid transmembrane transport
Response to abiotic stimulus
Response to gravity
Response to endogenous stimulus
Response to hormone
Response to radiation
Cellular response to endogenous stimulus

**Fig. S2.** Transcriptomic profiling of rice cells treated with OsRALF7.

(A) Gene Ontology (GO) term enrichment analysis for Biological Process terms among genes upregulated or downregulated by OsRALF7 relative to water only (mock) control. Only GO terms at the third layer of the GO term hierarchy are shown.
(B) KEGG pathway enrichment analysis for genes upregulated or downregulated by OsRALF7 relative to mock. (A, B) In the enrichment dot plots, dot color indicates the adjusted *P* value and dot size indicates the number of genes assigned to each GO term or KEGG pathway.
(C) Enriched GO terms in the Biological Process category among genes upregulated or downregulated by OsRALF7 relative to mock were visualized as network graphs. Node color represents the FDR (yellow to dark blue).

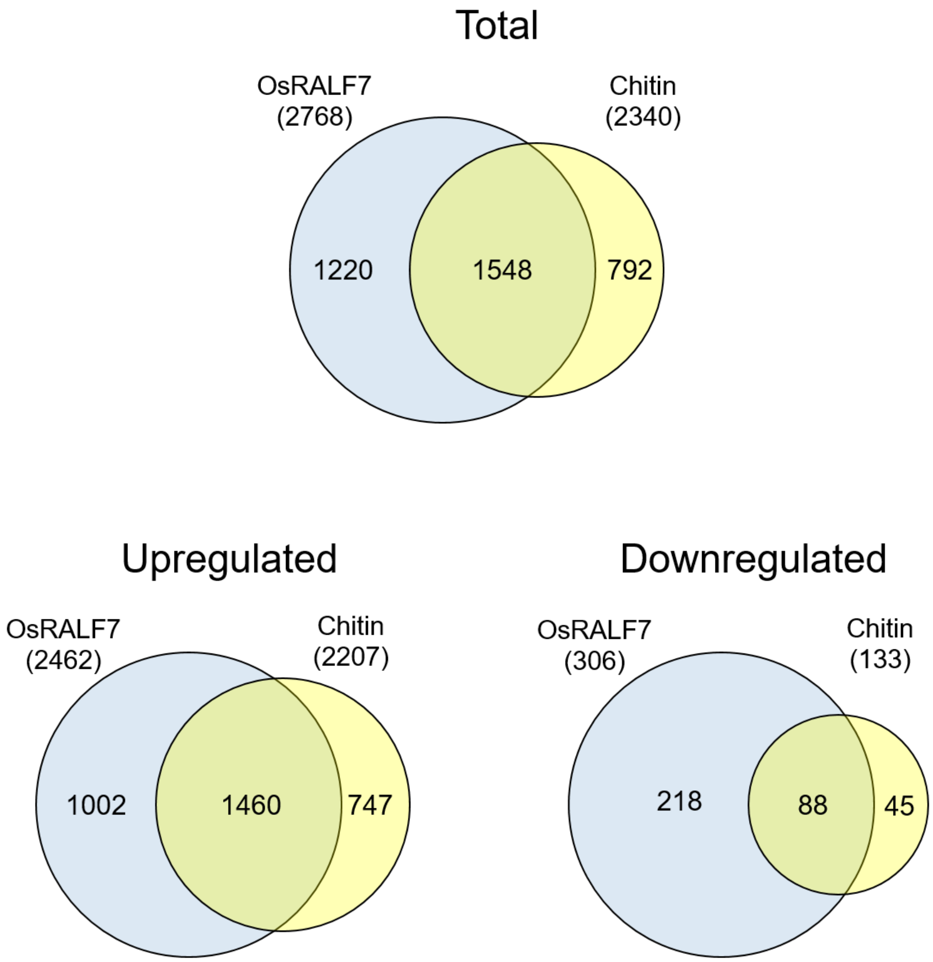


**Fig. S3.** Venn diagram of differentially expressed genes in rice suspension cells after OsRALF7 or chitin treatment. Venn diagrams show the overlap between genes differentially regulated by OsRALF7 or chitin treatment relative to water-treated (mock) cells. Blue and yellow circles indicate the sets of differentially expressed genes identified after OsRALF7 and chitin treatment, respectively. Numbers in parentheses indicate the total number of genes differentially expressed after each treatment. Numbers within the Venn diagram indicate genes unique to each treatment or shared between treatments. All overlaps were significant by hypergeometric test ($P$ < 0.001). Differentially expressed genes were defined as those with $|\log_2 FC| \geq 1$ and adjusted $P$ < 0.05, as listed in Dataset S1.

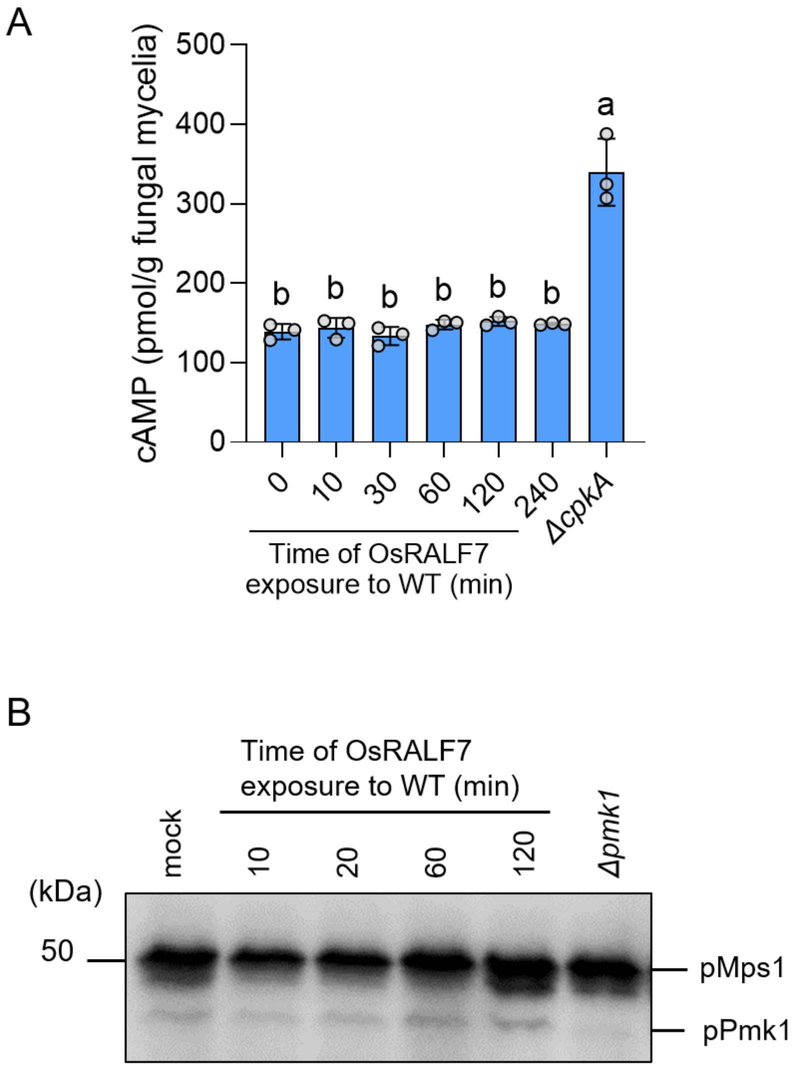


**Fig. S4.** Assessment of intracellular cAMP levels and Pmk1 phosphorylation in vegetative mycelia. (A) Intracellular cAMP levels in vegetative mycelia. *M. oryzae* mycelia were incubated in liquid 50% CM medium containing water (mock) or OsRALF7 (5 μM) for the indicated times, followed by cAMP measurement. Each circle represents an independently cultured flask ($n$ = 3). Error bars indicate SD. Statistical significance was assessed by one-way ANOVA followed by Tukey's multiple-comparisons test. Different letters indicate significant differences at $P < 0.05$. (B) Immunoblot analysis of phosphorylated Pmk1 in vegetative mycelia. *M. oryzae* mycelia were incubated in liquid 50% CM medium containing water (mock) or OsRALF7 (5 μM) for the indicated times before total protein extraction. Phosphorylated Pmk1 was detected using an anti-phospho-p44/42 MAPK antibody.

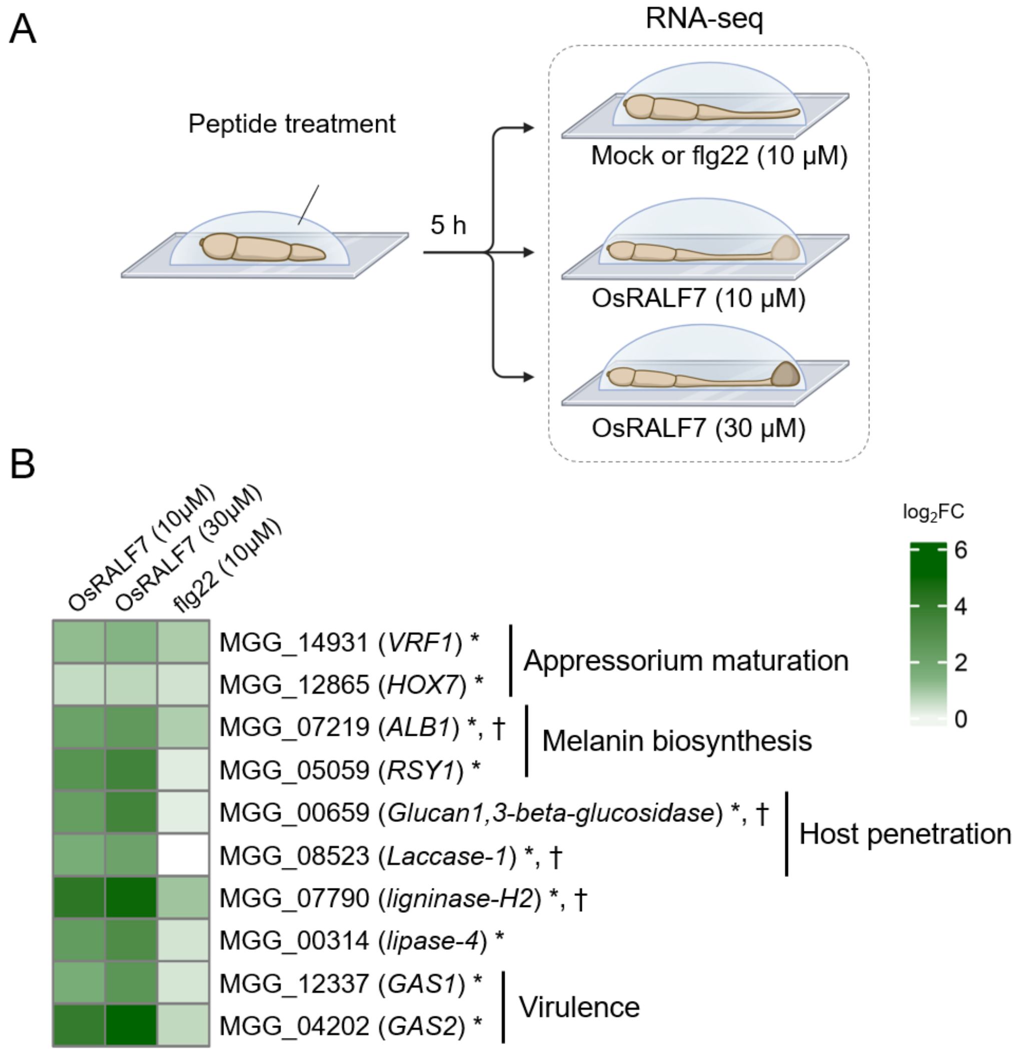


**Fig. S5.** OsRALF7 induces the expression of appressorium-associated genes.

(*A*) Diagram illustrating the RNA-seq experimental design. *M. oryzae* conidia were incubated in 30% CM medium with water only (Mock), OsRALF7 (10 μM or 30 μM), or flg22 (10 μM) for 5 h, followed by total RNA extraction for RNA-seq analysis.

(*B*) Heatmap representation of relative transcript levels for the indicated genes from *M. oryzae* treated with OsRALF7 or flg22. Color scale indicates $\log_2$ fold-change values relative to Mock treatment. *, down-regulated in the *Δpmk1* mutant (37); †, up-regulated by cAMP treatment (38, 39).

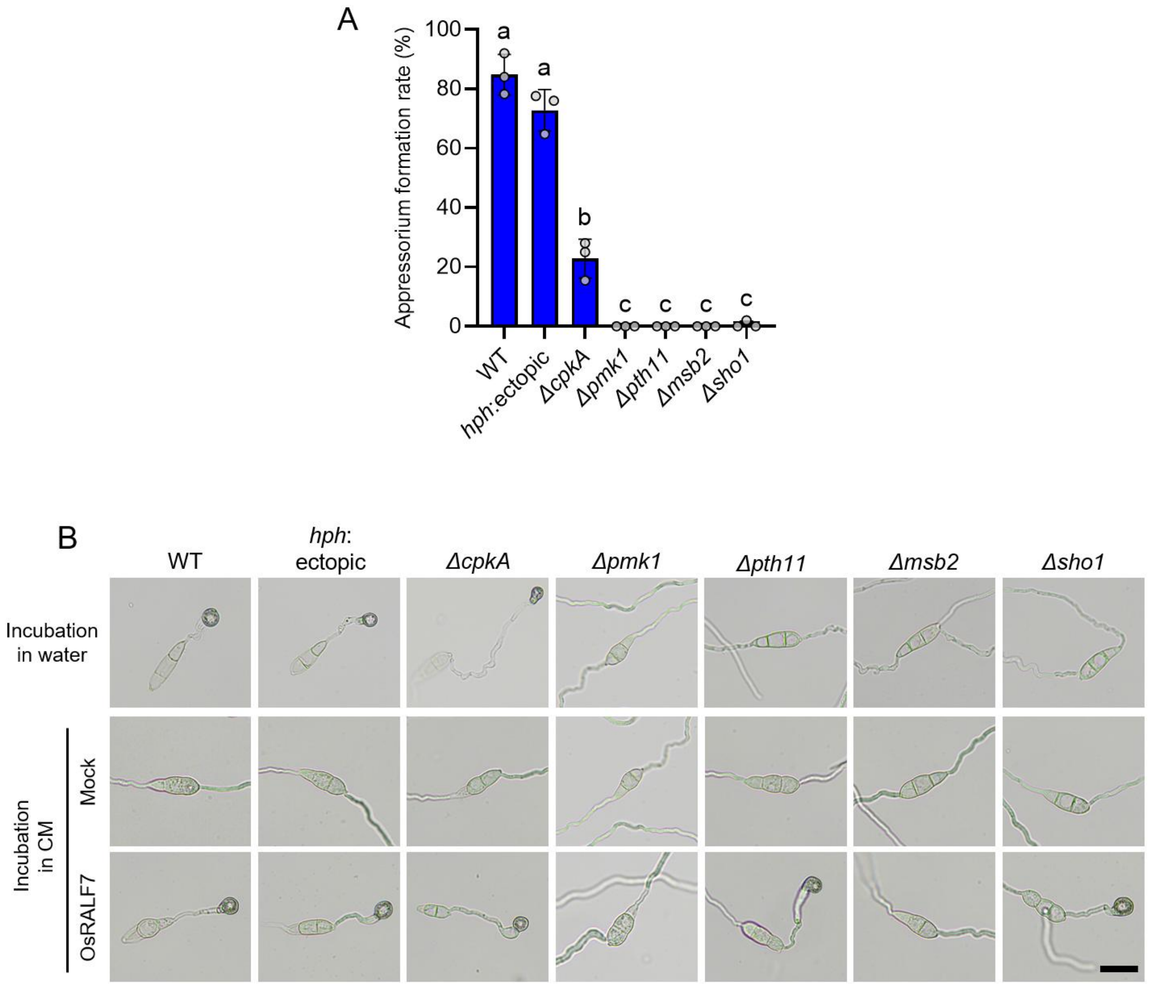


**Fig. S6.** Appressorium formation in *ΔcpkA*, *Δpmk1*, and surface-sensor mutants.

(A) Appressorium formation rates of *M. oryzae ΔcpkA*, *Δpmk1*, and surface-sensor mutant conidia incubated in water on hydrophobic coverslips for 24 h. Each circle represents an independently cultured plate ($n$ = 3); more than 50 conidia were scored per plate. Statistical significance was assessed by one-way ANOVA followed by Tukey's multiple-comparisons test. Different letters indicate significant differences at $P < 0.05$. Results are representative of three independent experiments.

(B) Representative micrographs of the *ΔcpkA*, *Δpmk1*, and surface-sensor mutant conidia incubated on hydrophobic coverslips in water or in 25% CM medium containing water (mock) or 5 μM OsRALF7 for 24 h. Images from the CM medium conditions correspond to the quantification shown in Fig. 3E. Scale bar, 20 μm.

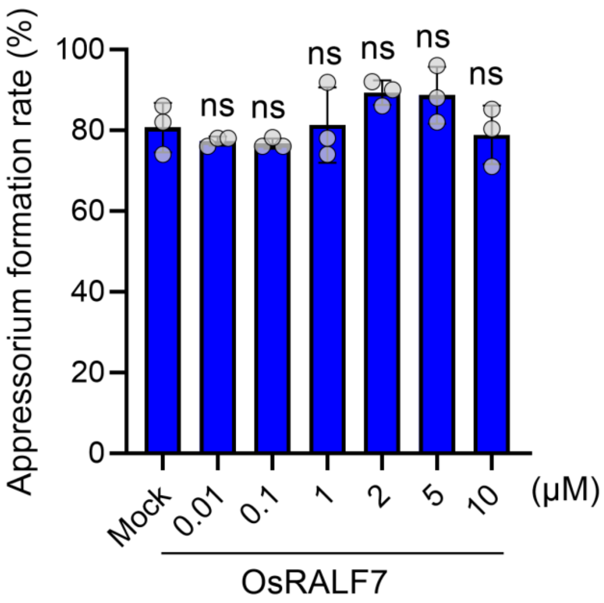


**Fig. S7.** Effects of OsRALF7 treatment on *M. oryzae* under water incubation.

Appressorium formation rates of *M. oryzae* conidia incubated in water with the indicated concentrations of OsRALF7 peptide for 8 h. Appressorium formation included non-melanized, melanized, and double appressoria; conidia showing only germ tube elongation were excluded from the appressorium-formed category. Each circle represents one independently cultured plate ($n$ = 3), with >50 conidia scored per plate. Statistical significance of differences was assessed by one-way ANOVA followed by Dunnett's multiple-comparisons test relative to water only (mock) control. ns, not significant. Experiments were performed three times, each time with similar results.

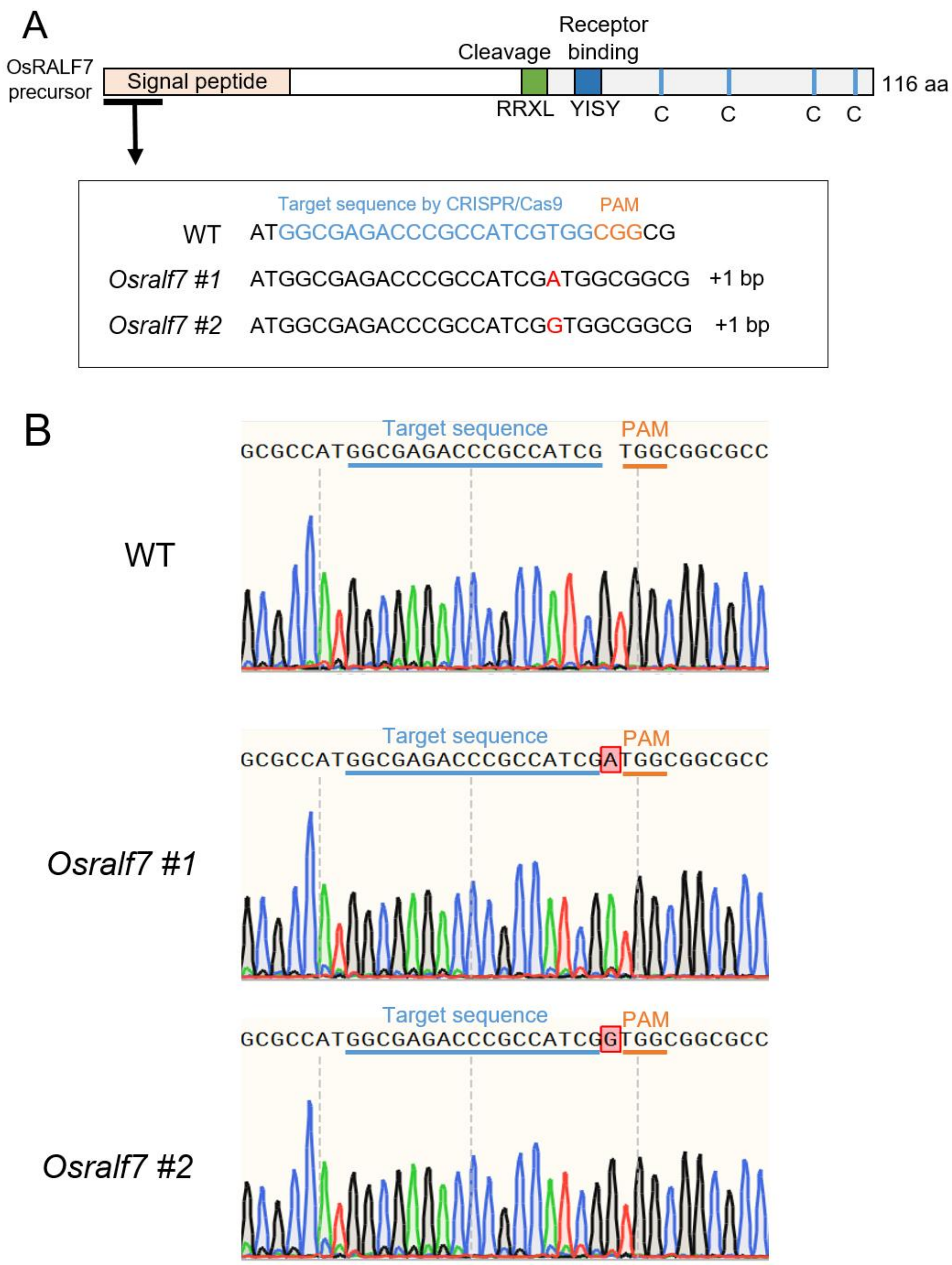


**Fig. S8.** Generation of *Osralf7* knockout mutants.

(A) Schematic representation of the *OsRALF7* precursor gene and the CRISPR–Cas9 target sites. The gRNA target sequence is shown in light blue and the PAM sequence in orange. Two independent *Osralf7* mutant alleles carry a 1-bp insertion at the target site, resulting in a frameshift and premature translational termination.

(B) DNA sequences of the *Osralf7* mutant alleles. The target sequence is underlined in light blue, and the PAM sequence is underlined in orange.

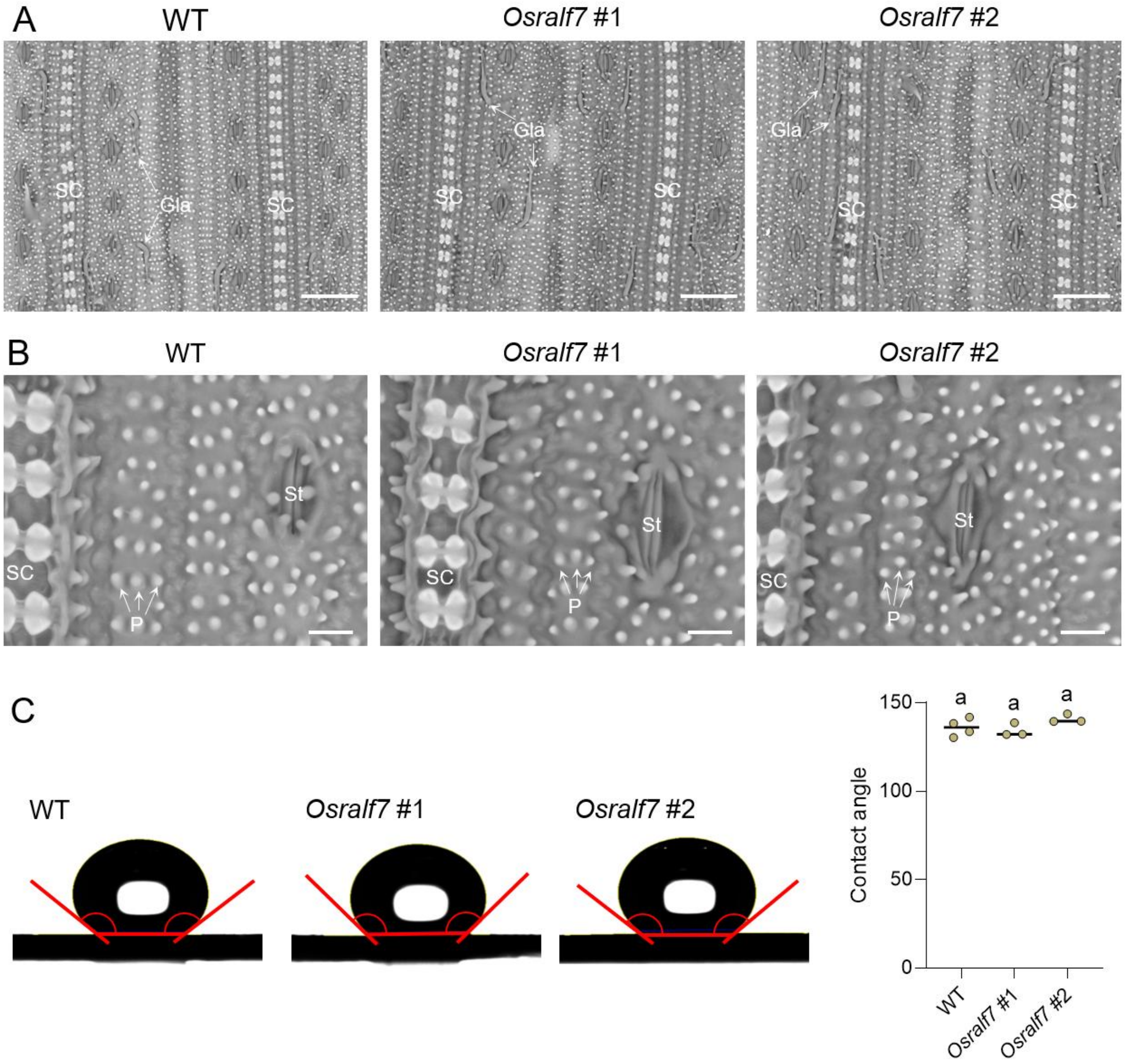


**Fig. S9**. Leaf-surface topography and hydrophobicity of wild-type and *Osralf7* mutant plants.

(A) Scanning electron micrographs of adaxial leaf surfaces from wild-type plants and two independent *Osralf7* mutant lines at six weeks of age. Images were obtained at 500× magnification. SC, silica cell; Gla, glandular hair. Scale bars, 50 μm.

(B) Higher-magnification scanning electron micrographs of leaf surfaces from WT and two independent *Osralf7* mutant lines at 2,000× magnification. SC, silica cell; St, stomata; P, papillae. Scale bars, 10 μm.

(C) Leaf-surface hydrophobicity of WT and two independent *Osralf7* mutant lines at five weeks of age, measured by water contact angle analysis. Each circle represents one independent rice plant.

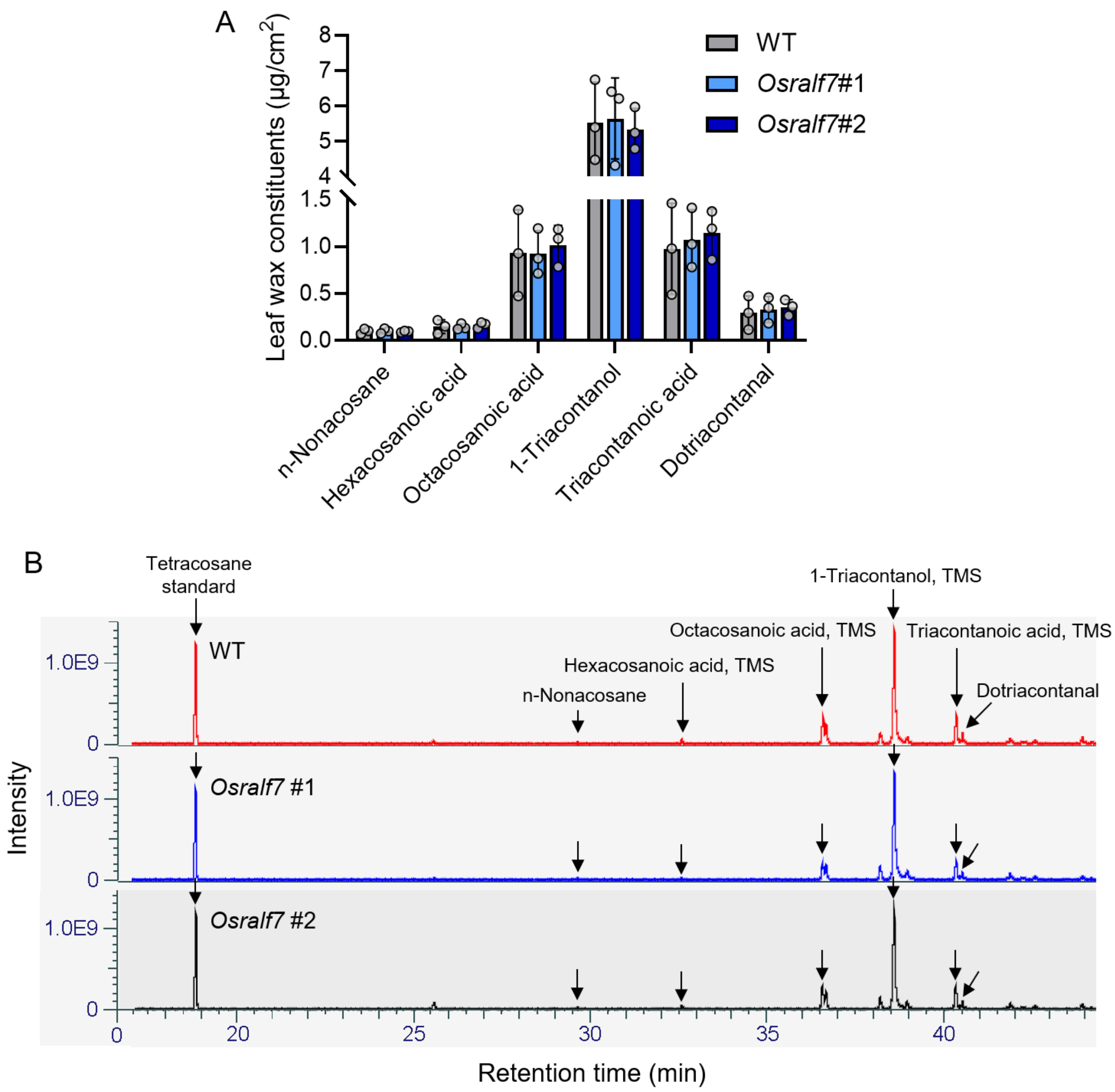


**Fig. S10.** GC–MS analyses of leaf-surface waxes in WT and *Osralf7* knockout plants.

(A) Quantification of abundant leaf-surface wax compounds in WT and *Osralf7* knockout plants at five weeks of age, based on GC-MS Total Ion Chromatogram (TIC) peak-area ratios relative to tetracosane (C24) and normalized to the extracted leaf-surface area. Each circle represents one independent rice plant. Error bars indicate SD.

(B) Representative GC–MS TICs of leaf-surface wax extracts from WT and *Osralf7* knockout plants. Tetracosane was used as an internal standard.

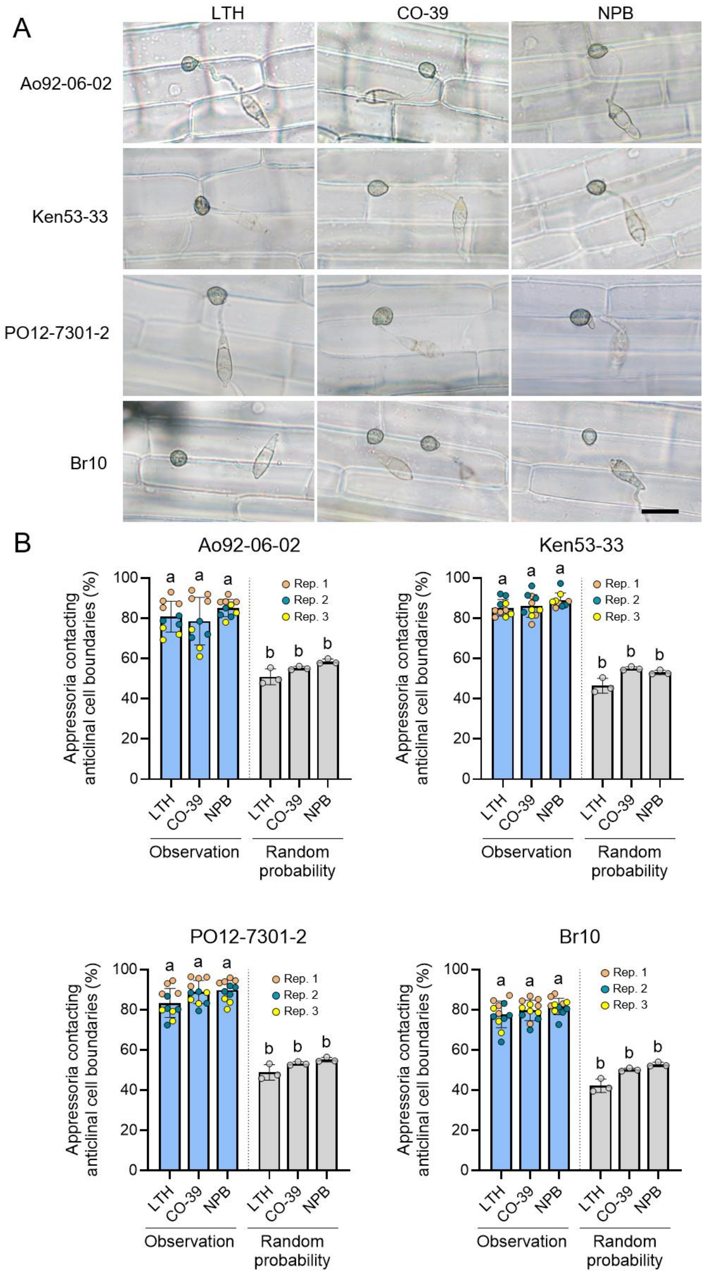


**Fig. S11. Preferential appressorium contact with anticlinal cell boundaries across *M. oryzae* strains and rice cultivars.**

(A) Representative micrographs of four *M. oryzae* strains on leaf sheaths of three rice cultivars at 6 h post inoculation (hpi). Scale bar, 20 μm.

(B) Quantification of the proportion of appressoria contacting anticlinal cell boundaries of rice leaf-sheath cells. Each circle represents one independent rice plant. For each *M. oryzae* strain–rice cultivar combination, three or four individual plants were analyzed, with 150 conidia scored per plant. To estimate the probability expected under random positioning, three individual plants were analyzed for each rice cultivar. For each combination, 90 rice epidermal cells and 60 associated *M. oryzae* appressoria were evaluated, and the expected probability was calculated using the equation shown in Fig. 4*C*. Error bars indicate SD. Statistical significance was assessed by one-way ANOVA followed by Tukey's multiple-comparisons test. Different letters indicate significant differences at $P < 0.05$. Observation data are from three independent experiments.

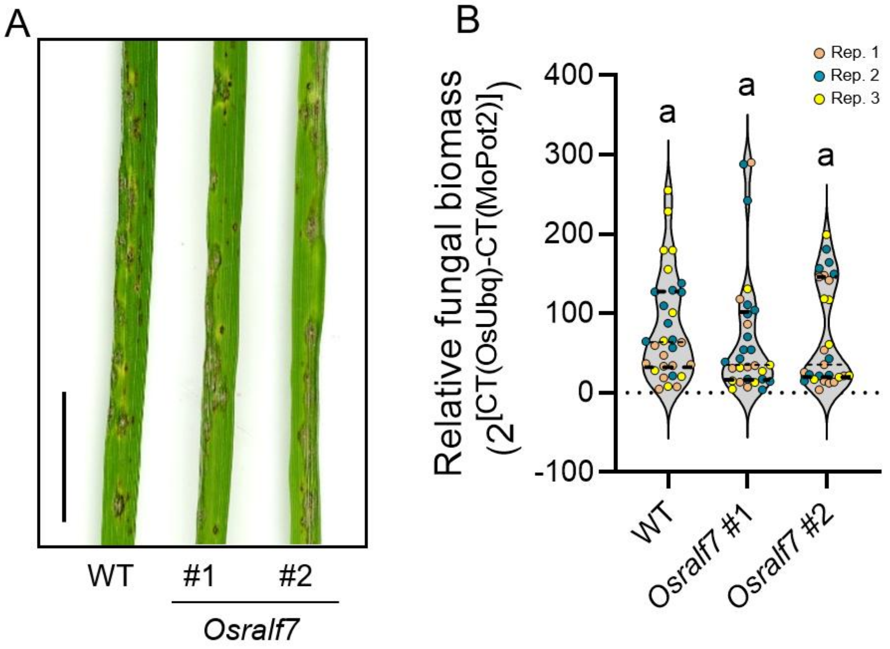


**Fig. S12**. *Osralf7* knockout plants do not show reduced blast lesion formation.

(A) Wild-type plants and two independent *Osralf7* knockout lines were spray-inoculated with the compatible *M. oryzae* isolate Ao92-06-02. Representative images were taken at 7 dpi. Scale bar, 10mm.

(B) Fungal biomass was quantified by qPCR after DNA extraction and is shown as $2^{[CT(OsUbq)-CT(MoPot2)]}$, representing fungal DNA relative to plant DNA. Each dot represents one plant from three independent experiments. Statistical significance was assessed by one-way ANOVA followed by comparison with the wild-type control.

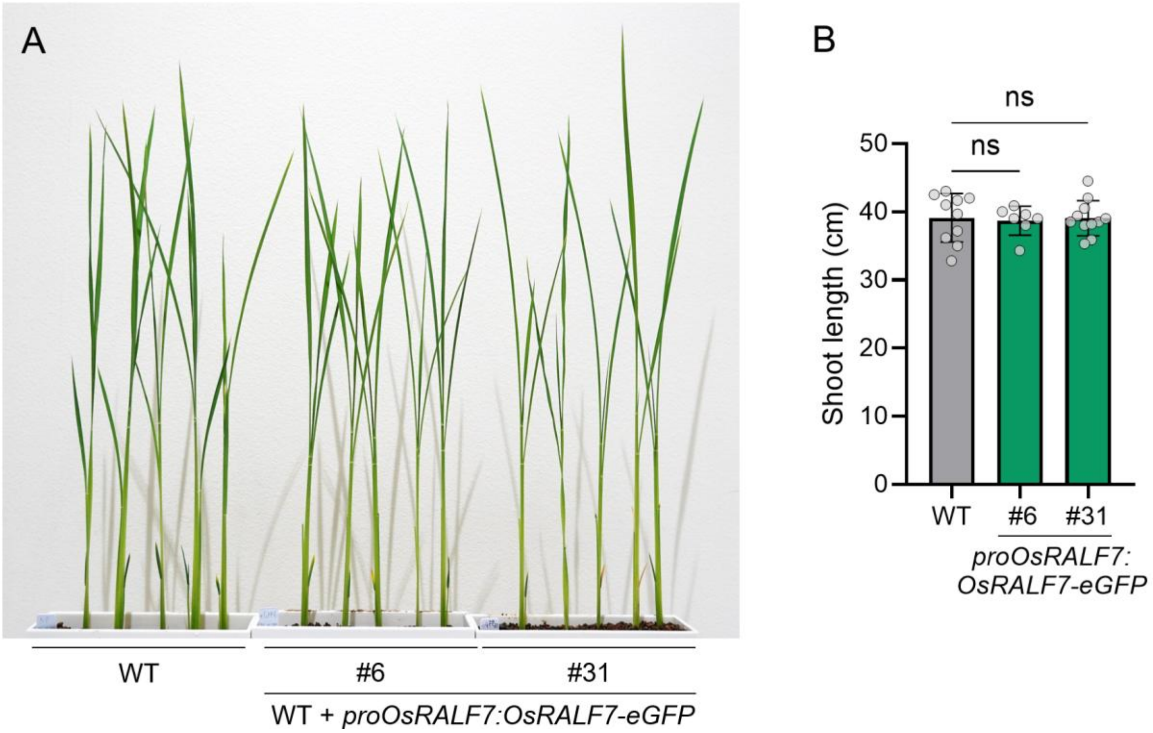


**Fig. S13.** Expression of *proOsRALF7:OsRALF7-eGFP* does not affect shoot growth.

(A) Representative images of shoots and (B) quantification of shoot length in 4-week-old WT (Nipponbare) and transgenic plants expressing *OsRALF7-eGFP* under the native *OsRALF7* promoter. Each dot represents one biological replicate ($n$ = 7–11). Statistical significance was assessed by one-way ANOVA followed by Dunnett's multiple-comparisons test. ns, not significant.

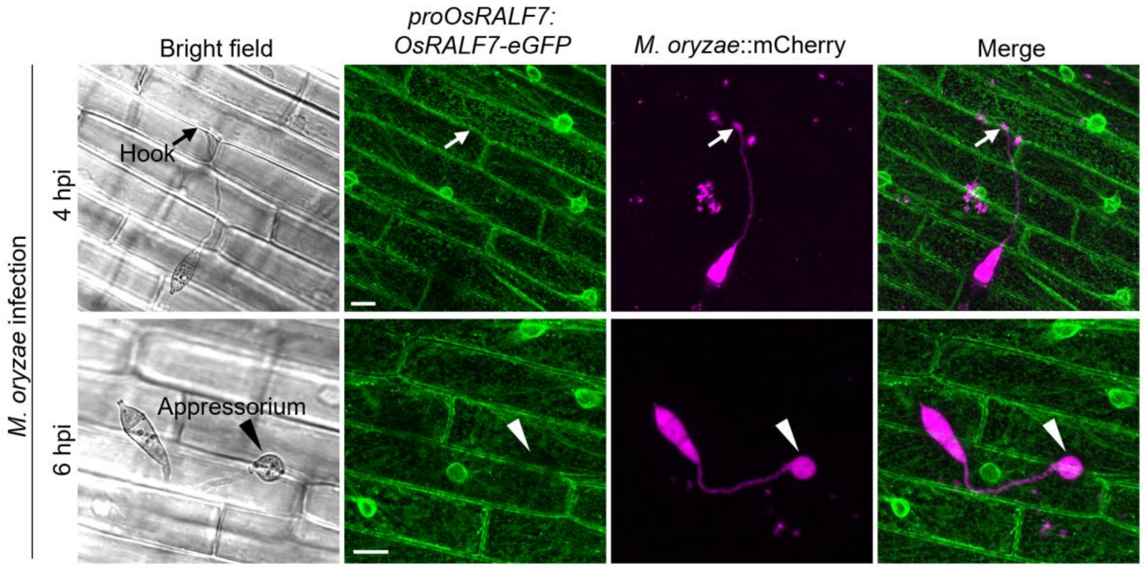


**Fig. S14.** Localization of OsRALF7-eGFP during early *M. oryzae* infection.

Representative fluorescence images of leaf sheath cells expressing *OsRALF7-eGFP* during hook formation (4 hpi) and appressorium formation (6 hpi) after inoculation with *M. oryzae* expressing cytosolic mCherry. Arrows indicate hooks, and arrowheads indicate appressoria. Scale bars, 10μm.

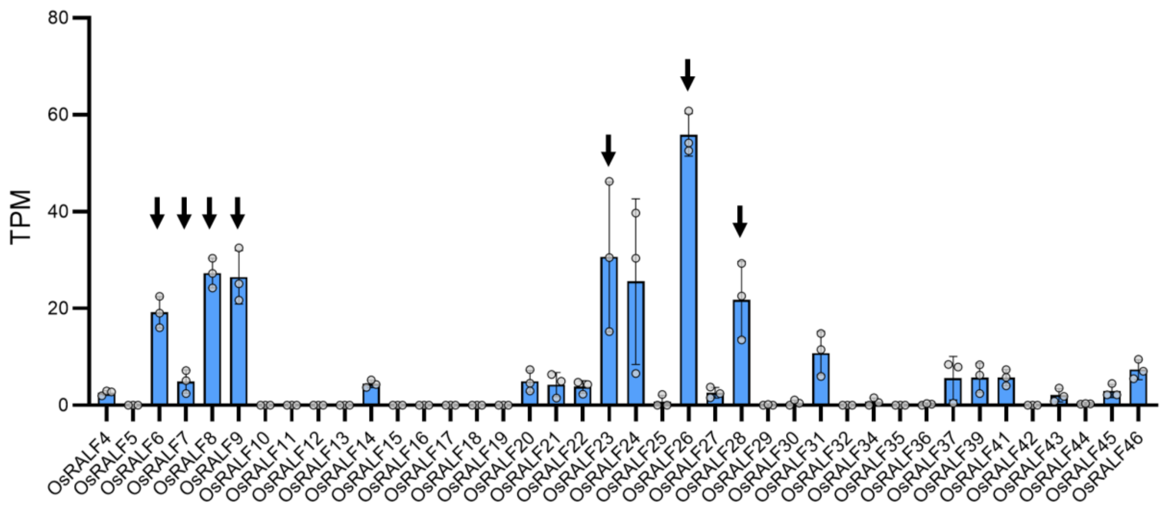


**Fig. S15.** TPM values of all *OsRALFs* in non-infected rice leaves.

TPM values of all putative *OsRALF* genes identified from the MSU database are shown based on previously published RNA-seq data from non-infected rice leaves (40). Arrows indicate the OsRALF peptides examined in this study.

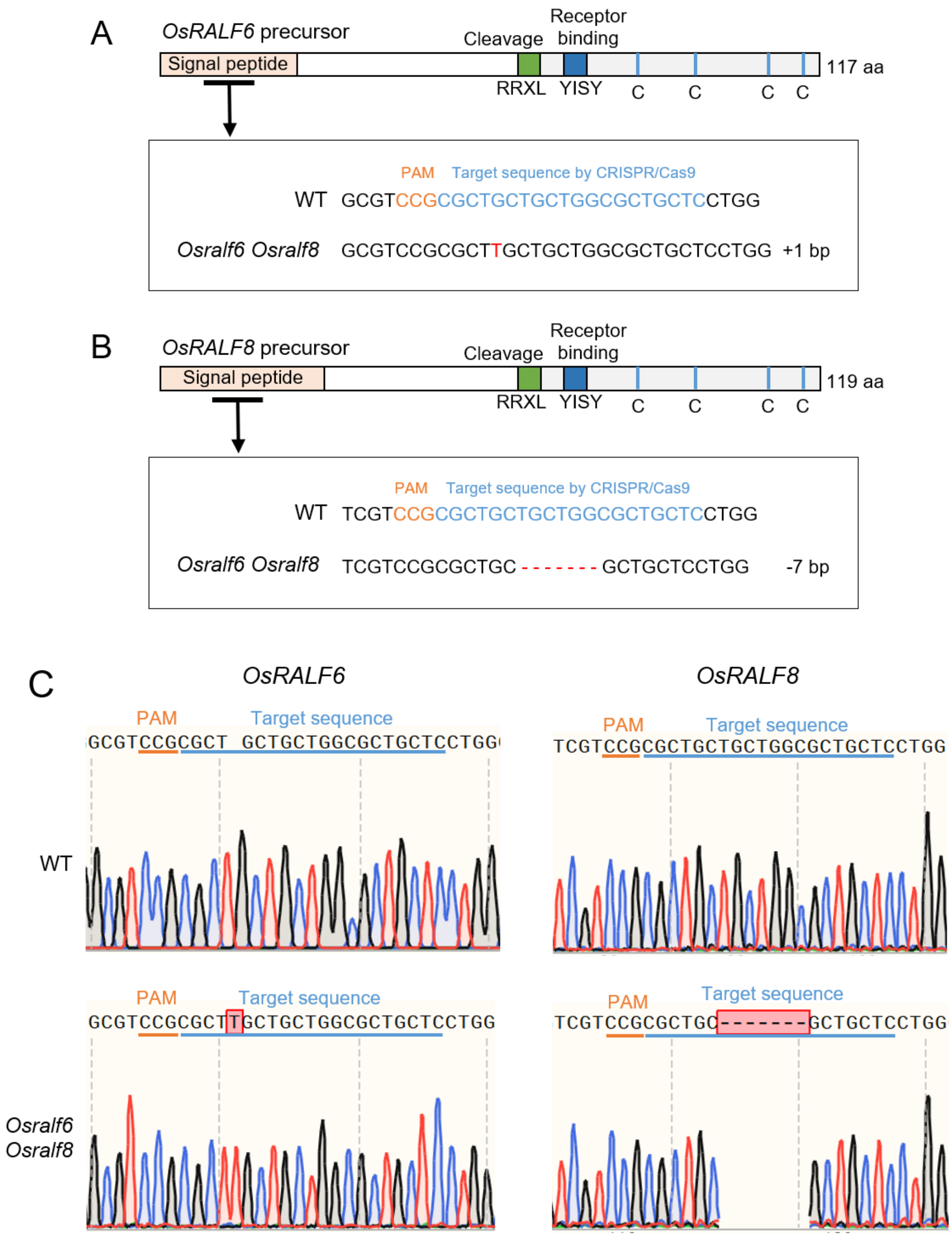


**Fig. S16.** Generation of *Osralf6/Osralf8* double-knockout plants.
(A) Schematic representation of the *OsRALF6* precursor gene and CRISPR–Cas9 target sites. The gRNA target sequence is shown in light blue, and the PAM sequence in orange. The *Osralf6/Osralf8* mutant alleles carry a 1-bp insertion at the target site, resulting in a frameshift and premature translation termination.
(B) Schematic representation of the *OsRALF8* precursor gene and CRISPR–Cas9 target sites. The *Osralf6/Osralf8* mutant alleles carry a 7-bp deletion at the target site, resulting in a frameshift and premature translation termination.
(C) DNA sequences of the *Osralf6/Osralf8* mutant alleles.

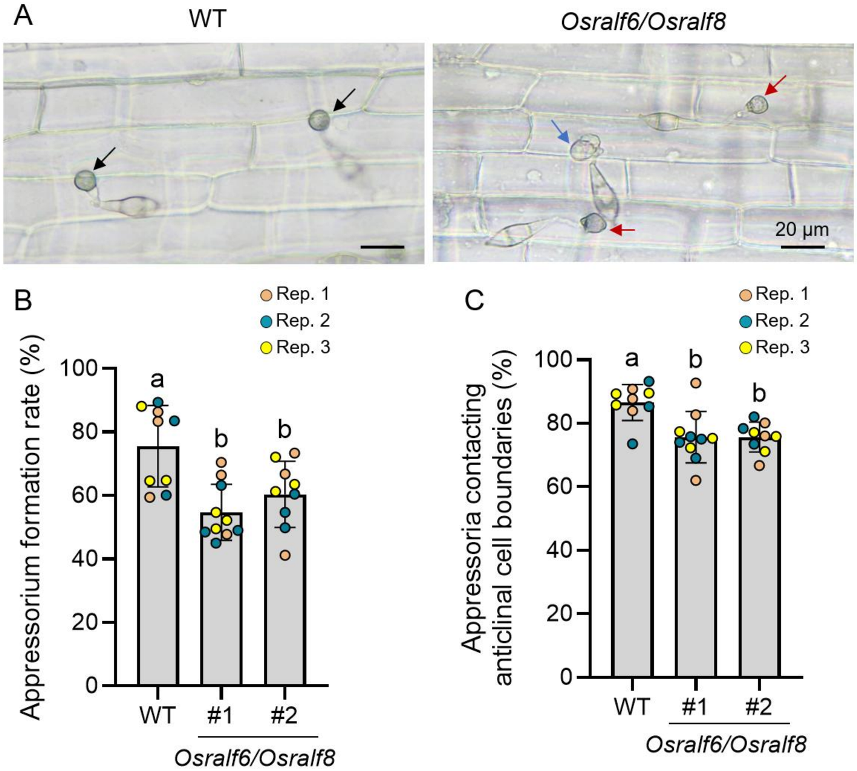


**Fig. S17.** Appressorium formation on *Osralf6/Osralf8* knockout mutants.

(A) Representative micrographs of *M. oryzae* on leaf sheaths of wild-type and *Osralf6/Osralf8* knockout rice plants at 6 h post inoculation (hpi). Black arrows, appressoria contacting anticlinal cell boundaries; red arrow, an appressorium confined to periclinal cell surfaces; blue arrows, immature appressoria. Scale bars, 20 μm.

(B) Appressorium formation frequency on rice leaf sheaths in wild-type and the *Osralf6 Osralf8* knockout. Each circle represents one independent rice plant. For each genotype, three or four individual plants were analyzed, and 150 conidia were scored per plant. Error bars indicate SD. Statistical significance of differences was assessed by one-way ANOVA followed by Tukey's multiple-comparison test. Different letters indicate significant differences at $P < 0.05$. Data are from three independent experiments.

(*C*) Quantification of the proportion of appressoria contacting anticlinal cell boundaries of rice leaf sheath cells. Each circle represents one independent rice plant. For each genotype, three or four individual plants were analyzed, and 150 conidia were scored per plant. Error bars indicate SD. Statistical significance of differences was assessed by one-way ANOVA followed by Tukey's multiple-comparison test. Different letters indicate significant differences at $P < 0.05$. Observation data are from three independent experiments.

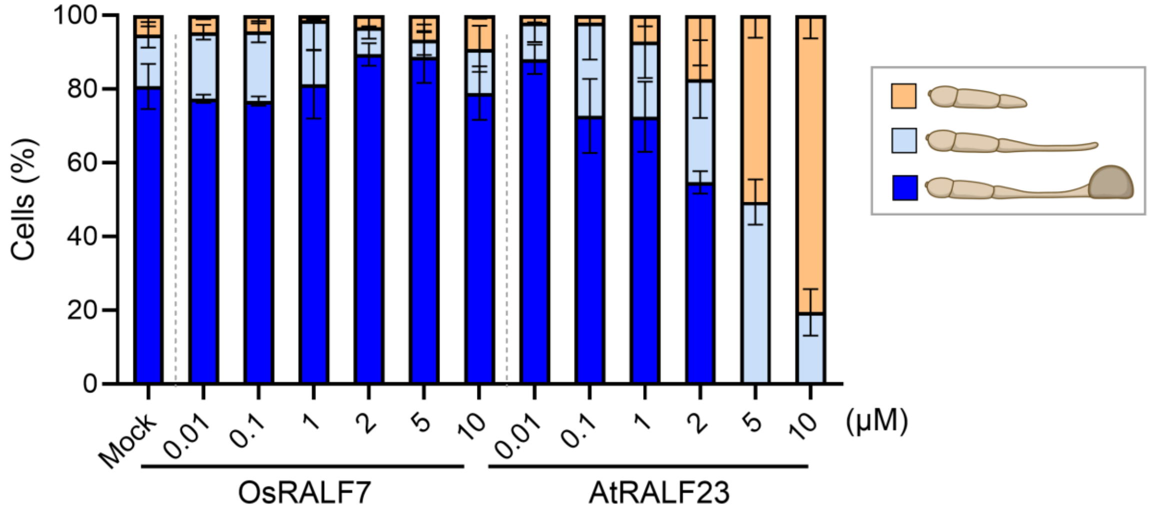


**Fig. S18.** AtRALF23 suppresses conidial germination of *M. oryzae* in water incubation.

*M. oryzae* conidia were treated with water only (Mock), OsRALF7, or AtRALF23 at the indicated concentrations and incubated in water for 8 h. For each treatment and condition, at least 50 conidia were scored per independently cultured plate ($n$ = 3), classified into three morphological categories (no germination, germ tube elongation, and appressorium formation), and the percentage of conidia in each category was calculated. Appressorium formation included both non-melanized and melanized appressoria. Error bars indicate SD. Experiments were performed three times, each time with similar results.

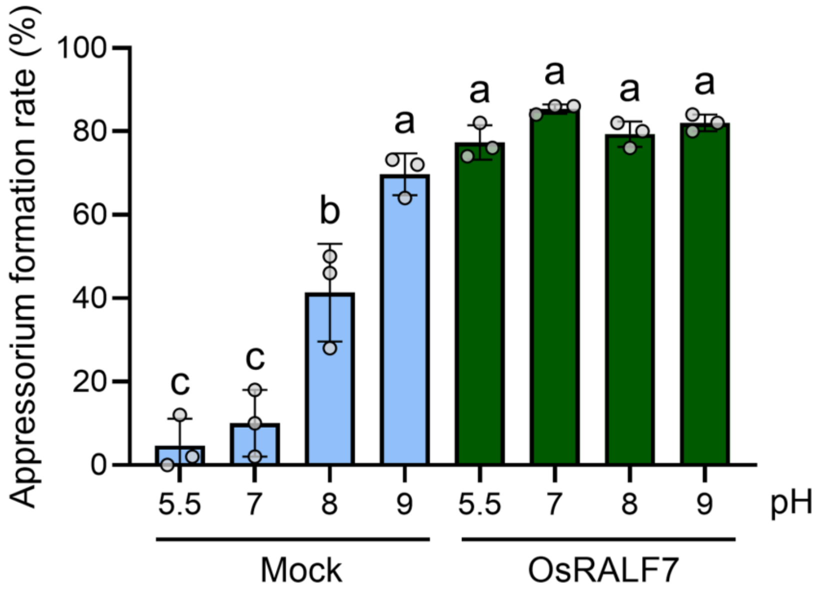


**Fig. S19.** Effect of medium pH on appressorium formation in *M. oryzae*.

*M. oryzae* conidia were treated in 50% CM medium adjusted to the indicated pH values; appressorium formation rates were scored 8 h later. Appressorium formation included non-melanized, melanized, and double appressoria; conidia showing only germ tube elongation were excluded from the appressorium-formed category. Each circle represents one independently cultured plate ($n$ = 3), with >50 conidia scored per plate. Error bars indicate SD. Statistical significance of differences was assessed by one-way ANOVA followed by Tukey's multiple comparison test. Different letters indicate significant differences at $P < 0.05$.

### Tables

**Table S1.** Read quality assessment and read mapping summary.

| sample | Total Raw Reads | Total Clean Reads | Clean Q30 Rate | Mapped Reads | Properly Paired Reads | Mapped Reads (Bowtie) | Properly Paired Reads (Bowtie) |
|---|---|---|---|---|---|---|---|
| WT_Mock_rep1 | 38,872,304 | 38,491,908 | 0.9576 | 38,327,271 | 38,303,898 | 57,955 | 9,678 |
| WT_Mock_rep2 | 33,642,096 | 33,289,918 | 0.9565 | 33,153,412 | 33,132,744 | 55,535 | 8,460 |
| WT_Mock_rep3 | 38,323,494 | 37,911,400 | 0.9553 | 37,772,656 | 37,749,170 | 54,373 | 10,464 |
| WT_Chitin_rep1 | 44,784,264 | 44,334,408 | 0.9577 | 44,178,009 | 44,153,006 | 63,955 | 11,144 |
| WT_Chitin_rep2 | 44,269,466 | 43,844,322 | 0.9577 | 43,699,141 | 43,674,014 | 57,441 | 10,254 |
| WT_Chitin_rep3 | 50,163,108 | 49,667,198 | 0.9576 | 49,497,208 | 49,467,824 | 68,650 | 12,114 |
| WT_RALF7_rep1 | 37,830,792 | 37,444,686 | 0.9569 | 37,304,162 | 37,281,378 | 58,362 | 10,212 |
| WT_RALF7_rep2 | 34,883,206 | 34,565,578 | 0.9591 | 34,427,735 | 34,408,516 | 53,641 | 8,136 |
| WT_RALF7_rep3 | 33,069,470 | 32,763,620 | 0.9577 | 32,642,788 | 32,623,516 | 46,905 | 8,402 |

**Table S2.** Fungal strains used in this study.

| Species | Strain or isolate designation | Figs presented in this study | Note |
|---|---|---|---|
| *Magnaporthe oryzae (syn. Pyricularia oryzae)* | Ao02-06-02 (MAFF101530) | Fig. 4<br>Figs. S12 and S17 | Obtained from NARO, Japanese isolate |
| | Ao02-06-02:TEF-mCherry | Fig. S14 | This study |
| | Hoku1: Histone-GFP | Figs. 2 and 6<br>Figs. S5, S6, S18, S19 | (41) |
| | P2-wild type | Fig. 3<br>Figs. S4 and S7 | Japanese isolate |
| | P2: hph-ectopic | Fig. 3E, Fig. S7 | This study |
| | P2: *Δmsb2* | Fig. 3E, Fig. S7 | This study |
| | P2: Δ*sho1* | Fig. 3E, Fig. S7 | This study |
| | P2: Δ*pth11* | Fig. 3E, Fig. S7 | This study |
| | P2: Δ*pmk1* | Fig. 3E<br>Figs. S4 and S7 | This study |
| | P2: Δ*cpka* | Fig. 3E<br>Figs. S4 and S7 | This study |
| | Br10 | Fig. S11 | Brazilian isolate |
| | Ken35-33 | Fig. S11 | Japanese isolate |
| | PO-12-7301-2 | Fig. S11 | Indonesian isolate |
| *Colletotrichum higginsianum* | MAFF 305635 | Fig. 7 | NARO,<br>Japanese isolate |

**Table S3.** List of highly expressed OsRALFs in rice leaves.

| gene_name | MSU_gene_ID | Full sequence[*1] (SP, RRXL, YIXY, **conserved C-term**) | SP[*2] | RRXL[*3] | YIXY[*4] | C[*5] | Publications |
|---|---|---|---|---|---|---|---|
| OsRALF6 | LOC_Os01g25540 | MAKSLLAASALLLALLLAVAGGAAAAGEVPLAWELGVGGGGGGGEEDSFGFSSEDAAADGAAVVRRVL**QGQGYISYGALRRDTTPCSVRGASYYNCRPGGQANPYSRGCSAITRCRG*** | yes | yes | yes | 4 | |
| OsRALF7 | LOC_Os01g25560 | MARPAIVAAPALLLLALLIALATGGADAAPGEVPLSWELGVVGADDAFGFPGEEAADSATAVVRRVL**QQGSYISYGALRRDTTPCSVRGASYYNCQPGAEANPYSRGCSAITQCRG*** | yes | yes | yes | 4 | (40) |
| OsRALF8 | LOC_Os02g44940 | MARAATVAAASSALLLALLLALAWGAAAGAGEVPLGWELGVGVGGGSGDDDGFGFSGAAADGAAVVRRVL**QGRGYISYGALRRDTTPCSVRGASYYNCRPGGQANPYSRGCSAITRCRG*** | yes | yes | yes | 4 | (40) |
| OsRALF9 | LOC_Os01g15320 | MAAAAVALLAFLLAAASAASSASAAAATLVEGGVVGRAAVVMRRGGRTCRGTVGECMEFLGVDGEGEDELAAAATGKRRVL**QGGSGYIGYDALRRDSVPCSQRGASYYNCQPGAEANPYSRGCSAITQCRG*** | yes | yes | yes | 4 | |
| OsRALF23 | LOC_Os11g26190 | MKLMLLFAATVVAAALVLVAAVDVGSSGALLPMHVLRGRVADVEDVVASVVEEEEAAYPRRRIL**YDSRYASYNGLTESKAACYGSCPGRGQPYSGRSCLNIYQCKG*** | yes | yes | YASY | 4 | |
| OsRALF26 | LOC_Os10g41980 | MMRKQSRSPCSSSSSAAVVSLCKVLLLVLALICTLETVSVEGGRVAAAALVGGGVPLNPRYPAI**PEEPYIGRRGDPYTGRGGEPYTRPGRGCTVAYGCYGGPPAAKP*** | yes | no | YIGR | 2 | (42, 43) |
| OsRALF28 | LOC_Os06g29730 | MASLRATTLLVLMQIIMVFYTVILSCSFSDA**RTFPGGEGGLDPNHPVCVGGACPTPGLPYTNPRDPCIYRNRCNPPGRMGDP*** | yes | no | PGGE | 4 | |

[*1] The full-length protein sequence was retrieved from the MSU Rice Genome Annotation Project database (https://rice.uga.edu/).
[*2] Signal peptide predicted using SignalP 6.0 (https://services.healthtech.dtu.dk/services/SignalP-6.0/).
[*3] Predicted cleavage site.
[*4] Conserved CrRLK1L/LLG receptor-binding motif.
[*5] Number of cysteine residues in the conserved C-terminal sequence.

**Table S4.** Evolutionary distances from OsRALF7 to AtRALFs.

| Gene name[a] | Gene ID | Distance[b] | C-terminal sequence[a] |
|---|---|---|---|
| AtRALF23 | At3g16570 | 0.301811919 | ATRRYISYGALRRNTIPCSRRGASYYNCRRGAQANPYSRGCSAITRCRRS |
| AtRALF33 | At4g15800 | 0.311901214 | ATTKYISYGALRRNTVPCSRRGASYYNCRRGAQANPYSRGCSAITRCRR |
| AtRALF22 | At3g05490 | 0.323432267 | AQKKYISYGAMRRNSVPCSRRGASYYNCQRGAQANPYSRGCSTITRCRR |
| AtRALF1 | AT1G02900 | 0.4716909 | ATTKYISYQSLKRNSVPCSRRGASYYNCQNGAQANPYSRGCSKIARCRS |
| AtRALF19 | AT2G33775 | 0.58889304 | AARRSYISYGALRKNNVPCSRRGRSYYDCKKRKRANPYRRGCSVITHCYRQTS |
| AtRALF34 | At5g67070 | 0.682918408 | YWRRTKYYISYGALSANRVPCPPRSGRSYYTHNCFRARGPVHPYSRGCSSITRCRR |
| AtRALF24 | At3g23805 | 0.696784434 | MMRKQYISYETLRRDMVPCQKPGASYYACRSGQANAYNRGCSVITRCARDTNDIKT |
| AtRALF31 | At4g13950 | 0.700801024 | MAQKRYIGYETLRRDMVPCQKPGASYYDCRSGQANSYSRGCDTITRCARDTNDINT |
| AtRALF4 | AT1G28270 | 0.795436543 | ARGRRYIGYDALKKNNVPCSRRGRSYYDCKKRRRNNPYRRGCSAITHCYRYAR |
| AtRALF11 | AT2G19030 | 0.809037527 | EPVESRNYIEYGAINKCAGPNPPPGCNPPGAEQKNPTPVNEYSRGCSKIHRCRRD |
| AtRALF32 | At4g14010 | 0.854959146 | EQAHKLSYGALRRNQPACDGGKRGESYSTQCLPPPSNPYSRGCSKHYRCGRDS |
| AtRALF12 | AT2G19040 | 0.904419275 | EPVESRNYIEYGAINKCAGPNPPPGCNPPGTEQKNPTPVNEYSRGCSKIHRCRRD |
| AtRALF28 | At4g11510 | 1.042644651 | NEIGYPGMGRGDRQPGCDHGNCPPDQPANPYHRGCEKSKRCRGPDPPALPRKMI |
| AtRALF9 | AT1G61566 | 1.084271902 | TRYITYPAIDRGDHAVHCDKAHPNTCKKKEANPYQRGCEKINRCRGG |
| AtRALF14 | AT2G20660 | 1.107208561 | QASRYISYEALKKNLPDNRRGEPDQRDNPYRRSCDVHSHCYRFTN |
| AtRALF13 | AT2G19045 | 1.174677985 | EPVESRNYIEYGAINKCAGPNPPPGCNPPGAEQKNPNPVNEYSRGCSKIHRCRRD |
| AtRALF15 | AT2G22055 | 1.213377942 | TRYISYRGMNHGDHAIHCDKAHPNTCKKQVANPYRRGCGTIERCRRDTGRK |
| AtRALF5 | AT1G35467 | 1.228405574 | AKRYIEYPPWQKHPCNPRFPTPDCYKRTPANPYRRGCTCISRCRRDCGGLSTWKKLLDTILKIPV |
| AtRALF25 | At3g25165 | 1.235569614 | NDNKRKYLLLDPCLRPNAPPGCHRQPYKPRTPVNVYSRGCTTINRCRRVQNP |
| AtRALF21 | At3g04735 | 1.25949071 | KVIGYPGLKPDLPCDHHRYPSACAPSEQPVNPYRRGCSKIHRCRRDSPPAPISRKMLIRGQLIYNN AYNAYIQYP |
| AtRALF18 | AT2G33130 | 1.302535035 | AKRFIDYEALKKNLPAKPDGKPDKPDNKYRRGCSAATGCYRFTN |
| AtRALF20 | At2g34825 | 1.313688268 | AKTIGNPAMREDEPKGCPPGSPASCKMQPANPYKPGCEASQRCRGT |
| AtRALF3 | AT1G23147 | 1.320516369 | RKEIGYPKQRFGEDRTNPYEEITPPLIGGCDPKNPQTCLPKQPANPYRRGCLKITRCQRDV |
| AtRALF2 | AT1G23145 | 1.330987572 | AQKVIGYPAIGRDGARGCSPKDPSCPQQPEKPYKRGCEKITRCERDRRKQAHLRNPRKVLDVVAVM AKAKQLY |
| AtRALF17 | AT2G32885 | 1.339815268 | NSIGAPAMREDLPKGCAPGSSAGCKMQPANPYKPGCEASQRCRGG |
| AtRALF6 | AT1G60625 | 1.346785045 | QTYINYNGMKGDIIPGCSSKNPKECVKIPAYSYNRGCEISTRCQRQQHSSSS |
| AtRALF26 | At3g25170 | 1.378488464 | RKGRKYLNPGVLDRCRGPNPPAGCHPHNSHHKPRVPVHNYSRGCSRITRCRRDA |
| AtRALF35 | AT1G60913 | 1.385125271 | RYIKYRAIAKDRVPDCTQDPKNCVRVPVNQYHLPPGCQNTTHCYREKYHI |
| AtRALF8 | AT1G61563 | 1.421216342 | SVRYITYPAIDRGDHAVHCDKAHPNTCKKKQANPYRRGCGVLEGCHRETGPKPT |
| AtRALF10 | AT2G19020 | 1.421350923 | VPVESRRKHLDYGVITKCAGPNPPPGCYPPGAQQKNPTPANEYRRGCSKITRCKRD |
| AtRALF7 | AT1G60815 | 1.424648555 | IKQINYKDLIKDTIPGCTSKNPKECVKVPANTYHRGCEISTRCHREQHSSSG |
| AtRALF16 | AT2G32835 | 1.457532137 | RTLGYGSIKGDRIPACGYKNPNSCVKQPVNHYHRGCEKITRCARDAARYTESFNVDDDESPIINLH |
| AtRALF29 | At4g11653 | 1.49958705 | AKRYIEYPIRLDLGKGCDPRFPTAACYKRTPANPYRRPCTTANRCRRSTSSTRVPSLKTFVEIPPM |
| AtRALF27 | At3g29780 | 1.577616917 | QQRKYLSYKTLQKQPTCDGRIAGNCIGTVNPKGATCTYYQRCKRAA |
| AtRALF36 | AT2G32785 | 1.587823068 | DTINREQVISYESMRVNHAWGCSQKYPQFCQKTRANPYTKPPPKNSEAS |
| AtRALF37 | AT2G32788 | 1.6472905 | EDVISYEVLLQDHAWGCSPKFPRLSCLKQKANP |
| AtRALF30 | At4g13075 | 2.154762831 | AGGGKFLNPGVLDPCLRPNPPPECQAPGSAGKPRERVNEYKVGCSKLTRCDRVG |
| AT2G32890 | AT2G32890 | 3.390287173 | GAAADVSRGGCGGGDGSLGDDNERCVEAVKEDDDDDVDDVYKVINKMRIYA* |

[a]Gene names and C-terminal sequences are as reported in (38).
[b]Following MUSCLE-based multiple sequence alignment of the amino acid sequences (C-terminal regions of OsRALF7 and AtRALFs), a maximum-likelihood distance matrix was computed in phangorn under the WAG substitution model (dist.ml).

**Table S5.** Synthesized peptides used in this study.

| Peptide name | Sequence |
|---|---|
| OsRALF7 | QQGSYISYGALRRDTTPCSVRGASYYNCQPGAEANPYSRGCSAITQCRG |
| OsRALF7 YISY→AAAA | QQGSAAAAGALRRDTTPCSVRGASYYNCQPGAEANPYSRGCSAITQCRG |
| OsRALF7 $^{\Delta YISY}$ | QGALRRDTTPCSVRGASYYNCQPGAEANPYSRGCSAITQCRG |
| YISY | QQGSYISY |
| OsRALF7 $^{4C\rightarrow 4A}$ | QQGSYISYGALRRDTTPASVRGASYYNAQPGAEANPYSRGASAITQARG |
| AtRALF23 | ATRRYISYGALRRNTIPCSRRGASYYNCRRGAQANPYSRGCSAITRCRRS |
| OsRALF6 | QGQGYISYGALRRDTTPCSVRGASYYNCRPGGQANPYSRGCSAITRCRG |
| OsRALF8 | QGRGYISYGALRRDTTPCSVRGASYYNCRPGGQANPYSRGCSAITRCRG |
| OsRALF9 | QGGSGYIGYDALRRDSVPCSQRGASYYNCQPGAEANPYSRGCSAITQCRG |
| OsRALF23 | YDSRYASYNGLTESKAACYGSCPGRGQPYSGRSCLNIYQCKG |
| OsRALF26 | GRVAAAALVGGGVPLNPRYPAIPEEPYIGRRGDPYTGRGGEPYTRPGRGCTVAYGCY GGPPAAKP |
| OsRALF28 | RTFPGGEGGLDPNHPVCVGGACPTPGLPYTNPRDPCIYRNRCNPPGRMGDP |
| flg22 | QRLSTGSRINSAKDDAAGLQIA |

**Table S6.** Primers used in this study.

| Primer ID | Primer name | Sequence | Note | Application |
|---|---|---|---|---|
| oFF51 | OsU6-2_F5 | tccccctttcgccagctggcgtaatagcgaagagg | Amplification of a 2394-bp pUC19 backbone | pU6gRNA_RALF7_gRNA2 and pU6gRNA_RALF6/8_gRNA construction |
| oFF52 | OsU6-2_R5 | aacctgtcgtgccagctgcattaatgaatcggcca | | |
| oFF38 | OsU6-2_F4 | gattcattaatgcagctggcacgacaggtttccc | Amplification of a 584-bp partial pUC19 backbone fragment containing the OsU6−2 promoter fused to the *OsRALF7* or *OsRALF6/OsRALF8* gRNA target sequence | |
| oFF108 | RALF7_Crisp_R1 | ccacgatggcgggtctcgcccAACACAAGCGACAGCGCGCGGGTT | | |
| oFF109 | RALF6-8_Crisp_F1 | gagcagcgccagcagcagcgGTTTTAGAGCTAGAAATAGCAAGTT | | |
| oFF107 | RALF7_Crisp_F1 | ggcgagacccgccatcgtggGTTTTAGAGCTAGAAATAGCAAGTT | Amplification of a 330-bp fragment containing the OsRALF7 gRNA target sequence fused to the gRNA scaffold, followed by a poly(T) terminator<br>Amplification of a 330-bp fragment containing the OsRALF7 or *OsRALF6/OsRALF8* gRNA target sequence fused to the gRNA scaffold, followed by a poly(T) terminator | |
| oFF39 | OsU6-2_R4 | tcgctattacgccagctggcgaaagggggatgtg | | |
| oFF110 | RALF6-8_Crisp_R1 | cgctgctgctggcgctgctccAACACAAGCGACAGCGCGCGGGTT | | |
| oFF36 | OsU6-2_F3 | ACAGGGTAATttaattaaGGTACCGAGCTCGGATCCA | Amplification of a 513-bp fragment containing the OsU6-2 promoter, the OsRALF7 gRNA target sequence, and a poly(T) terminator. | pZH_gYSA_PubiMMCas9-RALF7_gRNA2 construction |
| oFF37 | OsU6-2_R3 | CAGGGTAATggcgcgccAAGCTTATCGATACCGTCGA | | |
| oFF425 | RALF7_R8 | cgaactttgaatcacaaatcccaatttc | | Genopyting for *Osralf7* knockout plants |
| oFF426 | RALF7_F9 | aaagcaagcaaccaattcctcctc | | |
| oFF2 | RALF7_R1 | TCATCCCCTGCACTGGGTGAT | | |
| oFF121 | RALF6_F1 | agcaagcaaacaccaaaggcag | | Genopyting for *OsRALF6* gene in *Osralf6/Osralf8* double knockout plants |
| oFF127 | RALF6_R3 | actcgattcgattcgtgggcaccgac | | |
| oFF123 | RALF8_F1 | attacgctatccgccacgag | | Genopyting for *OsRALF8* gene in *Osralf6/Osralf8* double knockout plants |
| oFF124 | RALF8_R1 | tagacggagaggagcaacgaacgaa | | |
| oFF197 | GFP_R1 | gaagtcgatgcccttcagctcgatgc | | Genotyping for ProRALF7-RALF7-eGFP plants |
| oFF323 | eGFP_F1 | gggagccgacgcggcggtgagcaagggcgag | | |
| 1105 | Ubq qPCR F | AACCAGCTGAGGCCCAAGA | (45) | Fungal biomass analysis |
| 1106 | Ubq qPCR R | ACGATTGATTTAACCAGTCCATGA | | |
| 927 | MgPot2-RT-F q-PCR | ACGACCCGTCTTTACTTATTTGG | | |
| 928 | MgPot2-RT-R q-PCR | AGTAGCGTTGGTTTTGTTGGAT | | |
| YH375 | CPKA target-1 | ttcgtgtcttgtccggtacacgct | Oligonucleotides for construction of pcc-pka | Construction of CRISPR/Cas9 expression vector |
| YH376 | CPKA target-2 | aaacagcgtgtaccggacaagaca | | |
| TS44 | PMK1 target-1 | AAACcggaaaggactgtaagttac | Oligonucleotides for construction of pcc-pmk1 | |

| | | | | |
|---|---|---|---|---|
| TS45 | PMK1 target-2 | TTCGgtaacttacagtcctttccg | | |
| TS29 | MSB2 target-1 | AAACctgcggttgtctctaccaca | Oligonucleotides for construction of pcc-msb2 | |
| TS30 | MSB2 target-2 | TTCGtgtggtagagacaaccgcag | | |
| TS35 | SHO1 target-1 | AAACttttgcattggcgacgatat | Oligonucleotides for construction of pcc-sho1 | |
| TS36 | SHO1 target-2 | TTCGatatcgtcgccaatgcaaaa | | |
| TS23 | PTH11-target-1 | AAACcccttcatgtcgagcaatag | Oligonucleotides for construction of pcc-pth11 | |
| TS24 | PTH11-target-2 | TTCGctattgctcgacatgaaggg | | |
| YH333 | CPKA-FL1 | tagaactagtgagggaaagcatgtttgcttagctg | Amplification of 876 bp 5′ flanking region of *CPKA* with SpeI and HindIII sites | pPka-donor construction |
| YH334 | CPKA-FL2 | cgataagcttgattcctgcaagcaaagctagaagg | | |
| YH335 | CPKA-FR1 | ccttgcatgcacgaaaacacagtcatgctgtgacg | Amplification of 1,043 bp 3′ flanking region of *CPKA* with SpeI and XhoI sites | |
| YH336 | CPKA-FR2 | cccccccctcgagtggctggcataagcattgtgtcc | | |
| TS31 | MSB2-FL1 | gatAAGCTTTtcctttccttcgctaccctatcctc | Amplification of 1,111 bp 5′ flanking region of *MSB2* with HindIII site and overlap sequence for fusion PCR | pMsb2-donor construction |
| TS32 | MSB2-FL2 | aggctgtcaattcctgggccgaagacgacgaggcc | | |
| TS33 | MSB2-FR1 | ttcggcccaggaattgacagcctctggaactgatcgcaatgcca | Amplification of 1,030 bp 3′ flanking region of *MSB2* with SpeI site and overlap sequence for fusion PCR | |
| TS34 | MSB-FR2 | agaACTAGTgttgacaaaaagacaagggaaaatcg | | |
| TS37 | SHO1-FL1 | tccACTAGTgcctctcccgacattaaccctcctcc | Amplification of 1,011 bp 5′ flanking region of *SHO1* with SpeI site and overlap sequence for fusion PCR | pSho1-donor construction |
| TS38 | SHO1-FL2 | gctcatgcctttcctaccacctccgtactgtcctc | | |
| TS39 | SHO1-FR1 | gaggtggtaggaaaggcatgagcgcggactgaacgaagctcagg | Amplification of 1,300 bp 3′ flanking region of of *sho1* with SacI site and overlap sequence for fusion PCR | |
| TS40 | SHO1-FR2 | ctgGAGCTCcagtgccggagacagtagcccttgca | | |
| TS25 | PTH11-FL1 | ttgGGTACCcctagaagatgttcaacgtcgcatc | Amplification of 1,217 bp 5′ flanking region of *PTH11* with KpnI site and overlap sequence for fusion PCR | pPth11-donor construction |
| TS26 | PTH11-FL2 | gtttgttgatcgactatgactgaagccgctccgc | | |
| TS27 | PTH11-FR1 | tcagtcatagtcgatcaacaaacgactgcccggtggtctcatct | Amplification of 1,043 bp 3′ flanking region of of *PTH11* with SphI site and overlap sequence for fusion PCR | |
| TS28 | PTH11-FR2 | cttGCATGCggcgtccacgggcgccggcctggcca | | |
| TS46 | PMK1-FL1 | gatAAGCTTtgaaccagacatgtcagatacgatag | Amplification of 1,042 bp 5′ flanking region of *PMK1* with HindIII site and overlap sequence for fusion PCR | pPmk1-donor construction |
| TS47 | PMK1-FL2 | ggtggattggcgcgagacatggcggcgctgaaagg | | |
| TS48 | PMK1-FR1 | gccatgtctcgcgccaatccaccagttcatctaccaggaaatta | Amplification of 1,166 bp 3′ flanking region of *PMK1* with SpeI site and overlap sequence for fusion PCR | |
| TS49 | PMK1-FR2 | agaACTAGTctgtttgtgcgccggtttttttttacg | | |
| YH361 | CPKA-check1 | tgacaccaacaagcatccaacttgtcac | Binding outside the flanking regions of *CPKA* | Genotyping for Δ*cpka* using CPKA-check1/2, CPKA-check1/Hph-check2, and CPKA-check2/Hph-check1 primer sets |
| YH362 | CPKA-check2 | Cagttgccacttcgtaacttcgaacc | | |

| | | | | |
|---|---|---|---|---|
| TS56 | MSB2-check | atcacgtcggcgattaagggcttcgagttcaatgg | Binding outside the 3′ flanking region of *MSB2* | Genotyping for Δ*msb2* using MSB2-check/MSB2-FL1 and MSB2-check1/ Hph-check2 primer sets |
| TS41 | SHO1-check | cgtcaccaccaccaccaccacttgtaactccattg | Binding outside the 3′ flanking region of *SHO1* | Genotyping for Δ*sho1* using SHO1-check/MSB2-FL1 and SHO1-check/ Hph-check2 primer sets |
| TS57 | PTH11-check | aaggatggctgctcagccgcagtcgggcggcaatg | Binding outside the 3′ flanking region of *PTH11* | Genotyping for Δ*pth11* using PTH11-check/PTH11-FL1 and PTH11-check/ Hph-check2 primer sets |
| TS59 | PMK1-check | gagacgagcgccgccggccctgatccccatatacg | Binding outside the 3′ flanking region of *PMK1* | Genotyping for Δ*pmk1* using PMK1-check/MSB2-FL1 and PMK1-check/ Hph-check2 primer sets |
| TA693 | Hph-check1 | CCCAGCACTCGTCCGAGGGCAAAGGAATAG | Binding to *Hph* | Genotyping for knockout mutants |
| TA694 | Hph-check2 | TCGGACACGCTGTCGAACTTTTCGATCAGA | Binding to *Hph* | |

**Table S7.** Plasmids used in this study.

| Plasmid name | Note | |
|---|---|---|
| pU6gRNA_RALF7_gRNA2 | To generate pZH_gYSA_PubiMMCas9-RALF7_gRNA2 | This study |
| pU6gRNA_RALF6/8_gRNA | To generate pZH_gYSA_PubiMMCas9-RALF6/8_gRNA | This study |
| pZH_gYSA_PubiMMCas9-RALF7_gRNA2 | Used for transformation to generate *Osralf7*-ko | This study |
| pZH_gYSA_PubiMMCas9-RALF6/8_gRNA | Used for transformation to generate *Osralf6/Osralf8*-double ko | This study |
| pPBV[Exp]-Hygro-{RALF7}{RALF7}-3xGGGGS-EGFP | Used for transformation to generate NP: pOsRALF7-OsRALF7-eGFP | vector builder ID: VB231211-1141myr |
| pBI-TEF-mCherry-SUR | To generate Ao02-06-02:TEF-mCherry | (3) |
| pMK-dGFP | To generate donor vectors | (5) |
| pCRISPR/Cas9-FoU6-FoNLSx2 | To generate pCRISPR/Cas9-PU6-FoNLSx2 | (7) |
| pCRISPR/Cas9-PU6-FoNLSx2 | To generate CRISPR/Cas9 expression vector for *M. oryzae* | This study |
| pPka-donor | To generate Δ*cpka* | This study |
| pcc-pka | | |
| pMsb2-donor | To generate Δ*msb2* | This study |
| pcc-msb2 | | |
| pPmk1-donor | To generate Δ*pmk1* | This study |
| pcc-pmk1 | | |
| pSho1-donor | To generate Δ*sho1* | This study |
| pcc-sho1 | | |
| pPth11-donor | To generate Δ*pth11* | This study |
| pcc-pth11 | | |

**Table S8.** Accession numbers used in this study.

| MSU gene ID[*1] | RAP-DB gene ID[*2] | Gene name |
|---|---|---|
| LOC_Os01g25540 | Os01g0357900 | *OsRALF6* |
| LOC_Os01g25560 | Os01g0358100 | *OsRALF7* |
| LOC_Os02g44940 | Os02g0670500 | *OsRALF8* |
| LOC_Os01g15320 | Os01g0257100 | *OsRALF9* |
| LOC_Os11g26190 | Os11g0448700 | *OsRALF23* |
| LOC_Os10g41980 | Os10g0569400 | *OsRALF26* |
| LOC_Os06g29730 | Os06g0493100 | *OsRALF28* |
| LOC_Os06g18000 | Os06g0288100 | *SOBIR1* |
| LOC_Os09g36320 | Os09g0533600 | *BSR1* |
| LOC_Os12g35610 | Os12g0541300 | *RBOHH* |
| LOC_Os03g12390 | Os03g0225100 | *MKK10-2* |
| LOC_Os02g03410 | Os02g0126400 | *CDPK4* |
| LOC_Os03g57450 | Os03g0788500 | *CDPK10* |
| LOC_Os03g57310 | Os03g0787000 | *SYP121* |
| LOC_Os01g36950 | Os01g0550300 | *NRP1* |
| LOC_Os11g09010 | Os11g0195500 | *PAD4* |
| LOC_Os02g41680 | Os02g0627100 | *PAL1* |
| LOC_Os06g11290 | Os06g0216300 | *OPR1* |
| LOC_Os10g25230 | Os10g0391400 | *TIFY11E* |
| LOC_Os04g48850 | Os04g0578000 | *ACS2* |
| LOC_Os01g61080 | Os01g0826400 | *WRKY24* |
| LOC_Os05g25770 | Os05g0322900 | *WRKY45* |
| LOC_Os01g40260 | Os01g0584900 | *WRKY77* |
| LOC_Os10g33810 | Os10g0478300 | *MYB9* |
| LOC_Os02g41510 | Os02g0624300 | *MYB30* |
| LOC_Os04g23550 | Os04g0301500 | *BHLH6* |
| LOC_Os10g42430 | Os10g0575000 | *BHLH9* |
| LOC_Os01g60020 | Os01g0816100 | *NAC4* |
| LOC_Os11g03300 | Os11g0126900 | *NAC122* |
| LOC_Os12g03040 | Os12g0123700 | *NAC131* |

[*1] The MSU Rice Genome Annotation Project database (https://rice.uga.edu/)
[*2] The Rice Annotation Project Database (https://rapdb.dna.naro.go.jp/)
